# A dose-response model for statistical analysis of chemical genetic interactions in CRISPRi screens

**DOI:** 10.1101/2023.08.03.551759

**Authors:** Sanjeevani Choudhery, Michael A. DeJesus, Aarthi Srinivasan, Jeremy Rock, Dirk Schnappinger, Thomas R. Ioerger

## Abstract

An important application of CRISPR interference (CRISPRi) technology is for identifying chemical-genetic interactions (CGIs). Discovery of genes that interact with exposure to antibiotics can yield insights to drug targets and mechanisms of action or resistance. The objective is to identify CRISPRi mutants whose relative abundance is suppressed (or enriched) in the presence of a drug when the target protein is depleted, reflecting synergistic behavior. Different sgRNAs for a given target can induce a wide range of protein depletion and differential effects on growth rate. The effect of sgRNA strength can be partially predicted based on sequence features. However, the actual growth phenotype depends on the sensitivity of cells to depletion of the target protein. For essential genes, sgRNA efficiency can be empirically measured by quantifying effects on growth rate. We observe that the most efficient sgRNAs are not always optimal for detecting synergies with drugs. sgRNA efficiency interacts in a non-linear way with drug sensitivity, producing an effect where the concentration-dependence is maximized for sgRNAs of intermediate strength (and less so for sgRNAs that induce too much or too little target depletion). To capture this interaction, we propose a novel statistical method called CRISPRi-DR (for Dose-Response model) that incorporates both sgRNA efficiencies and drug concentrations in a modified dose-response equation. We use CRISPRi-DR to re-analyze data from a recent CGI experiment in *Mycobacterium tuberculosis* to identify genes that interact with antibiotics. This approach can be generalized to non-CGI datasets, which we show via an CRISPRi dataset for *E. coli* growth on different carbon sources. The performance is competitive with the best of several related analytical methods. However, for noisier datasets, some of these methods generate far more significant interactions, likely including many false positives, whereas CRISPRi-DR maintains higher precision, which we observed in both empirical and simulated data.

**Author Summary:** CRISPRi technology is revolutionizing research in various areas of the life sciences, including microbiology, affording the ability to partially deplete the expression of target proteins in a specific and controlled way. Among the applications of CRISPRi, it can be used to construct large (even genome-wide) libraries of knock-down mutants for profiling antibacterial inhibitors and identifying chemical-genetic interactions (CGIs), which can yield insights on drug targets and mechanisms of action and resistance. The data generated by these experiments (i.e., sgRNA counts from high throughput sequencing) is voluminous and subject to various sources of noise. The goal of statistical analysis of such data is to identify significant CGIs, which are genes whose depletion sensitizes cells to an inhibitor. In this paper, we show how to incorporate both sgRNA efficiency and drug concentration simultaneously in a model (CRISPRi-DR) based on an extension of the classic dose-response (Hill) equation in enzymology. This model has advantages over other analytical methods for CRISPRi, which we show using empirical and simulated data.

## Introduction

CRISPR technology is becoming an increasingly important tool for genome-wide identification of gene functions in various environmental conditions [1–3]. For example, several different approaches have been devised to exploit CRISPR to induce depletion of target proteins. In the earlier CRISPRko approaches, a nuclease-active form of CAS9 was used to deactivate target genes by cutting the DNA at a target locus and induce DNA repair, which could introduce indels causing frameshifts or inserting novel elements, abrogating their function completely [1–3]. Another approach, CRISPRa, utilizes dCAS9 fusions with effectors that actively enhance or suppress transcription through direct interaction with the RNA polymerase (such as transcription factors that can activate transcription) [4].

In CRISPR interference (CRISPRi), a catalytically-dead CAS9 protein (dCAS9) is recruited to a chromosomal locus by a single guide RNA (sgRNA) with a short (∼20 bp) complimentary sequence and physically blocks transcription [5]. dCAS9 nucleases from several different organisms are available for CRISPRi (e.g. *S. pyogenes, S. thermophilus,* [6]) and different promoters and chemicals have been used for dCAS9 induction. The degree of CRISPR interference can be tuned by modulating the level of dCAS9 expression [7], varying the sgRNA sequence with respect to its length, GC-content, targeting sequence complementarity, position in the gene, or similarity of targeted PAM (protospacer adjacent motif) sequence, to consensus for optimal dCAS9 recognition, [5, 6, 8–11]. While in mammalian systems, efficiency of sgRNAs can vary among multiple cell types, [9], for simplicity, our focus is on studying single defined lineages, as in bacterial strains. Tuning CRISPRi allows to deplete the targeted gene product to intermediate levels [5], which allowed the introduction of the concept of gene ‘vulnerability’ as describing the sensitivity of cells to partial depletion of individual proteins [12]. By this definition, highly vulnerable genes are genes for which even small depletion of the encoded protein causes growth impairment, which can be quantified efficiently on a genome-wide scale using high-throughput sequencing [12]. The vulnerability of a gene can be both condition dependent and strain or cell type dependent [12].

One interesting application of CRISPRi is to reveal targets of antibiotics or mechanisms of resistance through chemical-genetic interactions (CGI) [7, 13]. CRISPRi libraries can be designed to contain multiple sgRNAs targeting each gene, resulting in a set of thousands of individual depletion mutants [12]. In this context, ‘mutant’ refers to a cell line transformed with a integrative plasmid capable of expressing the dCAS9 protein and the unique targeting sgRNA, even though it contains the wild-type gene sequence. The abundance of each mutant can be quantified by amplifying the sgRNA targeting sequence which functions as a molecular barcode, and then performing deep sequencing to count the number of barcodes for each sgRNA in a treatment [6]. The analysis of such datasets is challenging, due to various sources of noise which introduce variability in the counts.

There are several previously published methods for statistical analysis of CRISPR datasets. One, called MAGeCK [14] (originally intended for CRISPRko screens), calculates a log-fold-change (of mean counts) for each sgRNA between a treatment condition and a reference condition (control), and uses a Gaussian distribution to estimate the significance of differences in mean sgRNA abundance between treatments and controls (based on the implementation in the source code, which differs from the description in the publication). To evaluate effects at the gene level, individual sgRNAs are combined in MAGeCK using Robust Rank Aggregation (RRA) to prioritize genes whose sgRNAs show greater enrichment or depletion on average than other genes in the genome. MAGeCK has been used for evaluating chemical-genetic interactions (CGI) with antibiotics [14]. A variant called MAGeCK-MLE [15] fits a Bayesian model by Maximum Likelihood that captures changes in mean counts with increasing time or concentration, along with effectiveness of each sgRNA through posterior probabilities of a binary variable, to determine the overall probability that a gene interacts. Other approaches such as CRISPhieRmix [16] use mixture models to separate effective from ineffective sgRNAs, and thereby identify interacting genes as those containing a significant subset of effective sgRNAs. DrugZ [17] identifies significant interactions by averaging together Z-scores (assuming a Normal distribution) of log-fold-changes of sgRNAs at the gene level. DEBRA [18] utilizes DeSeq, a method for transcriptomic analysis, which employs the Negative Binomial distribution for counts and a more sophisticated method for modeling variance and using it to discriminate genes displaying significant changes in mean counts.

However, most of these methods have one of two limitations when applied to identify genes affecting drug potency. First, CGI experiments are ideally carried out with multiple drug concentrations around the MIC (minimum-inhibitory concentration), since it is often difficult to anticipate what concentration will stimulate the right amount of growth inhibition in combination with CRISPRi-induced depletion of target proteins. However, many of the existing methods analyze the data for each drug concentration independently (i.e. comparing each concentration to a no-drug control). Since knock-down mutants might exhibit depletion at one concentration but not others, results from multiple concentrations must be combined post-hoc. As an example, the authors in [13] chose to combine results from analyzing different concentrations of a given drug using MAGeCK-RRA by taking the union of significant interacting genes at each individual concentration. Due to the noise in these CRISPRi experiments, analyzing concentrations independently increases the risk of detecting false positives (in the sense that non-interacting genes might be spuriously called as hits at different concentrations).

Second, many of the analytical methods do not explicitly take into account differences in sgRNA efficiency (i.e. take sgRNA efficiencies as an input in the model). Different sgRNAs can induce different degrees of depletion of their target genes, and this in turn causes different effects on growth rate, depending on sensitivity of the cells to protein depletion [10]. This can be quantified beforehand by evaluating the growth rate of individual CRISPRi mutants (with unique sgRNAs) in a growth experiment and determining the actual fitness defect caused by target knockdown [11, 12]. In highly vulnerable genes, the effect of protein depletion by sgRNAs on cell growth rate (efficiency) can span a range from no effect to severe growth defect. Early applications of CRISPR were primarily being used to fully inactivate genes (e.g. CRISPRko), rather than to produce graded depletion effects. Therefore, at the time some of these methods were developed, this information was often not used, as methods to quantify sgRNA efficiencies were not well developed. Even in MAGeCK, the Robust Rank Aggregation method treats all sgRNAs in a gene as “equal” a priori, without differentiating them based on the expected effects due to sgRNA efficiency. (Efficiency is not an input.) In contrast, it has been recognized that different sgRNAs can have different efficiency, and several papers have investigated the factors that are associated with stronger sgRNAs [19], especially sequence-based attributes such as similarity to optimal PAM sequence, length and GC content of targeting sequence, mismatches, etc. [5, 8, 10]. Mathis, Otto and Reynolds (11) exploit this to synthetically create a diverse set of sgRNAs with a range of efficiencies by mutating the guide RNA sequences, which they quantify by empirically fitting growth curves for each modified sgRNA with a logistic equation. Interacting genes are then found using differences in the fitted parameters that includes the quantified growth rates and the Hill coefficient. Among all the existing CRISPR analytical methods, MAGeCK-MLE [15] is the only other method that explicitly includes sgRNA efficiencies as an input, which are used to set the prior probabilities that each sgRNA is effective or not (because of their focus on CRISPRko) in the joint probability formula, to initialize for the Expectation Maximization iterations.

In the application to CGI data, a regression model can be used to integrate data over multiple drug concentrations [20]. The degree of a gene-drug interaction is reflected by the coefficient (or slope) for the dependence of CRISPRi mutant abundance on drug concentration. This regression approach was previously introduced in CGA-LMM for analysis of hypomorph libraries (where there is typically just one mutant representing each gene) [20]. It was based on the theory that depletion of the target of a drug should ideally synergize with increasing concentrations of the drug. While exposure to an inhibitory compound will challenge the growth of all the mutants in a hypomorph library, mutants with depletion of a gene that interacts with a drug (e.g. prototypically, an essential gene that is the drug target) will exhibit excess depletion relative to others in the library due to the combined effect of both the growth-inhibition due to the drug treatment in conjunction with the growth-impairment due to knock-down of an vulnerable gene, making these hypomorphic mutants even more sensitive to the drug. For genes that genuinely interact with a given drug, this depletion effect should be exacerbated at higher drug concentrations (i.e. be dose-dependent); thus, genes of greatest relevance would be those that exhibit concentration-dependent effects. While the (log of) abundance of a depletion mutant does not have to decrease perfectly linearly with the (log of) drug concentration to obtain a significant negative coefficient (slope) in the regression, there should be a general trend supporting that relative abundance decreases as concentration increases.

One of the challenges in extending this prior regression approach (CGA-LMM) to CRISPRi screens was incorporating information on sgRNA efficiency. Even in essential genes, some sgRNAs may produce strong depletion of the target, while others might be almost completely ineffective. While sgRNA strength can be partially predicted (with intermediate accuracy) from sequence alone [9, 12], the actual growth phenotype depends on vulnerability of the target gene (sensitivity of cells to depletion of the protein product), which is what is meant by sgRNA efficiency. Even sgRNAs that are predicted to be strong might not cause a growth defect if they are in a non-essential gene. sgRNA efficiency must be empirically quantified by measuring growth rates in standard growth media (e.g. by fitting exponential growth curves based on optical density, or using a reporter gene) with versus without induction of dCAS9, and then calculating relative fitness defects [11]. An alternative approach is to fit the abundance of depletion mutants to a piecewise linear model that allows for a preliminary lag phase, and then extrapolating the model to predicted log-fold-change (LFC) at a fixed number of generations [12]. Any such measure of sgRNA efficiency can be incorporated as a term in the CRISPRi-DR model we present below. Although one could contemplate adding the efficiency of each sgRNA into a simple regression model to predict abundances for each gene, a significant problem (expanded upon below) is that sgRNAs of different efficiency can show different concentration dependence, resulting in non-linear interactions among variables.

In this paper, we propose a modified regression approach for CRISRPi data (called CRISPRi-DR) that incorporates both drug concentration and sgRNA efficiency. The approach is based on the classic dose-response (DR) model for inhibition activity of drugs; the activity of a target protein typically transitions from high to low in the shape of an S-curve as concentration increases (on a log scale), which can be modeled with a Hill equation. The parameters of the Hill equation for a given drug can be fit by performing a log-sigmoid transformation of the mutant abundance data and then using ordinary least-squares regression. We show how sgRNA efficiency can be incorporated into this model as a multiplicative term in the Hill equation, which becomes an additive effect in the log-sigmoid transformed data. The benefit of this model is that it decouples the concentration-dependence from the sgRNA efficiency, so they can be fit as independent (non-interacting) terms in the regression, which ultimately amplifies effects that may be apparent only for a subset of sgRNAs in an optimal efficiency range.

CRISPRi-DR is applicable to libraries where there are multiple sgRNAs representing each gene with a range of efficiencies, which can be quantified empirically as an effect on growth rate (fitness defect). The diversity of efficiencies is useful for identifying synergistic effects with treatments/conditions. Thus, the main requirements for CRISPRi-DR are that: a) there are multiple sgRNAs for each target in the library, b) the sgRNAs vary in predicted strength, and c) the actual efficiencies of the sgRNAs (i.e. growth defects due to target depletion) have been experimentally quantified in control conditions, as an input to the analysis method. The primary use case we focus on is identification of chemical-genetic interactions, with drug concentration as a covariate. We demonstrate the value of the CRISPRi-DR analysis method by re-analyzing the data from a recent paper using CRISPRi for chemical-genetic interactions to identify targets of antibiotics in *M. tuberculosis*. However, the approach can be generalized to analyze experiments with other covariates, such as time-points of a treatment, where there is a sigmoidal response in growth. We illustrate this by using CRISPRi-DR to analyze an *E. coli* CRISPRi dataset from an experiment to determine genes differentially required for growth on different carbon sources [11].

## Methods

The CRISPRi-DR method applies to CRISPRi experiments that involve using high-throughput sequencing to tabulate sgRNA counts representing abundance of individual CRISPRi mutants in a population (pooled culture). Each mutant has an sgRNA (on a plasmid) mapping to a target gene that can reduce its expression (e.g. with dCAS9 induction). In CGI applications, the culture is treated with antibiotics or inhibitors at various concentrations, along with a no-drug control, and DNA is extracted, PCR-amplified, and sequenced, producing counts representing each sgRNA. If 𝑌_𝑖j𝑘_ is the abundance (i.e. count) for an sgRNA 𝑖 in a condition 𝑗 for replicate 𝑘, normalized abundance can be calculated by 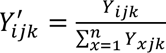 , where each count is divided by the sum of counts of all the sgRNAs observed in a given condition and replicate. Let 𝑈′_𝑖_be the normalized abundance of sgRNA 𝑖 in the uninduced condition, then the normalized relative abundances of an sgRNA 𝑖 in all induced samples can be calculated as: 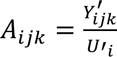 , assuming that the counts in the uninduced condition represents full abundance of each clone (normal growth without target depletion).

### CRISPRi Dose-Response Model

The CRISPRi-DR model for analyzing CRISPRi data from CGI experiments is an extension of the basic dose-response model, extended to incorporate sgRNA efficiencies. The dose-response effect of an inhibitor on the activity of an enzyme is traditionally modeled with the Hill-Langmuir equation.

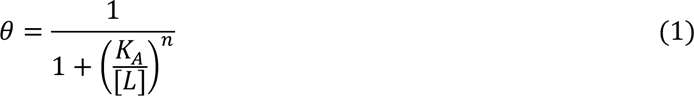

where 𝜃 is the fraction of abundance (relative to no drug), [L] is the ligand concentration, K_A_ is the concentration at which there is 50% activity and 𝑛 is the Hill coefficient.

Applying Eq (1) to the CGI data, the relative abundance of sgRNAs 𝐴_𝑖j𝑘_ is used as the predictor variable and [D_j_] is the concentration of drug *j* that the *k*th replicate count of sgRNA *i* was extracted from,

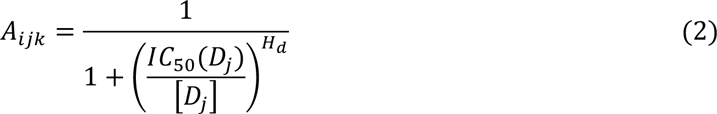

The unknown parameters are the IC_50_ value (inhibitory concentration that causes 50% growth inhibition) and the Hill coefficient 𝐻_𝑑_. The plot of the concentration versus relative abundance of an sgRNA (𝐴_𝑖j𝑘_) produces a sigmoidal curve, demonstrating how activity decreases as concentration increases, with the IC_50_, representing the mid-point of the transition.

The dose-response model seen in Eq 2 can be extended to account for sgRNA efficiency by incorporating a multiplicative factor in the denominator:

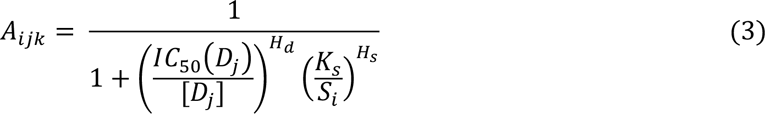

sgRNA efficiency, *S_i_*, is an empirical measure of the degree of growth impairment resulting from target depletion. This can be assessed in several ways, such as estimating change in exponential growth rate in a reference condition in a growth experiment [21]. Alternatively, Bosch et al [12] use estimated log-fold change of abundance (induced vs uninduced) at a fixed number of generations of growth in-vitro in the absence of drug, extrapolated from a model fit to empirical data (passaging experiment) that allows for a lag phase. 𝐾_𝑠_ represents the unknown intermediate sgRNA efficiency that causes 50% depletion of mutant abundance (half-way between no depletion and full depletion), and the 𝐻_𝑠_ is the unknown Hill coefficient that represents how sensitive mutant abundance is to depletion of the target protein.

### Relationship between drug concentration and gene depletion within the CRISPRi-DR model

Abundance of mutants in a CRISPRi CGI experiment can be affected simultaneously by both presence of an inhibitor and depletion of an interacting gene. However, the concentration-dependent effect of a drug on mutant abundance can be different for sgRNAs of different efficiency. Fig 1 illustrates the interaction between these two effects for *rpoB* (RNA polymerase beta chain) in an *Mtb* CRISPRi library treated with rifampicin with 5 days pre-depletion. The lines in Fig 1A are regression fits obtained for each sgRNA in *rpoB* using regression of log abundances against log concentration of rifampicin, log(𝐴_𝑖j𝑘_) = 𝐶 + 𝐵 · log([𝐷_j_]) , where *C* is in the intercept and *B* is the slope of the regression, representing concentration dependence, and log(𝐴_𝑖j𝑘_) are log relative abundances obtained as described above. The left-most side of Fig 1A shows the range of abundances in the no-drug control (induced library in media without rifampicin). These differences in abundances (dispersion along Y-axis) are due solely to the growth impairment caused by depleting RpoB. As concentration of RIF increases, some of the sgRNAs show very negative slopes, while other sgRNAs show slopes closer to 0. A parabolic-type curve emerges in Fig 1B when the slopes 𝐵 from the regressions are plotted against the sgRNA efficiencies. Both the most efficient sgRNAs (colored red) and the least efficient sgRNAs (purple) have slopes around 0 (no concentration dependence). Highly efficient sgRNAs (red) can cause excessive depletion (even without drug), making it difficult to detect additional decreases due to increasing drug concentration. Comparatively, sgRNAs with very low efficiency (purple) might not induce enough depletion to synergize with the drug. The sgRNAs surrounding the minimum point of the parabolic curve (dashed line) in Fig 1B reflect those of intermediate efficiency where the ability to detect synergy with the drug is maximized. These are the sgRNAs in Fig 1B that show the most negative slope with increasing concentration (dark green-indigo). As Fig 1C shows, the efficiency where the slopes reach their extremes (most negative; or most positive for those showing enrichment) can be different for each gene but tend to fall in an intermediate region of sgRNA efficiency (0 to −5). The histogram shows that sgRNA efficiency at which the most extreme (largest or smallest) concentration-dependent slope is achieved over all interacting genes (236 for RIF D5). Hence, the sgRNAs that are optimal for detecting CGIs are not necessarily the strongest (most efficient). The variability of concentration-dependence (slope) with sgRNA efficiency suggests a possible non-linear interaction between the variables. This nonlinearity is captured in the multiplicative terms of the dose-response model (Eq (3)).

**Fig 1.**
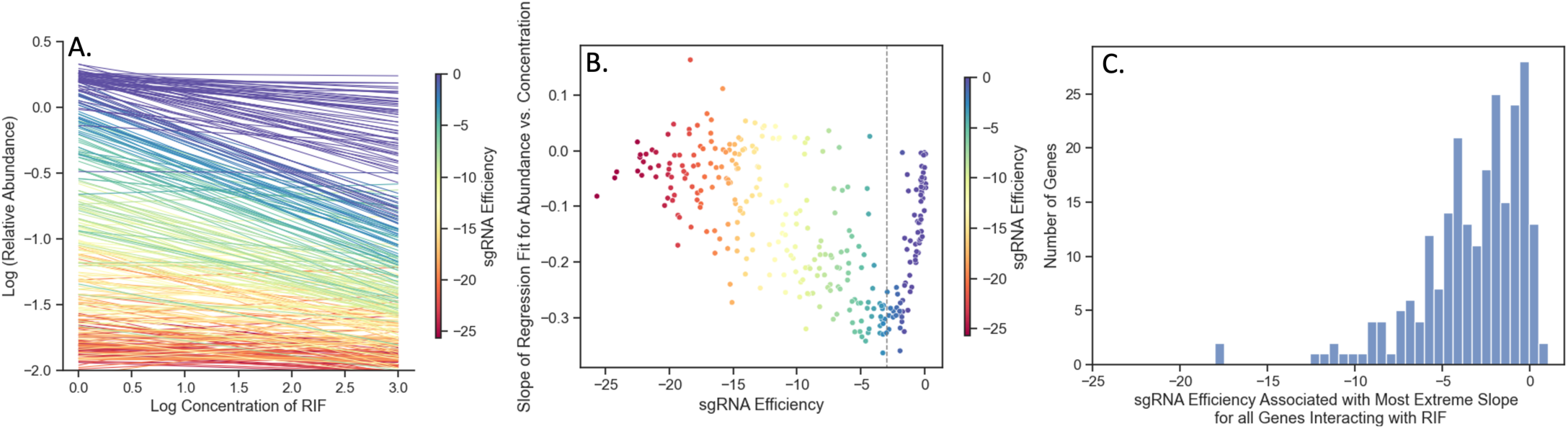
Effect of sgRNA efficiency on concentration dependence for sgRNAs in *rpoB* in a CRISPRi library treated with RIF (D5). (A) Regression lines for log(relative abundance) against log(concentration) for all sgRNAs in *rpoB* in a library treated with RIF for 5 days pre-depletion. The lines that reflect the extremes of the sgRNA efficiency (red or purple), are flat and do not show much change in abundance. Comparatively, intermediate sgRNA efficiency (dark green to indigo) shows the most negative slopes, reflecting maximum synergy with drug. (B) Comparison of sgRNA efficiency and slopes of the regressions seen in Panel A for each sgRNA. Each point is an sgRNA colored by its efficiency. The most efficient sgRNAs (purple) and the least efficient sgRNAs (red) show concentration slopes around 0. The dotted line reflects the minimum of the parabolic curve. (C) Histogram of sgRNA efficiencies where the slopes reach their most extreme (positive or negative) for 236 interacting genes in RIF D5. The distribution shows that most of the extrema sgRNAs for interacting genes fall in the range of −5 to 0 (note: not the strongest sgRNAs, which would have efficiencies around −25).

### Linearization and parameter estimation

The dose-response model Eq (3) can be linearized through a log-sigmoid transformation.

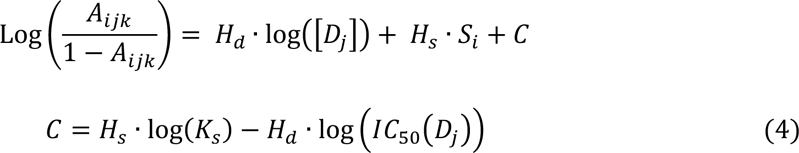

In this log-sigmoid transformed space, the concentration-dependence and effect of sgRNA efficiency have been decoupled, appearing as independent linear terms with the Hill coefficients (𝐻_𝑠_ and 𝐻_𝑑_) as the variables to solve for by a standard regression. The inflection parameters of the sigmoid curve (𝐾_𝑠_and IC_50_) are combined in the intercept *C* in the model. Importantly, this model implies that the effects of growth impairment due to the depletion of a vulnerable gene and growth inhibition due to the drug on the overall (relative) abundance of a given mutant become additive in this log-sigmoid-transformed space. To illustrate this, the CRISPRi-DR equation is simulated by plotting idealized relative abundances (in Fig 2) using parameters chosen to emulate what is seen in Fig 1A, the plot of slopes over a systematic range of sgRNA efficiencies and drug concentrations for *rpoB*. In Fig 2A, the slopes of the concentrations are plotted against abundances calculated using the dose-response model. The slopes vary as a function of the starting depletion (left-hand side), which is due to sgRNA efficiency alone (colored gradient based on sgRNA efficiency value). The slopes are most negative for intermediate sgRNA efficiency, colored with a dark blue-green hue representing sgRNA efficiency around −10. Fig 2B illustrates the result of the linearization (log-sigmoid transformation) of the Hill equation. All the individual sgRNA regression lines over concentration become parallel, eliminating the dependence on sgRNA efficiency, and allowing them to be fit by a single common slope representing the concentration-dependence averaged over all the sgRNAs.

**Fig 2.**
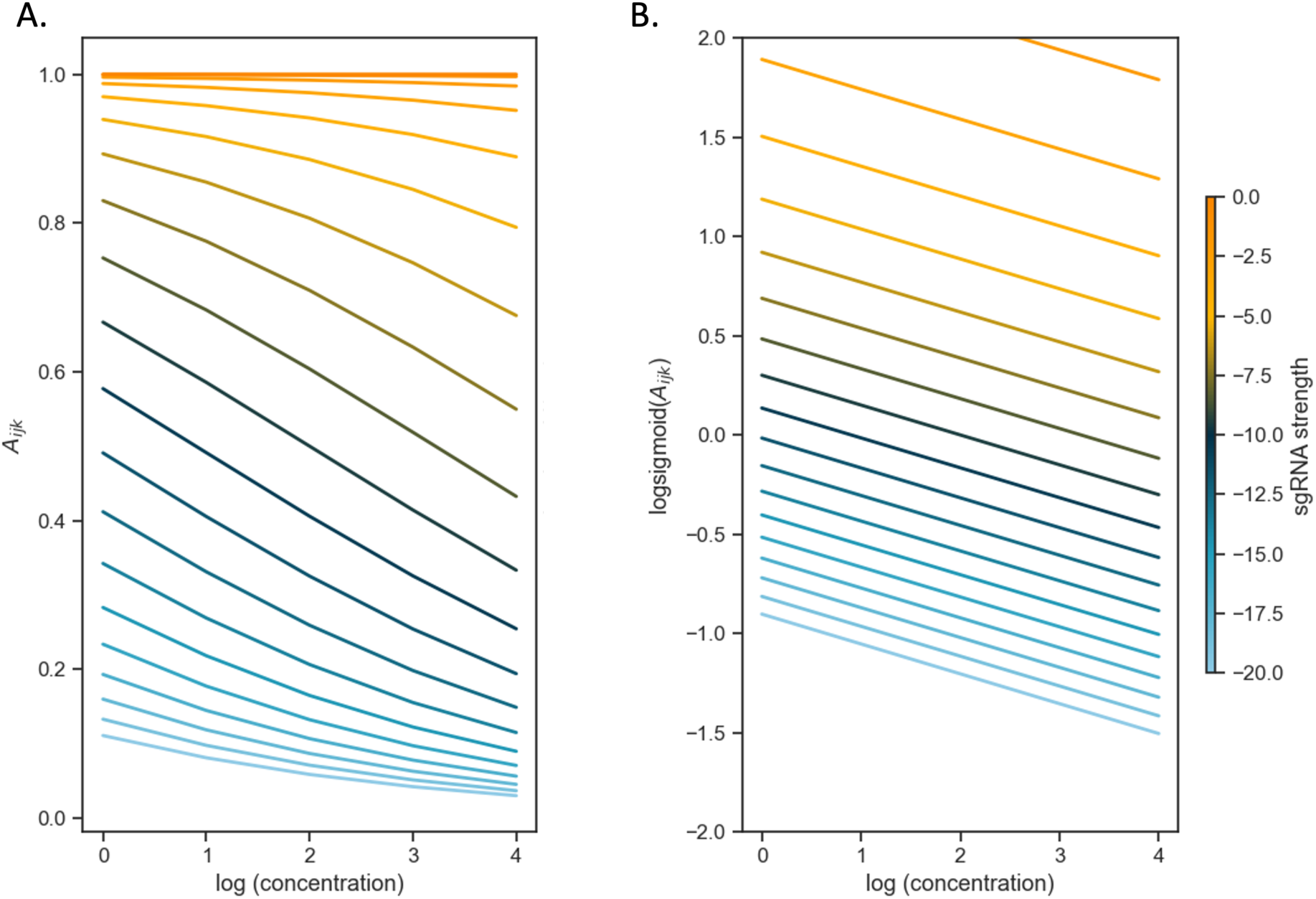
The log-sigmoid transformation of abundances allows the CRISPRi-DR model to factor in the non-linear effect of sgRNA strength on concentration dependence. (A) Simulation of sgRNAs abundances for an ideal essential gene. Parameters used in simulation: H_s_ = −4, IC_50_ = 0.25, K_s_ = −10 and H_d_ = −0.5 over a range of sgRNA efficiencies and drug concentrations. (B) When the log-sigmoid transformation of the abundances is applied, we see all the regression fits are parallel to one another, allowing to be fit by a single common slope, representing the concentration dependence over all sgRNAs, regardless of sgRNA efficiency.

Experimental data (i.e. counts from sequencing, converted to relative abundances for mutants with each sgRNA) are fit on a gene-by-gene basis using ordinary least-square (OLS) regression by the following formula:

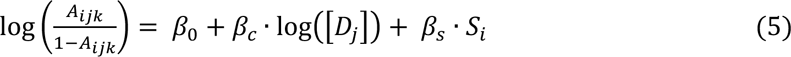

where *A* (relative abundance for each CRISPRi mutant at given drug concentration), *S_i_* (sgRNA efficiency) and *[D_j_]* (concentration of drugs) are columns of a melted matrix. To include the control samples (no-drug, dCAS9-induced controls) in the regression, they are treated as one two-fold dilution lower than the lowest available concentration tested for the drug (to avoid taking the log of 0). Since the log-sigmoid transform of the relative abundances is taken, they must be within the range of (0,1). Although relative abundances greater than 1.0 are possible in treated conditions (relative to uninduced, no-drug controls), especially in cases where target depletion confers a growth advantage and consequent enrichment, we use a squashing function to ensure the relative abundances range between 0 and 1, which is required to take the log-sigmoid transform.

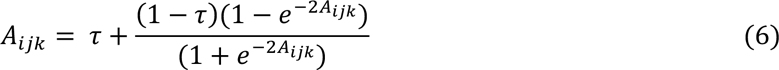

where 𝜏=0.01 is a pseudo count needed to make abundances non-zero for taking logarithms. Relative abundances that are greater than 1.0 are mapped to just below 1.0, though the mapping is monotonic, so the order among sgRNAs is still preserved (higher abundances become exponentially closer to 1.0).

### Significance Testing

The statistic that indicates the degree of interaction of each gene with a given drug is the coefficient for the 𝑙𝑜𝑔([𝐷]) term (i.e. slope) in the model. To determine whether the interaction is statistically significant, a Wald test [22] is applied to calculate a P-value reflecting whether the coefficient is significantly different than 0, adjusting for a target FDR (false discovery rate) of 5% over the whole genome using the Benjamini-Hochberg procedure [23]. However, the Wald test by itself yields many genes predicted to interact with the drug (often thousands) with adjusted P-value < 0.05. The test selects genes with slopes that are technically different than 0, but not necessarily large enough to be relevant to the drug mechanism. Our assumption is that most of genes in the genome do not interact with a given drug (at least not directly involved in the mechanism of action or resistance). Many genes have small positive and negative slopes, possibly due to some source of noise in the experiment or generalized phenotypic interactions, which should be filtered out. Therefore, genes are filtered based on the magnitude of the slopes (analogous to the requirement of |LFC|>1 used by Li, Poulton (13) to filter significant genes by MAGeCK). The distribution of slopes over all genes is assumed to be a Normal distribution, and Z-scores are computed for every gene 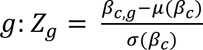, where 𝜎(𝛽_𝑐_) is the standard deviation of the slopes of log concentration dependence and 𝜇(𝛽_𝑐_) is the mean of the slopes. Genes with |𝑍_g_| < 2.0 are filtered out. This produces hits whose slopes are significant outliers (>2𝜎) from the rest of the population (i.e. genes in the genome). There are two groups of hits, corresponding to the two tails of the distribution: enriched hits where 𝑍_g_ > 2.0, and depleted hits, 𝑍_g_< −2.0.

## Results

### CRISPRi Dataset and Pre-processing

A chemical-genomics dataset was obtained from high-throughput sequencing of a CRISPRi library of *M. tuberculosis* (*Mtb*) that had been treated with several antibiotics. The library consists of 96,700 sgRNAs targeting all 4019 genes in the *Mtb* H37Rv genome [13]. This library was intentionally constructed to focus on probing essential genes (based on prior TnSeq analysis [24]), with a mean of 83 sgRNAs per essential gene, but there are some sgRNAs in each non-essential gene too (mean of 10 sgRNAs per non-essential gene).

The library was individually treated with 9 anti-TB drugs (rifampicin, RIF; isoniazid, INH, ethambutol, EMB; vancomycin, VAN; levofloxacin, LEVO; linezolid, LZD; streptomycin, STR; clarithromycin, CLR; bedaquiline, BDQ) to evaluate and validate the CRISPRi system in preparation for target identification for novel inhibitors (from high-throughput screens). These drugs were selected because certain genes are expected to interact for each (based on known mechanisms of action), although additional genes might also exhibit interactions, which could extend our knowledge. We note that some drug targets are members of a complexes; although a drug may bind directly to one subunit, other subunits in those complexes often show similar CRISPRi phenotypes. RIF binds RpoB (RNA polymerase subunit) inhibiting transcription and compensatory mutations are often found in *rpoC* [25], BDQ binds and inhibits AtpE (subunit of the ATP synthase) [26] and *mmpL5* effluxes the drug [27], GyrA and GyrB (subunits of DNA gyrase) would be expected to interact with fluoroquinolones like LEVO [28], EMB targets *embABC* in the arabinogalactan pathway [29, 30], CLR, LZD and STR bind to the ribosome and inhibit translation, which can be protected by rRNA methyltransferases [31–33], VAN binds to peptidoglycan and is expected to interact with genes in the peptidoglycan synthesis pathways [34, 35], and genes such as *inhA, katG, ahpC, ndh, mshA* and *cinA* are implicated in the mechanism of action or resistance for isoniazid, an inhibitor of mycolic acid synthesis [36–39]. These define selected interactions that would be expected to be observed in a CRISPRi CGI experiment.

Samples of the library (pooled cultures) were treated with each of the drugs, with induction of the Sth1 dCAS9 by ATC (anhydrotetracycline), and were sequenced in triplicate at several concentrations for each drug at 2-fold dilutions around the MIC, along with control samples representing the no-drug samples (0 concentration). Three periods of pre-depletion were evaluated: 1, 5, and 10 days (D1, D5, and D10), since it was initially unknown how many days would be optimal for reducing protein expression after induction of CRISPRi. The measurements reported in this experiment are observed counts of sgRNAs, representing the relative proportion of each mutant in the population (pooled culture of CRISPRi mutants). Abundance of a mutant increases or decreases if silencing of the targeted gene causes a change in fitness. Although target proteins are knocked down by inhibiting transcription via CRISPRi, intracellular protein levels are not directly measured in the experiment. Instead, unique nucleotide barcodes representing each sgRNA are amplified from (integrated) plasmids in the cells, sequenced, and counted. The counts reflect the relative abundance of each CRISPRi mutant. Samples were normalized by dividing individual counts for each sgRNA by the sample total (sum over all sgRNAs).

In this dataset, prior estimates of sgRNA efficiency were obtained from empirical data by fitting a piecewise-linear equation to fitness over multiple generations, and then using the model for to extrapolate the predicted log-fold change (LFC) each sgRNA at 25 generations [12]. The scale for these efficiencies ranged between −25 (highest depletion) and 0 (no depletion). To determine the effect of depletion solely due to the sgRNA (without drug), uninduced samples (in the absence of dCAS9 induction, -ATC) were also sequenced, to provide counts representing mutant abundances in the absence of depletion of targets as an input to the model.

### The CRISPRi-DR model accurately predicts sgRNA abundances from sgRNA strength and drug concentration

The CRISPRi-DR model was fitted for all chemical-genetic interaction datasets from Li, Poulton (13) , which included nine drugs tested at three different concentration levels (after 1, 5, and 10-days of pre-depletion without drug). The analyses by CRISPRi-DR found a range of tens to hundreds of significant genes for each dataset. Table 1 show a more detailed account of the significant genes founds in these CRISPRi screens by CRISPRi-DR, categorized into depleted (mutant abundance decreases with drug concentration) and enriched (mutant abundance increases with drug concentration).

**Table 1.** Number of Significant Genes found by CRISPRi-DR across the nine drugs CRISPRi screen for each of pre-depletion days.

| DRUG | D1 |  | D5 |  | D10 |  |
| --- | --- | --- | --- | --- | --- | --- |
|  | Depleted | Enriched | Depleted | Enriched | Depleted | Enriched |
| **BDQ** | 89 | 99 | 121 | 48 | 116 | 37 |
| **CLR** | 182 | 23 | 75 | 89 | 79 | 71 |
| **EMB** | 15 | 160 | 6 | 161 | 51 | 130 |
| **INH** | 33 | 57 | 9 | 93 | 16 | 96 |
| **LEVO** | 80 | 47 | 50 | 50 | 19 | 4 |
| **LZD** | 45 | 123 | 44 | 140 | 54 | 65 |
| **RIF** | 117 | 65 | 165 | 57 | 146 | 53 |
| **STR** | 57 | 90 | 44 | 37 | - | - |
| **VAN** | 193 | 8 | 149 | 26 | 135 | 45 |

The significant genes identified by CRISPRi-DR generally have coefficients of concentration dependence that are outliers with respect to the rest of the genes. Fig 3 shows the distribution of the slopes calculated for genes in a library treated with EMB (one day of pre-depletion, D1). The threshold for this distribution where |𝑍_g_|>2.0 and adjusted P-value < 0.05, is at slope = −0.37 and slope = 0.26 (vertical bars). The 164 total genes in the tails outside the vertical lines are significant genes. These genes include the targets of EMB: *embA, embB* and *embC* [29, 30], which have slopes −0.45, −0.43 and −0.32, respectively.

**Fig 3.**
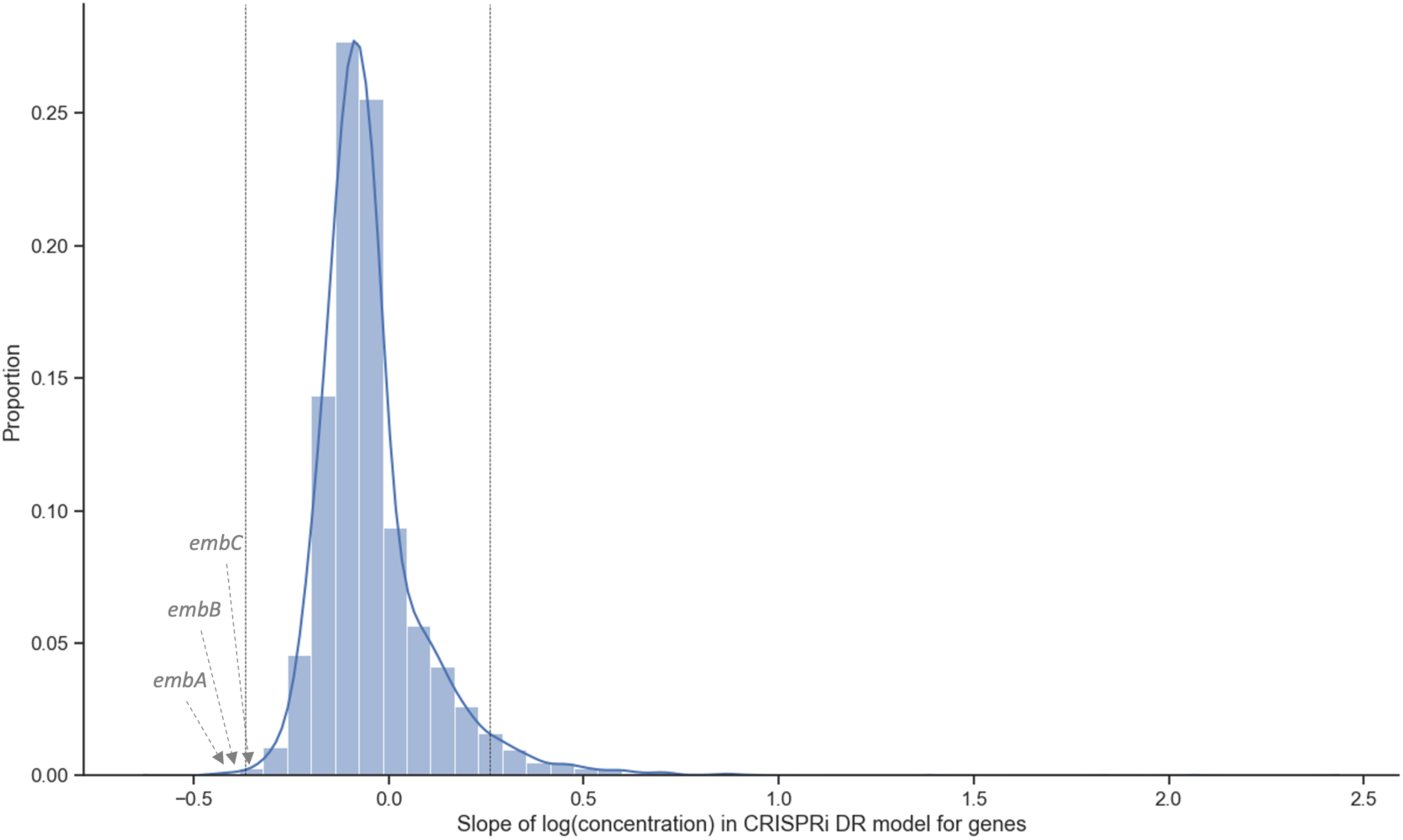
Coefficients of concentration-dependence from CRISPRi-DR model fitted for EMB D1 (1 day of pre-depletion). The distribution of the slopes of concentration dependence, extracted from the model fit for each gene. The vertical lines are at slope = −0.37 and slope = 0.26. These are the slopes adjusted P-value < 0.05 and the |Z-score|> 2.0. 164 genes have significant slope values, i.e., 164 genes show a significant change in abundance with increasing EMB concentration while accounting for sgRNA strength.

To evaluate the relative importance of the sgRNA efficiency and drug concentration features to the CRISPRi-DR model, each gene was fit with two ablated models: M_d_ and M_s_. The M_d_ model contained only log concentration as a predictor: 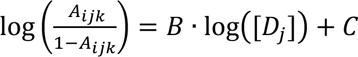 and the M_s_ model only contained sgRNA efficiency as a predictor: 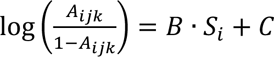. In the EMB D1 experiment, the average *r^2^* (% variance explained) across all genes in full CRISPRi-DR model is 0.43. Comparatively, the average *r^2^* is 0.29 for M_s_ and 0.13 for M_d_. *embA* also appears as one of the genes in the M_d_ set of significant interactors, but the other targets of the drug, *embB* and *embC* do not appear in the sets of significant interactors for either of these ablated models. As a measure of the model quality (goodness of fit), the Akaike Information Criterion (AIC) for the full model in the EMB D1 experiment is 87.6, whereas the AIC of M_d_ is 300.7 and AIC of M_s_ is 124.7. The full model has the lowest AIC, indicating it is the best fitting model of the three. The AIC for the model incorporating only drug concentrations but not sgRNA efficiency (M_d_) is highest (worst), suggesting that sgRNA efficiency encodes critical information needed for predicting mutant abundance. A Likelihood Ratio Test shows that the differences between these models is highly significant (P-value << 0.05; 𝜒^2^distribution using one degree of freedom, since the ablated models each have one parameter less than full model). The *r^2^* values and results of the AIC-based likelihood comparison indicate that sgRNA efficiency contributes strongly to accuracy of the model, and reinforces the importance of including sgRNA efficiency as a term in the CRISPRi-DR model.

The improved performance of CRISPRi-DR over the reduced models for EMB extends to the other drugs tested, as seen in Fig. S1. In all the experiments, the number of genes with fits with *r^2^* > 0.5 is the greatest in the full CRISPRi-DR model, and the number of genes with fits that have *r^2^* > 0.5 is greater in model M_s_ than M_d_. This demonstrates that in all conditions, both concentration and sgRNA strength are needed to make accurate estimates of mutant abundance.

### CRISPRi-DR and MAGeCK have a high concordance of predicted gene-drug interactions

Most of the significant CGIs identified by the CRISPRi-DR model were also identified by MAGeCK (MAGeCK-RRA) as reported in Li, Poulton (13), but MAGeCK often identifies many additional genes that are not detected as significant by the CRISPRi-DR model. Although there are some datasets where MAGeCK and CRISPRi-DR detect about the same number of significant interactions, as shown in Fig 4A and the Extended Figure S2 from Li, Poulton (13), there are quite a few datasets where MAGeCK finds substantially more hits than CRISPRi-DR, such as VAN D1, where MAGeCK finds over 1066 significantly depleted genes (even with the filter of |LFC|>1 applied), whereas CRISPRi-DR finds only 196 significant interactors. As seen in the Venn diagrams in Fig 4B, there is high overlap of calls made by the two methodologies (enriched and depleted combined). Across all the datasets, an average of 62.2% of genes identified as significant by CRISPRi-DR are also found to be significant by MAGeCK. In the depicted datasets in Panel B, nearly all the calls made by CRIPSRi-DR overlap with those made by MAGeCK. However, MAGeCK makes quite a substantial number of calls (significant interacting genes) that are not found by CRISPRi-DR. Additional details of the overlap of significant interacting genes in MAGeCK and CRISPRi-DR can be found in Table S3.

**Fig 4.**
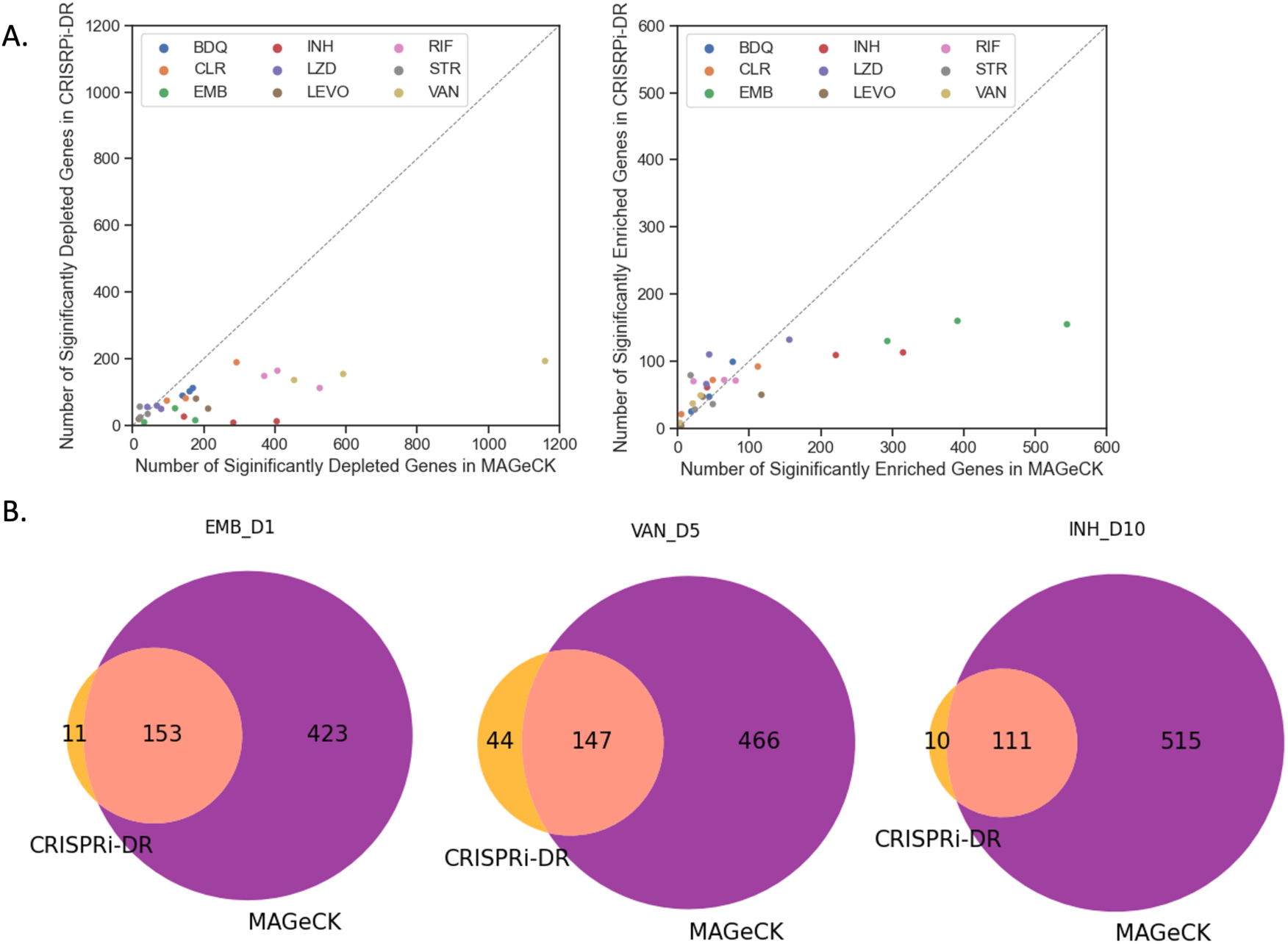
Comparison of significant interactions found by CRISPRi-DR and MAGeCK. (A) The points in the plots are the analyses of CRISPRi screens by both MAGeCK and CRISPRi-DR, colored by drug treatment. The left plot compares the depleted hits called by the two methodologies and the right plot compares the enriched hits called by the two methodologies. The number of hits (both enriched and depleted) are slightly greater in MAGeCK than in the CRISPRi-DR model. (B) Venn Diagram of significant genes, both depleted and enriched, found by CRISPRi-DR and MAGeCK for select drug-treated libraries. The genes identified by CRISPRi-DR are primarily a subset of the hits found by MAGeCK.

### CRISPRi-DR model correctly detects genes known to interact with anti-tubercular drugs

When genes are ordered by coefficients of the slope representing the dependence of abundance on drug concentration from the CRISPRi-DR model, genes known to affect the potency of the anti-mycobacterial drug tested are ranked highly, as expected (Table 2). The more positive a gene’s coefficient is, the higher the gene’s enrichment ranking, and the more negative a gene’s coefficient is, the higher it’s depletion ranking.

**Table 2:** Ranking of Select Genes using the CRISPRi-DR model in 1 Day pre-depletion of treated libraries.

| *Drug* | *Gene* | *D1 Depletion*<br>*Ranking* | *D1 Enrichment*<br>*Ranking* |
| --- | --- | --- | --- |
| *BDQ* | *atpA* | 11 | 4022 |
| *BDQ* | *atpB* | 6 | 4027 |
| *BDQ* | *atpC* | 51 | 3982 |
| *BDQ* | *atpD* | 14 | 4019 |
| *BDQ* | *atpE* | 25 | 4008 |
| *BDQ* | *atpF* | 9 | 4024 |
| *BDQ* | *atpG* | 12 | 4021 |
| *BDQ* | *atpH* | 8 | 4025 |
| *BDQ* | *mmpL5* | 2 | 4031 |
| *CLR* | *RVBD3579c* | 40 | 3993 |
| *CLR* | *erm(37)* | 1 | 4021 |
| *EMB* | *embA* | 2 | 4031 |
| *EMB* | *embB* | 3 | 4030 |
| *EMB* | *embC* | 19 | 4014 |
| *INH* | *inhA* | 3 | 4030 |
| *INH* | *ahpC* | 2 | 4031 |
| *INH* | *cinA* | 5 | 4028 |
| *INH* | *katG* | 4031 | 2 |
| *INH* | *ndh* | 4028 | 5 |
| *INH* | *mshA* | 4025 | 8 |
| *LEVO* | *gyrA* | 4012 | 21 |
| *LEVO* | *gyrB* | 4021 | 12 |
| *LZD* | *erm(37)* | 3865 | 168 |
| *LZD* | *tsnR* | 4032 | 1 |
| *RIF* | *rpoB* | 94 | 3939 |
| *RIF* | *rpoC* | 147 | 3886 |
| *STR* | *RVBD2477c* | 4021 | 12 |
| *STR* | *gidB* | 4022 | 11 |

For each drug, the CRISPRi-DR model is run on each gene (using data from D1). The coefficient for the slope of concentration dependence (𝛽_𝑐_) is extracted from the fitted regression and used to rank the genes both in increasing order (for depletion) and inversely (for enrichment). Green reflects results consistent with expectations based on knowledge of known gene-drug interactions

Genes that encode the target of a drug would typically be expected to have a high depletion rank, i.e., show a negative slope, indicating that as concentration increases, abundance for the given depletion-mutant decreases. This can be seen in S1 Table, in the ranking for genes using the CRISPRi-DR model. These genes rank the highest in D1 and not as well in D10. This can be attributed to the fact that, after 10 days of pre-depletion, these mutants are already quite depleted, even at concentration 0, increasing noise, and making it difficult to pick up on concentration-dependent signals (further depletion). Therefore, the ranking of relevant genes in D1 was assessed in this analysis (Table 2).

For isoniazid (INH), there are multiple relevant genes identified by CRISRPi-DR, including *inhA, ahpC*, *ndh* [40], and *katG* [41]*. inhA* (enoyl-ACP reductase) is an essential gene in mycolic acid pathway that is the target of INH, and AhpC (alkyl hydroperoxide reductase) responds to the oxidative effects of isonicotinic radicals in the cells, MshA is a protein involved in synthesis of mycothiol, which helps maintain redox balance [39], and CinA is a NADH metabolizing protein that can hydrolyze the isoniazid-NAD adduct [38]. Therefore, as dosage of the drug increases, the abundances of the mutants of these genes should decrease. These genes are in the top 10 highest ranked depletion genes for INH (see Table 2). In contrast, *katG* and *ndh* are found among the top 5 enriched hits, exhibiting increased survival when the proteins are depleted. KatG (catalase) is the activator of INH, and the most common mutations in INH-resistant strains occur in *katG*, decreasing activity [42]. *Ndh* (type II NADH reductase) mutants have also been shown to decrease sensitivity to INH by shifting intracellular NADH levels (needed for INH-NADH adduct formation), and mutations in *ndh* have been shown to be defective in target enzyme (NdhII) activity [40], which is consistent with the observation in the CRISPRi data that depletion of *ndh* leads to increase survival in the presence of INH. Similarly, *mshA* is highly enriched, consistent with mutations found in resistant mutants.

For EMB, *embA, embB,* and *embC* (subunits of the arabinosyltransferase, target of ethambutol, EMB) rank within the top 100 depleted genes for all three pre-depletion conditions [29, 30]. However, interactions with the other genes in the arabinogalactan pathway, like *ubiA* (which sometimes acquires resistance mutations [43]), were not observed.

In RIF, *rpoB* and *rpoC*, subunits of the core RNA polymerase, are ranked within the top 150 genes. Significant negative interacting genes for RIF also include many cell wall related genes such as *ponA2, rodA, ripA, aftABCD, embABC,* etc., consistent with recent studies that show RIF exposure (or mutations in *rpoB*) leads to various cell wall phenotypes [44–46]. Similarly, the target of bedaquiline (BDQ), the F0F1 ATP synthase (which includes 8 subunits encoded by *atpA-atpH,* of which AtpE is the one bound by BDQ) [26], and *mmpL5,* which can eflux the drug [27], are ranked within the top 40 depleted genes in BDQ.

The significantly interacting genes in vancomycin (VAN) involve many genes in the cell wall/membrane/envelope biogenesis pathway (as defined by in COG pathways [47]) (adjusted P-value for pathway enrichment = 0.0004 using Fisher’s Exact Test). This follows previous studies that show cell wall genes are targets of vancomycin [48, 49], which binds to peptidoglycan in the cell wall.

In levofloxacin (LEVO), CRISPRi mutants of *gyrA* and *gyrB* (subunits of the DNA gyrase, the target of fluoroquinolones) are also observed to be enriched. The reason that depletion of this drug target leads to enrichment of mutants (hence a growth advantage, rather than the expected growth impairment) is likely due to reduced generation of double-stranded breaks in the DNA and other toxic intermediates as a side-effect of inhibiting the gyrase, an effect that has been observed in *E. coli* [50].

For clarithromycin (CLR), an inhibitor of translation, *Rv3579c* and *erm(37)* are observed as hits. *Erm(37)* adds a methyl group on the A2058/G2099 nucleotide in the 23S component of the ribosome, the same site in which clarithromycin binds [51]. This natively increases tolerance to CLR in *Mtb*. As this gene is depleted, CLR has greater opportunity to bind, reducing the cells’ natural tolerance to the drug. Consistent with this observation, e*rm(37)* has a depletion rank of #1 in the CLR D1 condition. Rv3579c is another methyltransferase with a similar function that ranks highly (#35) in CLR.

In contrast to methylation inhibiting the binding of CLR, there are ribosome methyltransferases in *Mtb,* where methylation facilitates binding of a drug. Mutants for these genes would be expected to show a high enrichment rank in presence of drug. For instance, streptomycin (STR) interferes with ribosomal peptide/protein synthesis by binding near the interaction of the large and small subunits of the ribosome [52]. Two relevant genes that influence the binding of STR include *gidB* and *Rv2477c/ettA*. GidB is an rRNA methyltransferase that methylates the ribosome at nucleotide G518 of the 16S rRNA, the position at which STR interacts [33], increasing native affinity for STR. This is consistent with the observation that one of the most common mutations in STR-resistant clinical isolates is loss of function mutations in *gidB* [53]. Rv2477c is a ribosome accessory factor, also known as EttA, which is an ATPase that enhances translation efficiency. It has also recently been shown to bind the ribosome near the P-site (peptidyl transfer center), potentially interfering with binding of aminoglycosides [54], and loss-of-function mutations observed in drug-resistant clinical isolates of *M. tuberculosis* have shown to confer resistance to STR [13]. The ranking of both genes using the CRISPRi-DR model are within the top 12 enriched genes in STR. For linezolid (LZD), relevant genes identified are *erm(37)* and *tsnR.* TsnR is an rRNA methyltransferase, analogous to GidB, and results in tolerance to LZD in a similar manner as GidB does for STR [13]. Following this expectation, *tsnR* has an enrichment ranking of #1 in LZD. Whereas depletion of Erm(37) gives tolerance to CLR, it increases sensitivity to LZD. The nucleotides that Erm(37) methylates in the 23S RNA are proximal in 3D space to where mutations conferring LZD-resistance are found, which both lie in the PTC (peptidyl-transfer center) of the ribosome [55].

### The CRISPRi-DR model is less sensitive to noise than MAGeCK

A reason that the CRISPRi-DR model shows lower consistency with MAGeCK (RRA) in some datasets could be due to different sensitivity to noise. There is some noise in these experiments due to variability in sequencing sgRNA counts across multiple concentrations and replicates. This can differentially affect the accuracy of predictions of gene-drug interaction made by these models. Three replicate counts were collected for estimating the relative abundance of each CRISPRi mutant (with a unique sgRNA) in the presence of a drug at a given concentration. The coefficient of variation (CV) can be used to measure the relative consistency of measurements across these observations, which in turn can be used to evaluate the sensitivities of CRISPRi-DR and MAGeCK to noise in the raw data.

For each sgRNA *s_i_* the coefficient of variation (CV) was calculated across the relative abundances for the 3 replicates for each concentration ( C ) in drug (D) 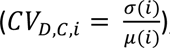, where 𝜎(𝑖) is the standard deviation of the 3 relative abundances in concentration C and 𝜇(𝑖) is the mean. In Fig 5A, the 𝐶𝑉_𝐷=𝐷𝑀𝑆@,𝐶=0,𝑖_(C of abundances for a random subset of sgRNAs (∼5%) in a dCAS9-induced, no-drug condition (concentration 0) is compared to the average abundance. For sgRNAs of medium to high relative abundance (i.e., less depletion), the CV is fairly constant at approximately 10%. However, at low relative (to uninduced) abundances (i.e. higher depletion), CV value increases substantially to over 100%. If a gene contains multiple such sgRNAs with high CV values, then the variation may be misconstrued as a genetic interaction by a methodology that is susceptible to noise.

**Fig 5.**
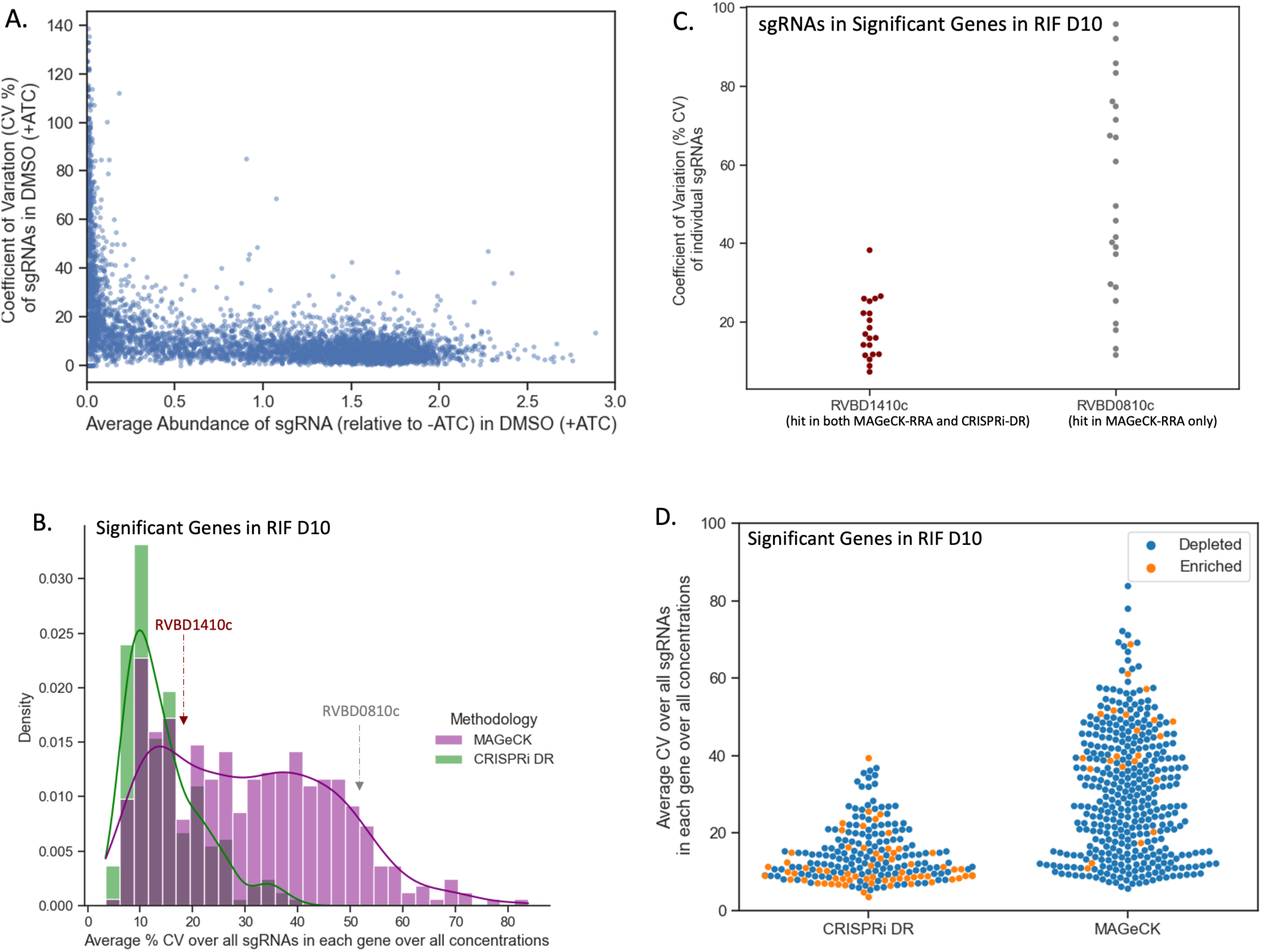
CRISPRi-DR model shows less sensitivity to noise than MAGeCK. (A) Comparison of average relative abundance and average CV across replicates in no-drug control samples for a sample of sgRNAs: For each sgRNA, we looked at the average CV of sgRNAs in the 3 control replicates against the average abundance of the sgRNA across those replicates. The lower the average abundance, the greater the noise present for the sgRNA. (B) Distribution of average CV of gene for significant genes in MAGeCK and significant genes in CRISPRi-DR in RIF D10: The distribution of average CV of significant genes in CRISPRi-DR model is more skewed and has a peak at CV ≈ 10%. Although most significant genes in MAGeCK show an average CV around 15%, there are quite a few genes with higher average CVs not found significant by the CRISPRi-DR model. (C) Coefficient of Variation (CV) of each sgRNA in two genes with similar number of sgRNAs for a library treated with RIF D10: *Rv1410c* is significant in both methodologies and *Rv0810c* significant in MAGeCK but not in CRISPRi-DR. The majority of CV values for sgRNAs in *Rv1410c* is around 20%. Although both genes have about 20 sgRNAs, *Rv0810c* shows 8 sgRNAs whose CV values exceed 60.5%, which is the maximum CV present in *Rv1410c.* (D) Distribution of average CV for enriched and depleted significant genes in MAGeCK and CRISPRi-DR in a RIF D10 library. This plot shows the distribution plot of Panel B, separated by depletion, and enriched significant genes. The average CV values for significant genes in the CRISPRi-DR model are low for both enriched and depleted genes. As seen in Panel B, significant genes in MAGeCK show low average CV, but they also show high average CV. Although there is a substantially lower number of significantly enriched in MAGeCK, they still show a large amount of noise compared the significantly enriched genes in CRISPRi-DR model.

The average noise in a gene *g* for a given drug *D* can be quantified as the average 𝐶𝑉_𝐷,𝐶,𝑖_, for all concentrations *C* and all sgRNA*s* in the gene (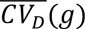). Therefore, 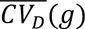 reflects the measure of overall noise present in a gene in a drug D. The distribution of 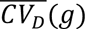 in RIF D10 for the 215 total significant genes (enriched and depleted combined) in the CRISPRi-DR model and in 218 total significant genes (enriched and depleted combined over all concentrations) in MAGeCK can be seen in Fig 5B. The distributions for both methodologies share a peak at about 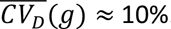. The distribution of 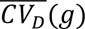 for significant genes in MAGeCK has a fatter tail than the distribution of 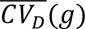 for significant genes in the CRISPRi-DR model. Fig 5D also shows that the average CV of significant genes found by MAGeCK is much higher than CRISPRi-DR (colored by depleted and enriched) for the RIF D10 screen. In addition to lower CV for significant genes, CRISPRi-DR makes more balanced calls between enriched and depleted, whereas MAGeCK calls are more asymmetric (more depleted than enriched, for this drug). This trend of higher noise in MAGeCK hits is seen not only in RIF D10, but across all the experiments conducted (See S2 Fig). This indicates that although MAGeCK is identifying genes with low noise (like the CRISPRi-DR model), it is also detecting many genes with high noise that the CRISPRi-DR model is not.

An example of such a gene is *Rv0810c*. The gene has 22 sgRNAs and has a 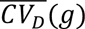 value (average CV over sgRNAs in a gene) of 51.4%, one of the highest measures in the RIF D10 experiment. In RIF D10, it is reported to be significantly depleted only in MAGeCK and not in the CRISPRi-DR model. The dispersion of the CV values of the sgRNAs in *Rv0810* are compared to those of *Rv1410c* in Fig 5C. *Rv1410c* has 20 sgRNAs, an 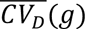 of 16.3% and is reported to be significantly depleted in both MAGeCK and the CRISPRi-DR model. Although both genes have some sgRNAs with low CVs (below 40%), *Rv0810c* shows 8 sgRNAs with CVs of at least 60.5%, which is the maximum CV of sgRNAs in *Rv1410c*. The CRISPRi-DR model considers the abundances at all concentrations, whereas MAGeCK compares each concentration to the baseline independently. Therefore, if sgRNAs have a high CV value at a particular concentration, they can be picked up as a significant genetic interaction by MAGeCK. The average relative abundance for the 3 replicates at concentration 0 for all sgRNAs in *Rv0810c* is 0.19, whereas the average relative abundance in *Rv1410c* for the same is 1.08. As Fig 5A shows, *Rv0810c* falls in the low abundance/high noise section of the graph, with an average sgRNA no-drug CV of 47.9%, whereas *Rv1410c* falls in the low noise section of the graph, with an average sgRNA no-drug CV of 11.2%. This demonstrates that MAGeCK reports genes such as *Rv0810c* with low abundances resulting in a large 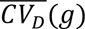, which the CRISPRi-DR model does not, i.e., MAGeCK is more suspectable to noise than the CRISPRi-DR model.

### Effects of noise on model performance using simulated CRISPRi data

The sensitivity and accuracy of the CRISPRi-DR model, MAGeCK-RRA and MAGeCK-MLE was assessed under different sources of noise using simulated sgRNA counts sampled from the Negative Binomial distribution [56], with means at different concentrations determined by the dose-response model (Eq (3)). sgRNAs with empirical efficiencies sampled from a uniform distribution from −25 to 0 were used to simulate the combined effects of CRISPRi depletion and exposure to a virtual inhibitor at four concentrations (1μM, 2μM, 4μM, and 8μM), with three replicates each. The aim was to determine how noise within and between concentrations affects the performance of each method. Detailed information on the simulation is provided in the Supplementary File S1.

Nine datasets (LL, LM, LH, ML, MM, MH, HL, HM and HH) were simulated by varying two noise parameters: variability of abundances *between* concentrations (σ*_C_*), and variability among replicates *within* a concentration (*P_nb_*, probability parameter of the Negative Binomial distribution), each with low (L), medium (M), and high (H) setting. A total of 1000 genes was simulated with 20 sgRNAs each. The first 50 genes are chosen as true negative interactions (with a virtual drug), the second 50 as positive interactions, and the last 50 as negative controls (for MAGeCK-RRA and MAGeCK-MLE). For interacting genes, slopes are chosen from a Normal distribution around +0.8 or −0.8, with a standard deviation of 0.2. For non-interacting genes, slopes are chosen from a Normal distribution around 0, with a standard deviation of 0.2. CRISPRi-DR, MAGeCK-RRA and MAGeCL-MLE were run ten times each on these 4 scenarios. MAGeCK was run independently for each drug concentration (2uM, 4uM, 8uM, compared to a no-drug control) and combined using Fisher’s method post-hoc, while CRISPRi-DR and MAGeCK-MLE were run on all four concentrations simultaneously.

In lowest noise scenario (LL = low noise between concentrations and low noise among replicates), CRISPRi-DR identified 74% of the simulated interacting genes, MAGeCK-RRA identifies 56.5% and MAGeCK-MLE identifies 99.9%. As noise increases, the recall rate of MAGeCK-MLE remains quite high at 88.3% in the highest noise scenario (HH), and MAGeCK-RRA increases to 87.5%. The recall rate of CRISPRi-DR drops down to 30.1%. However, the false positive rate of CRISPRi-DR remains low at 2.2% in this HH scenario, and the false positive rates of MAGeCK-MLE and MAGeCK increase substantially (MLE = 42.5%, RRA = 42.1%), diluting the sets of predicted enriched and depleted genes with non-interacting genes (false positives). Therefore, although CRISPRi-DR identifies less of the true interacting genes in higher noise, it maintains its ability to keep the set of reported interacting genes from being diluted with non-interacting. Across most of the 9 noise scenarios, CRISPRi-DR has higher F1-scores than the other two methods, where 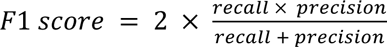, reflecting a better tradeoff between recall and precision (see Supplemental for more details).

The effect of noise on the true and false positive calls made by the methods can be seen in Fig 6, where number of significant genes is plotted for each of the adjusted noise parameters. For MAGeCK-MLE, significant genes were identified as those with adjusted P-value (based on a Wald test) less than 0.05. For MAGeCK-RRA, significant genes were identified as those with adjusted combined P-value less than 0.05 and an |LFC| greater than 1. MAGeCK-RRA is more affected by noise among replicates than between concentrations, as evident by the orange bar for *P_nb_*=0.1. This is likely a result of stochastic fluctuations of counts at individual drug concentrations that are not necessarily supported at other concentrations. This could help explain the poor performance of MAGeCK-RRA on certain drug-treated screens that may be especially noisy, resulting in many hits, such as in the case of VAN at 1 day pre-depletion; many of these hits could be false positives. Comparatively, CRISPRi-DR and MAGeCK-MLE seem to be more affected by noise between concentrations than noise between replicates, showing lower precision as σ*_C_* increases. Since these methods rely more on increasing or decreasing trends in abundance that must be (at least somewhat) consistent across concentrations, noise between concentrations may make these trends more difficult to identify.

**Fig 6.**
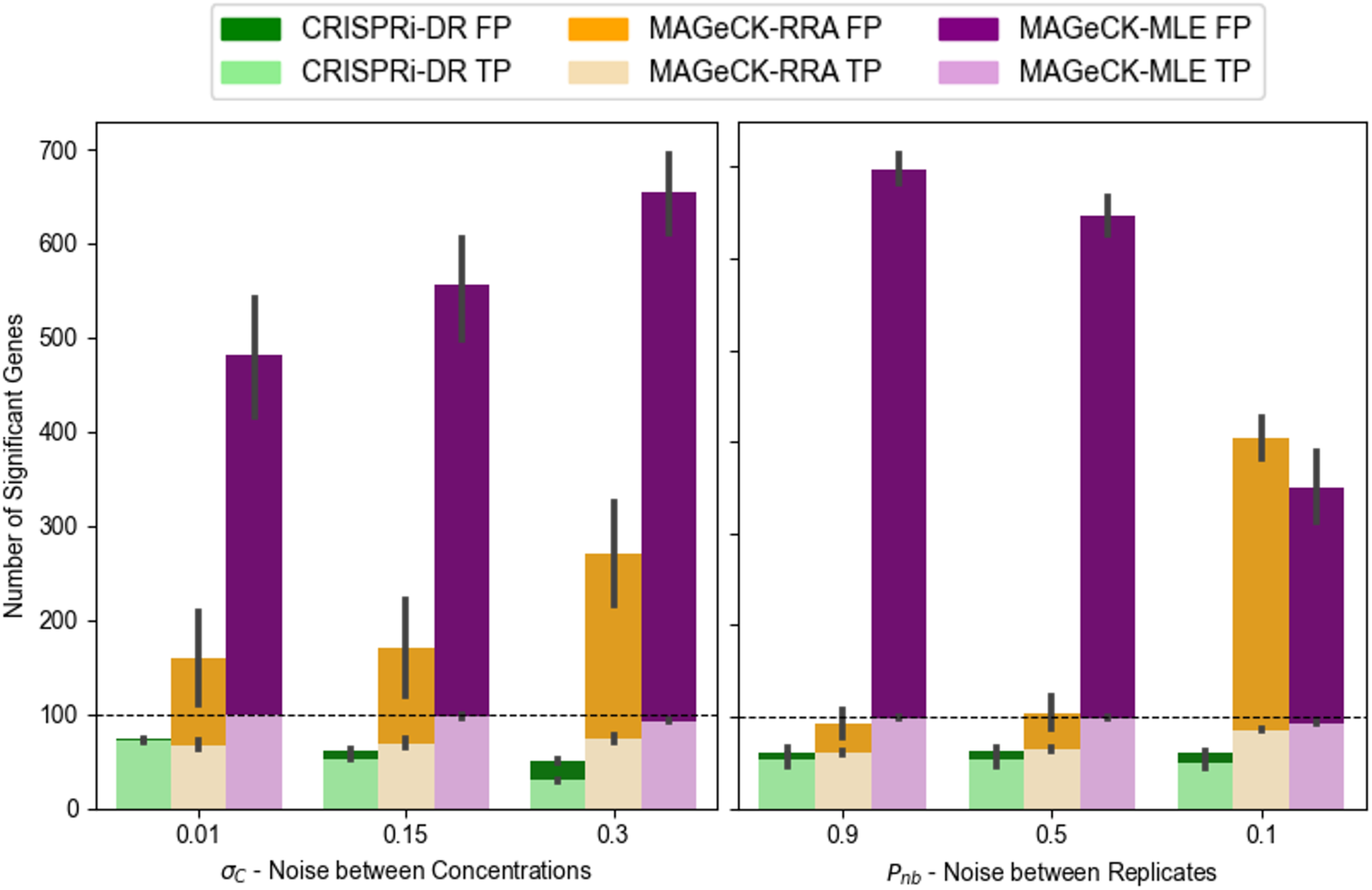
Average True Positives (TP) and False Positives (FP) found by CRIPSRi-DR, MAGeCK-RRA and MAGeCK-MLE as Simulated Noise Increases. The horizonal dashed line in both panels is the number of total simulated interacting genes (100 total). The parameters in the x-axis are ordered to reflect increasing noise. The leftmost bars of the two plots are the lowest noise and the rightmost bars are the highest noise. MAGeCK-MLE produces a high false positive rate for all scenarios and MAGeCK-RRA is more sensitive to noise among replicates as seen by the orange bar for *P_nb_*=0.1.

To assess the impact of performing a CRISPRi screen at multiple drug concentrations on the performance of CRISPRi-DR, MAGeCK and MAGeCK-RRA, we conducted the simulation above with high-noise settings (HH) and varying numbers of drug concentrations (1, 2, or 3) for 10 iterations each. The recall of the methods held fairly constant as concentrations were added. However, increasing the number of concentration points caused a significant increase in false positive calls by MAGeCK-RRA from 200 to 400. While MAGeCK-RRA shows susceptibility to false positives when evaluating only a single concentration point, this effect was amplified with more concentrations. This accumulation of errors explains the decrease in precision with additional concentration points. In contrast, CRISPRi-DR is more robust with respect to false-positive errors. By incorporating data from all available concentrations and identifying significant trends, CRISPRi-DR maintains higher precision that does not diminish with the addition of more concentration points. Although MAGeCK-MLE makes many more calls, including false positives, the number of false positives did not increase as concentrations were added, because, like CRISPRi-DR, MAGeCK-MLE incorporates data from all available concentrations.

### Comparison of CRISPRi-DR to Alternative Methods for CRISPRi Analysis

To understand how well CRISPRi-DR performs relative to other CRISPR analysis methods, we applied the following methods on the *M. tuberculosis* CGI data from [13] described above: CGA-LMM [20], MAGeCK-RRA [14], MAGeCK-MLE [15], DrugZ [17], DEBRA [18], and CRISPhieRmix [16]. Each method offers a unique approach to analyzing CRISPRi data. Some of these methods, such as CGA-LMM do not explicitly incorporate multiple sgRNAs per gene or account for differences in sgRNA strength. Other methods, such as DEBRA, MAGeCK-RRA and drugZ, do not explicitly account for different drug concentrations in a CGI experiment, and so they must be run independently on each concentration and the results combined. Only CRISPRi-DR and MAGeCK-MLE incorporate both of these factors in their statistical analysis.

The details of applying each method, including parameter settings, handling of negative controls, and merging of results, are described in the Supplement. Several of the methods, including MAGeCK-MLE, produced more significant interactions (in the thousands, in some cases), whereas other methods, like CRISPRi-DR, produced much more focused lists of significant hits for each drug (often less than 100) (see details in the Supplement).

To evaluate the accuracy of the predictions by each method, we ranked the genes by significance (usually based on P-value, for most methods) and then generated ROC (Receiver-Operator Characteristic) curves. To define a list of expected hits (i.e. interacting genes) for isoniazid (INH D1, with one day of pre-depletion), we obtained a list of 90 conditionally essential genes from a previously published TnSeq study of *M. tuberculosis* H37Rv exposed to sub-MIC concentrations of antibiotics [35]. While changes in essentiality due to knock-out of a gene by transposon insertion are not technically the same as fitness defects resulting from CRISPRi depletion of a target gene, there is substantial overlap between essentiality and vulnerability [12]. Many genes known to play a role in INH resistance (*fabG1, katG, ndh, ahpC*, *cinA,* etc.) are highly interacting (enriched or depleted) in both experiments. Thus, the list of TnSeq conditional essentials serves as a proxy for the genes that are expected to exhibit an interaction effect in the CRISPRi screen (even though, admittedly, not all necessarily will). Importantly, conditional essentiality in this context includes genes whose disruption causes either a growth defect or growth advantage (hypothetically corresponding to depletion or enrichment in a CRISPRi experiment). Similarly, to define a list of expected hits for rifampicin, we used a list of 75 conditionally essential genes based on exposure of the TnSeq library to rifampicin, which does not include subunits of the RNA polymerase because they are essential, but includes conditionally essential genes that might play a biological role in tolerating inhibition of transcription [35]. For levofloxacin (LEVO), we used 83 genes in the DNA damage-response pathway (based on the KEGG annotation [57]), plus *pafABC* (recently shown to be involved in DNA damage signaling [58]). Levofloxacin binds to the DNA gyrase (*gyrAB*), which produces a variety of types of damage to DNA, including double-stranded breaks, and requires several DNA replication and repair mechanisms to survive, such as recombination and the SOS response [59, 60]. The genes that will exhibit a chemical-genetic interaction with LEVO are likely to overlap substantially with some of the genes in this DNA damage-response pathway.

Each of the CRISPR analysis methods was evaluated using these approximate lists of expected hits for each drug. Since some of the methods were not designed to integrateinformation from multiple concentrations, the methods were initially evaluated by analyzing each concentration (LOW, MED, HIGH) of a given drug independently. Unsurprisingly, the ROC curves showed considerable dispersion of performance (Fig 7A), which was a consequence of both the method and concentration used (expected interactions were often not well-detected at low drug concentrations). Therefore, to make fairer comparisons to methods like CRISPRi-DR, CGA-LMM, and MAGeCK-MLE, we combined the results of each of the other methods over multiple concentrations by using Fisher’s method [61] to combine P-values of genes at each concentration (by summing the logs of the P-values, which is similar to taking the geometric mean) and using this to re-rank the genes. This strategy for combining results from multiple concentrations produced more uniform ROC curves for all the methods, as illustrated in Fig 7B. For methods which required a single set of counts per gene, like DEBRA and CGA-LMM, the most efficient sgRNA was chosen per gene.

**Fig 7.**
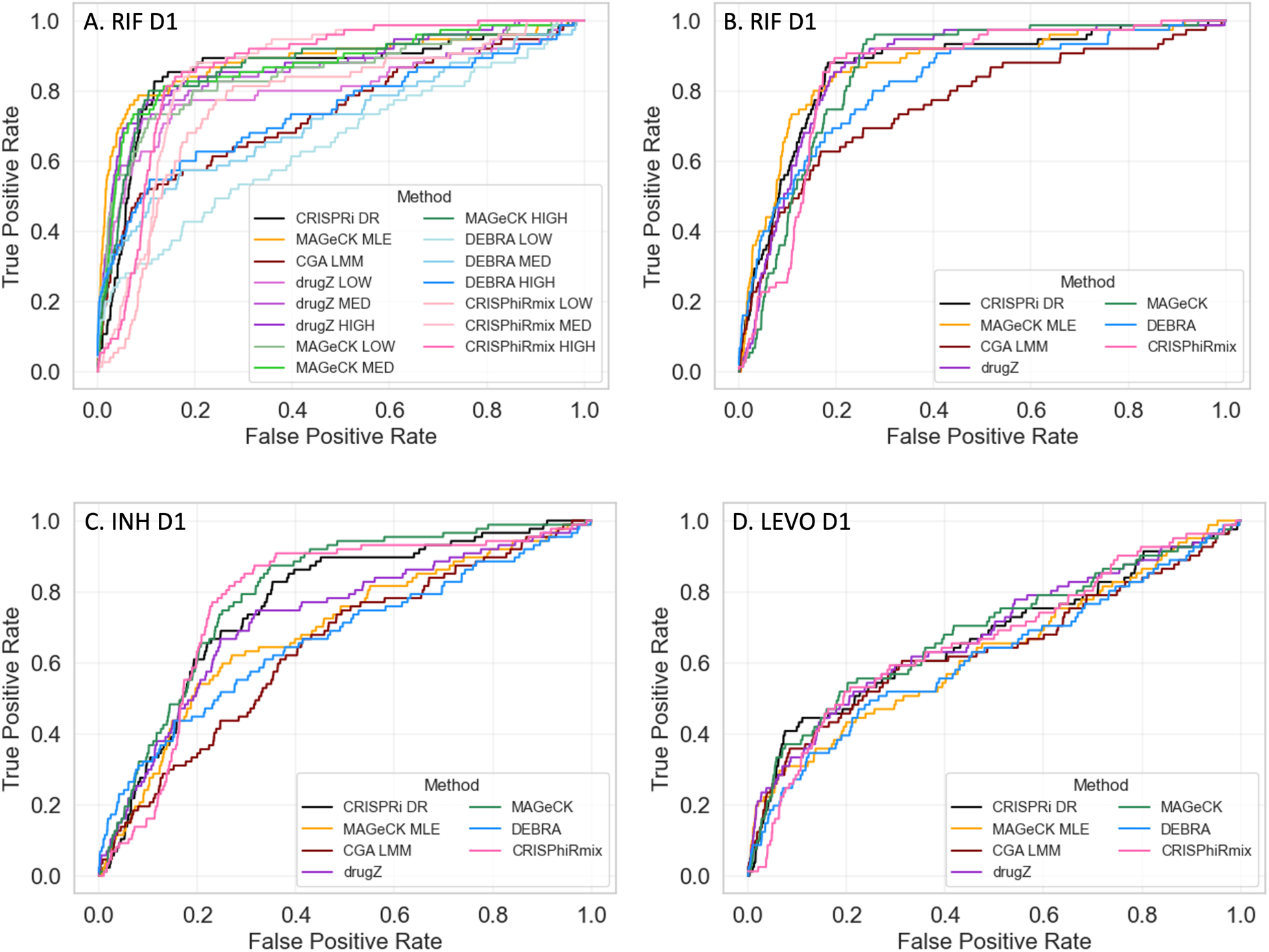
ROC Curves for RIF, INH and LEVO with 1 day pre-depletion. Using expected interactions derived from TnSeq studies [35] (INH and RIF) and the DNA-damage pathway (for LEVO), ROC Curves are plotted for CRISPRi-DR and 6 other CRISPR analysis methods. A) For methods that do not take concentration into account (MAGeCK, drugZ, DEBRA and CRISPhieRmix), each concentration (LOW, MED, HIGH) was analyzed independently, producing distinct ROC curves. B-D). For methods that do not take concentration into account, results of the 3 concentrations were combined using Fisher’s method for combining P-values.

When the results for different concentrations were combined using Fisher’s method, many of the methods exhibited reasonably good performance, ranking expected hits highly (Figs 8b-d). For example, for INH, 50% of the expected interactions were ranked in roughly the top 20% of all genes by most of the methods, and for RIF, the identification of expected interactions (based on TnSeq) was even better (producing higher rankings of expected hits). For LEVO, the ROC curves show lower AUCs for all of the methods, probably due to the fact that not all the genes in the DNA damage response pathway are required to tolerate exposure to fluoroquinolones. Though there were some variations in performance from drug to drug, indicating that differences in performance were drug-specific, the overall performance was matched fairly well, as quantified by the AUC values in Table 3. In particular, the performance of CRISPRi-DR, while not uniformly the best, was comparable to that of the other methods evaluated. It is notable methods such as CGA-LMM and DEBRA that do account for multiple sgRNAs often had the worst performance (lowest AUC values). The similarity in performance suggests that genes that exhibited CGIs (enrichment or depletion, at least at some concentration) in this experiment were easily detected by all the methods evaluated, despite their different analytical frameworks. Although the AUC values for all the methods were comparable, the other methods often reported many more false positives than CRISPRi-DR. CRIPSRi-DR tends to have slightly lower recall but much higher precision than the other methods (see Supplemental Table S2), suggesting it makes more conservative calls (see Supplement). However, it has the highest F1-scores in nearly all drug screens evaluated, which reflects the best tradeoff of recall and precision.

**Table 3.** AUC values for 7 CRISPR analysis methods, showing comparative performance on 3 datasets (drug treatments, with 1 day of pre-depletion), based on the ROC curves in Figure 7.

|  | **INH D1**<br>**AUCs** | **RIF D1**<br>**AUCs** | **LEVO D1**<br>**AUCs** |
| --- | --- | --- | --- |
| Definition of Hits: | 90 TnSeq conditional<br>essentials (Xu et al,<br>2017) | 75 TnSeq conditional<br>essentials (Xu et al.,<br>2017) | 83 genes in DNA<br>damage response<br>pathway (KEGG) |
| **CRISPRi-DR** | 0.767 | 0.850 | 0.669 |
| **CGA-LMM** | 0.641 | 0.765 | 0.638 |
| **MAGeCK-RRA** | 0.799 | 0.855 | 0.684 |
| **MAGeCK-MLE** | 0.683 | 0.865 | 0.629 |
| **drugZ** | 0.726 | 0.866 | 0.678 |
| **DEBRA** | 0.665 | 0.822 | 0.615 |
| **CRISPhieRmix** | 0.771 | 0.844 | 0.666 |

**Fig 8.**
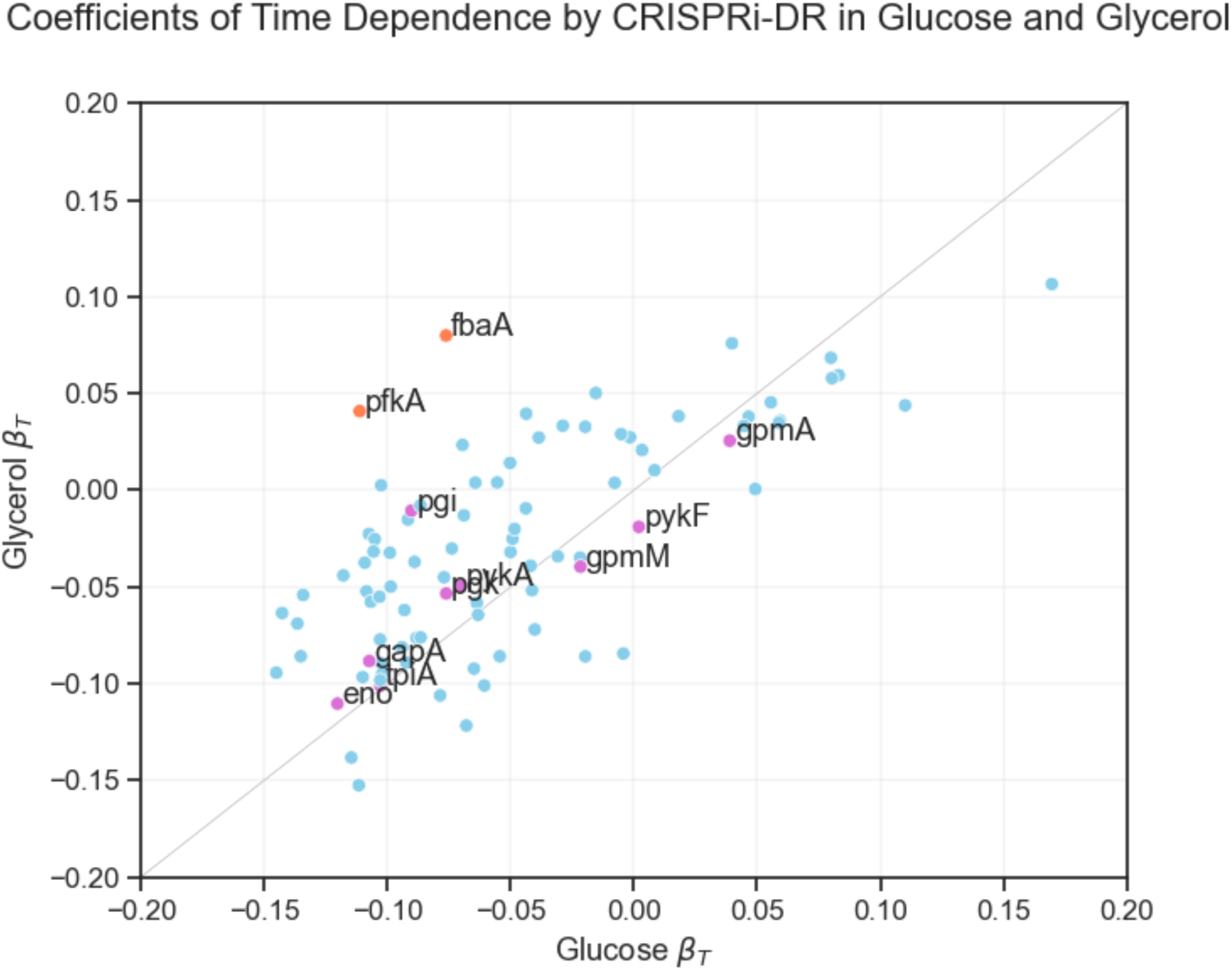
Coefficients of time dependence from CRISPRi-DR models fit for glucose and glycerol *E. coli* datasets. Each point in the scatterplot represents the coefficients of time dependence of a gene from the fit of the two models (glucose and glycerol). Individually, the gene show a range of growth defect over time, but the coefficients for most genes are equally negative for both conditions, except for a few outliers. The genes colored fuchsia are involved in both gluconeogenesis and glycolysis, hence, as expected, have similar time dependence coefficients in both carbon sources. The points farther away from this line, the orange labeled points (*pffiA* and *@aA),* are genes involved in glycolysis but not gluconeogenesis and, as expected, they have more negative coefficients in glucose than in glycerol.

### Analysis of CRISPRi Data for *E. coli* Genes Required for Growth on Different Carbon Sources

To illustrate the application of the CRISPRi-DR method to other datasets, we re-analyzed the data from a CRISPRi library in *E. coli* that was used to investigate differential requirements for growth on glycerol versus glucose as a carbon source [11]. While this is not technically a chemical-genetics experiment, the data included multiple time points. The growth curves of CRISPRi knock-down mutants (depletion over time) follows sigmoidal behavior very analogous to dose-response curves for antibiotic exposure (depletion with increasing concentration). Furthermore, while only 88 genes were analyzed instead of a whole-genome screen, this dataset is suitable for analysis by CRISPRi-DR because multiple unique sgRNAs were synthesized for each gene (68 per gene on average), spanning a range of efficiencies (which were quantified by fitting growth data to a logistic curve).

We ran CRISPRi-DR on this data for each carbon source independently (fitting the model to 7 timepoints for glucose, 5 for glycerol) (see Supplemental Material for additional details). Many genes exhibited significant depletion effects (reduced fitness), because many of the 88 genes were essential for growth (on either carbon source). However, when the coefficients of the time parameter from the CRISPRi-DR analysis were plotted as a scatter plot between the carbon sources, two genes stood out as being preferentially required for growth on glucose (highlighted in orange in Fig 8, most divergent from the diagonal): *@aA* (fructose bisphosphate aldolase) and *pffiA* (phosphofructokinase). These genes are well-known examples required for preliminary steps in glycolysis but not for incorporation of glycerol, and were identified in the analysis by [11]. Additional metabolic genes needed for growth on both carbon sources are observed to lie along the diagonal. This demonstrates that the CRISPRi-DR method can be applied to other datasets, including those not explicitly designed for chemical-genetics. The modified dose-response model nicely incorporates the simultaneous effects of time and the variable efficiency of sgRNAs on mutant abundance.

## Discussion

There are a variety of ways to use CRISPRi technology for probing the biological roles of genes by modulating their expression levels in-situ. While early experiments utilized the intrinsic nuclease activity of the CAS9 to knock-out genes entirely [1–3], more recent approaches have enabled partial knock-down of targets, generally using an inactive CAS9 (dCAS9) to bind to target genes and block transcription [5]. One way of controlling the level of depletion is through manipulating the expression of the dCAS9 itself. However, a second approach to creating variability in levels of target depletion is to utilize multiple sgRNAs of different efficiency. The nucleotide sequence of both the PAM and target-specific parts of the guide RNA can impact the hybridization and recruitment of the dCAS9 [9, 10]. This variability can be useful for gauging or titrating phenotypic effects. Rather than all-or-none responses, one can look for genes whose level of depletion correlates with the phenotype of interest.

While CRISPRi libraries can be constructed with multiple sgRNAs per target, most CRISPR analytical methods do not explicitly handle such, and those that do (such as MAGeCK-RRA and CRISPhieRmix) are essentially designed to identify significant genes by focusing on a subset of apparently effective sgRNAs (i.e. allowing for ineffective sgRNAs, which are filtered out for each target). However, sgRNA efficiency can be quantified a priori, such as by running a growth experiment to determine the fitness effect of inducing the depletion of the target gene. If this information is available (collected beforehand), then it can be incorporated into the analysis as a “covariate”, to enable comparison of the impact of treatment conditions on the expected magnitude of the phenotypic effect. We note that sgRNA efficiency is different than predicted strength, because it also depends on the vulnerability of the gene. In an essential gene, some sgRNAs might be more efficient than others. In contrast, typically, all the sgRNAs targeting a non-essential will turn out to be non-efficient (i.e. have 0 growth defect, or relative fitness of around 1), at least under control conditions, since the cells are unaffected by depletion of these proteins and continue to grow at the same rate. However, they might cause growth impairment if expressed in certain stress conditions where they might play a role in survival/tolerance. In fact, in chemical-genetic interaction experiments, variable sgRNA efficiency can be further exploited to identify genes whose level of depletion synergizes with increasing drug concentration. We developed the CRISPRi-DR model with this use case in mind, extending the Hill equation, which quantifies dose-response behavior of a growth inhibitor, to incorporate an extra term representing the relative efficiency of each of the sgRNAs targeting a gene. This approach, however, is not limited to CGI experiments. It can be applied to other treatments that induce a sigmoidal response. For example, in re-analysis of data from the Mathis, Otto and Reynolds (11) paper, we showed the same equation could be adapted for modeling the effect of *E. coli* cultures grown on medium with different carbon sources; the time parameter could be substituted for the concentration, since depletion of essential genes caused a gradual killing with an S-curve shape over time.

Therefore, the CRISPRi-DR approach we developed has 3 main requirements. First, the CRISPRi library should contain multiple sgRNAs per target gene. Anecdotal evidence suggests that at least 5 sgRNAs per gene are necessary to maintain overall sensitivity for detecting expected interactions and maximizing AUC (based on experiments where we subsampled a limited number of sgRNAs per screen; see Supplement). Fewer sgRNAs per gene reduced the stability of the regression and increased variance of the fitted parameters (specifically the slope of concentration dependence). Second, ideally, sgRNAs of differing strength should be included. Strength can be predicted from sequence features using various types of trained models [9, 12]. This covers both essential and non-essential genes. For essential (or vulnerable) genes, sgRNA efficiency correlates with predicted strength, so this is equivalent to choosing sgRNAs with a range of efficiencies (that create varying growth defects). For non-essential genes, one could choose a set of sgRNAs with a range of predicted strengths, even though they might all turn out to be non-efficient experimentally in standard growth conditions. This diversity could be created by selecting sgRNAs that deviate from the optimal PAM sequence [6], choosing hybridizing sequences of different length or GC content [5, 8], or adding random nucleotide substitutions [10]. Third, the actual efficiency of each sgRNA must be empirically quantified a priori, such as by running a growth experiment and comparing growth rates with and without induction of the dCAS9 (hence, with and without depletion of target genes). These quantities become inputs to the model. The CRISPRi-DR method can be applied to any CRISPRi dataset that meets these requirements. The methodology works best when treatment produces a sigmoidal effect on mutant abundances.

Doench, Fusi (9) have proposed several systems for design/optimization of CRISPRi libraries. These were more focused on minimizing off-target effects while maximizing sensitivity for detecting of genuine interactions. They do not give a specific recommendation about how many sgRNAs per gene to select. Their library design guidance is to prefer more efficient sgRNAs (e.g. Rule Set 1 selects top 20% of sgRNAs by empirical efficiency and uses these to build a model to predict sgRNA strength; Rule Set 2 extends this with a machine learning model based on additional sequence features to predict sgRNA strength, and prefers sgRNAs with highest score [9]). This contrasts with our approach, where we advocate selecting sgRNAs with a diversity of efficiencies, since we observed that the sgRNAs that exhibited the most synergy with drug treatments were not always the strongest or weakest, but somewhere in the middle of the range.

For application to CGI experiments, the availability of CRISPRi data for multiple sgRNAs of varying strengths for each target gene presents new challenges for statistical analysis. In previous work [20], we showed that regressing the relative abundances of mutants in hypomorph libraries over multiple concentrations of a drug (on log-scale) can be used to improve detection of CGIs. This regression approach captured dose-dependent behavior, i.e. genes whose decreased expression caused either suppressed or enhanced fitness that increases in magnitude with drug concentration (i.e. exhibits a trend, which is important for statistical robustness). The CRISPRi-DR method described in this paper extends this previous work by showing how effects of both drug concentration and sgRNA efficiency can be accommodated in the same model. Ideally, interacting genes would be expected to exhibit synergistic behavior with a drug, where depletion of a target protein induces excess depletion (or enrichment) of the mutants grown in the presence of an inhibitor, and this effect is concentration-dependent (exhibits dose-response behavior).

In theory, both CRISPRi depletion of essential genes and exposure to antibiotics should impair growth of CRISPRi mutants (at least for depletion of essential genes). One might expect to observe a depletion effect due to either increasing sgRNA efficiency, or drug concentration, each producing regression “slopes” (in log-transformed space), with slopes for sgRNAs targeting non-essential genes being expected to be flat, regardless of predicted sgRNA strength. However, we observed that sgRNA efficiency and concentration effects are not independent -they interact in a non-linear way. sgRNAs that are too weak do not produce enough depletion of a drug target to cause sensitization, and sgRNAs that are too strong deplete a mutant to such low abundances that concentration-dependent effects are difficult to quantify. Often, there is a “sweet spot”, or an intermediate sgRNA strength which maximizes the concentration-dependent effect (which could be different for each gene). Our CRISPRi-DR model incorporates both sgRNA efficiency and drug concentration as parameters, and reproduces the non-linear interaction between them, where the “slopes” for the effect of drug concentration on relative abundance of mutants can be larger in magnitude for sgRNAs of intermediate strength, while being flatter (slopes closer to 0) for sgRNAs of high or low strength. MAGeCK-MLE is the only other analytical method that take sgRNA efficiencies as an input; in that method, the empirical measures of efficiency are used to initialize the prior probability that each sgRNA is effective (assuming each gene is represented by a subset of sgRNAs that are effective and others that are not), which is combined with other conditional probabilities in a Bayesian framework to determine the posterior probability of interaction for each gene. However, we observed that MAGeCK-MLE often reports far more significant interactions that CRISPRi-DR or several other methods and has lower precision.

In this paper, we showed that this non-linear interaction between sgRNA efficiency and drug concentration can be modeled using an augmented dose-response equation, in which terms for both effects are included. By fitting the parameters in this equation to CRISPRi data from a CGI experiment (normalized mutant abundances from sgRNA counts), one can estimate the degree to which depletion of a given gene sensitizes cells to an inhibitor, and thereby identify CGIs. While various computational methods exist for fitting non-linear equations, such as the Levenberg–Marquardt algorithm [62], we chose to linearize the modified Hill equation by applying a log-sigmoid transform. The transformation enables us to express the equation in a linear form, where the parameters (IC_50_, Hill slopes, etc.) appear as coefficients of linear terms or constants. Consequently, we can use ordinary least-squares regression (OLS) to fit the model to the CRISPRi dataset.

Sometimes positive and/or negative controls are included in a CRISPRi experiment [8]. While negative controls can be used in methods like MAGeCK-RRA, CRISPRi-DR is not designed to use controls explicitly in the statistical analysis of CGIs. Hypothetically, negative controls could be used in the final filtering step to calculate Z-scores for each gene. Instead of basing the Z-scores on the mean and standard deviation of slope coefficients in the whole set of genes, they could be based on the distribution of slope coefficients from the negative controls. While we tested this idea (using 1750 non-targeting sgRNAs included in the *Mtb* CRISPRi dataset as negative controls), it resulted in many more genes being labeled as interactions (up to half the genome). It appears that unrelated genes (not involved in the mechanism of action or resistance to a drug) often have slightly positive or negative random slopes, due to some source of noise in the experiment that is unaccounted for. Some genes could exhibit weak phenotypic effects, conferring slight growth defects or advantages under antibiotic stress, even though they do not play any direct role in the mechanism of action or resistance to the drug. This is the reason that we advocate identifying genes that are outliers with respect to the rest of the population of genes, achieved through the filtering step at the end (|Zscore|>2), instead of just reporting all genes with slope coefficient statistically different from 0.

We compared CRISPRi-DR to several other analytical methods, including MAGeCK-RRA, MAGeCK-MLE, DEBRA, CRISPhieRMix, CGA-LMM, and drugZ. Some of these methods incorporate multiple drug concentrations, while other incorporate sgRNA efficiency as an input to their models. However, only MAGeCK-MLE incorporates both types of input. The importance of incorporating both inputs in CRISPRi-DR was demonstrated via an experiment with ablated models; the model fits (AICs) for each gene were significantly worse for models that regressed abundances against either drug concentration or sgRNA efficiency alone. For those methods that do not explicitly combine data from multiple drug concentrations and must be run on each concentration independently, we employed Fisher’s method of combining P-values to create a merged ranking of genes. Using ROC curves to comparing ranking of expected interactions, CRISPRi-DR performed comparably to the best of these methods, though method with the highest AUC differed depending on the drug. This evaluation was facilitated by using lists of conditionally essential genes from TnSeq experiments (exposure to same drugs) to define an objective list of expected interactions for each drug for making fair comparisons of performance. However, a major difference observed among the methods was in the number of significant interactions detected. Methods like CRISPRhieRMix, DEBRA, MAGeCK-RRA, and MAGeCK-MLE produced hundreds to thousands of hits for each drug, whereas CRISPRi-DR reported a more conservative list of typically less than a hundred interacting genes. It is likely that many of the interactions detected by the former methods could be false positives. This was borne out in simulation experiments, where MAGECK-RRA, and MAGeCK-MLE exhibited substantially lower precision than CRISPRi-DR. In both the simulated data and real drug screen datasets, CRISPRi-DR had the highest F1-scores, reflecting the best tradeoff between precision and recall compared to other methods. Reducing false positives is important because experimental validation of hits can be expensive, and follow-up is usually only applied to a handful of top-ranked genes. Furthermore, we used simulated datasets to explore how noise within or between drug concentrations could affect both the recall and precision of CRISPRi-DR, MAGECK-RRA, and MAGeCK-MLE. Both types of noise increasingly degrade the recall of all methods, but noise within concentrations (i.e. sgRNA counts among replicates) seemed to cause the greatest decrease in precision, especially for MAGeCK-RRA. The outlier analysis in CRISPRi-DR (filtering by Z-score in the last step) partially helps to mitigate this, producing a more focused list of candidate interactions, and hopefully eliminating genes with small random slopes of concentration dependence that are not genuine interactions (i.e. false positives).

## Data and Code Availability

A python-based implementation of the CRISPRi-DR method for analyzing CRISPRi data is publicly available as part of Transit2: https://transit2.readthedocs.io/en/latest/

The output files from analyses of the *Mtb* CRISPRi CGI screens from Li, Poulton (13) using CRISPRi-DR are available for download at: https://orca1.tamu.edu/CRISPRi-DR/

## Acknowledgments

This work was supported by NIH grant P01 AI143575 (TRI, JR, and DS) and by grant INV-004761 from the Bill and Melinda Gates Foundation (DS and TRI). The funders had no role in study design, data collection and analysis, decision to publish, or preparation of the manuscript.

## Supporting Information

**Fig S1.**
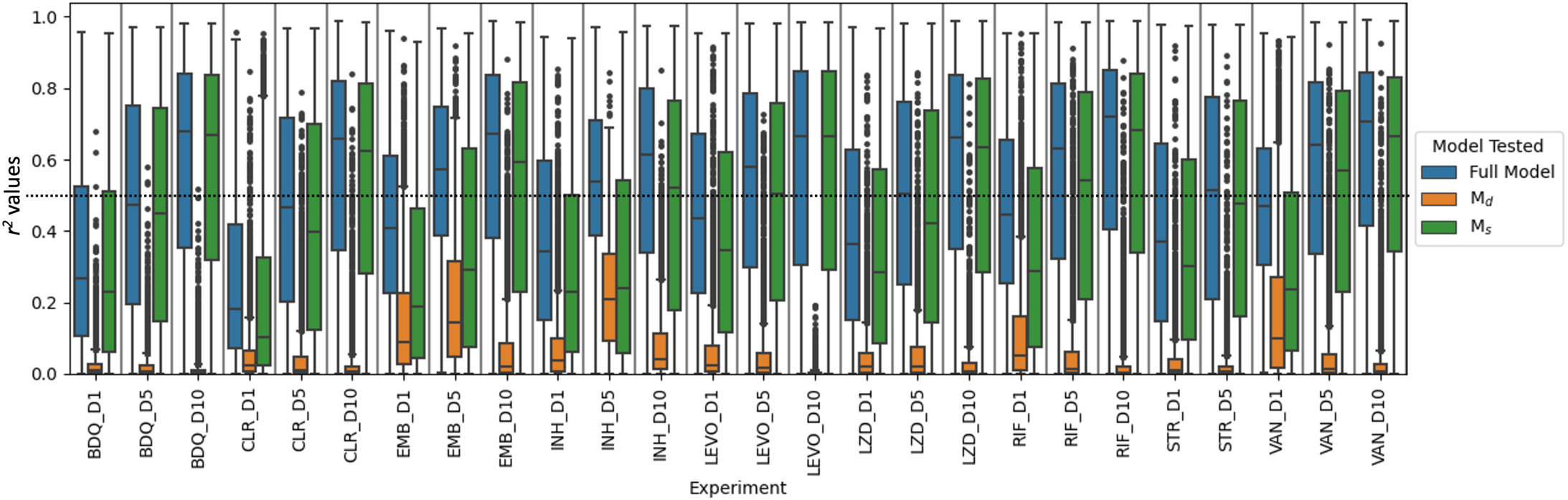
Evaluation sgRNA strength and log concentration as predictors of CRISPRi-DR model through comparison of distribution of *r^2^* values of full (CRISPRi-DR) and ablated (M_s_ and M_d_) models for each gene in each experiment. The horizontal line is where *r^2^* = 0.5. The average *r^2^* M_s_ model for all genes across all the experiments is 0.42, the average *r^2^* for the M_d_ model is 0.07. This alongside the Log-likelihood tests indicate sgRNA strength is the more significant predictor. However, the full CRISPRi-DR model outperforms both M_d_ and M_s_ (average *r^2^* is 0.50) indicating the inclusion of both sgRNA strength and log concentration is needed for accurate assessment of significant sgRNA depletion in a gene in a condition.

**Fig S2.**
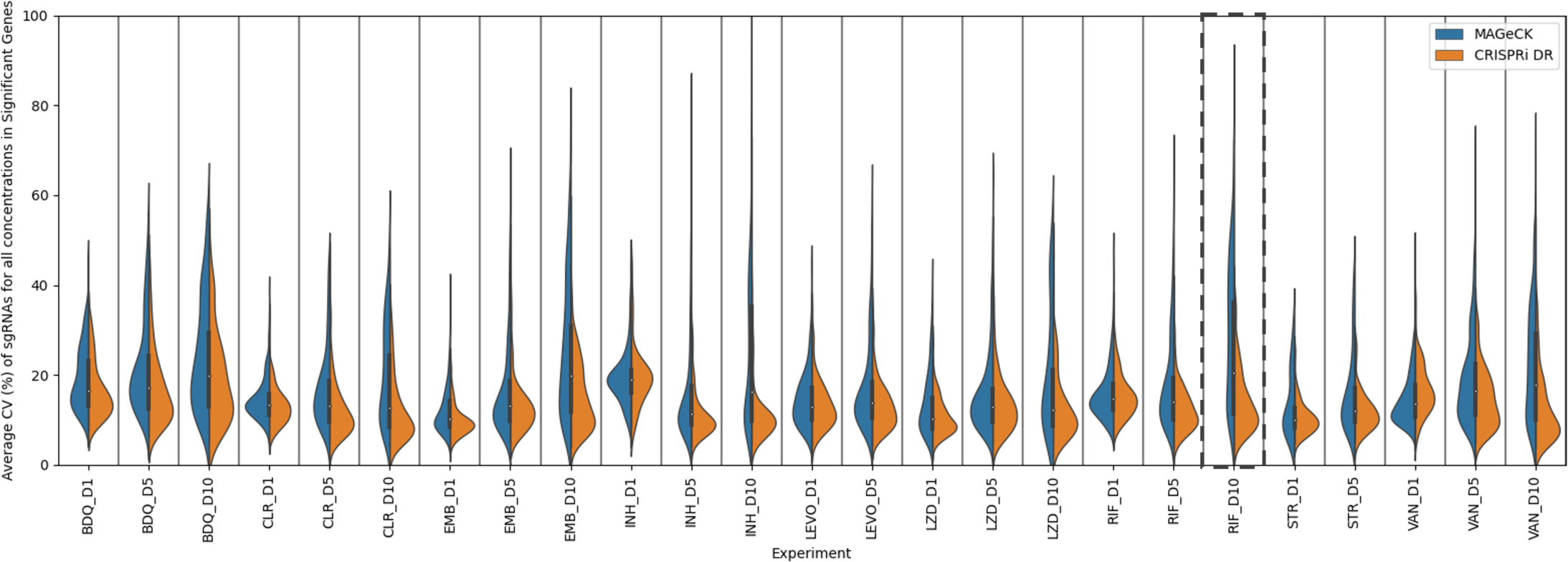
Distribution of average CV of sgRNAs in significant genes (depleted and enriched) in the CRISPRi-DR model and MAGeCK. In this Fig, we see all the noise distributions for hits in MAGeCK and the CRISPRi-DR model for all experiments. The dashed panel is that of RIF D10. The same distribution of noise of hits can be seen in Fig 5. The trend seen with RIF D10 is present with all the experiments except LEVO D10. We see that the CRISPRi-DR model is unimodal with a low CV as the mode, whereas MAGeCK shows significant genes with low average CV values but also a significant amount of genes with high average CV values. LEVO D10 was left out of this plot due to the low number of hits in either model.

**Table S1. Ranking of Select Genes using the CRISPRi-DR model in 1 Day, 5 day and 10 Day pre-depletion of treated libraries.**

An extended version of Table 2, where the CRISPRi-DR model is run on each gene for each drug and pre-depletion day. The coefficient for the slope of concentration dependence (𝛽_𝑐_) is extracted from the fitted regressions and used to rank the genes in both increasing order (for depletion) and inversely (for enrichment). Green reflects results consistent with expectations based on knowledge of known gene-drug interactions.

**Table S2. Comparison of significant interactions Identified by CRISPR analysis methods of EMB, INH, LEVO, VAN and RIF CRISPRi screens**

For each drug and pre-depletion day of the selected datasets, all 7 CRIPSR methods were run. For methods that do not account for multiple concentrations, they were run separately for each concentration and the overall significant interactions are also addressed post-combination of the individual runs using Fisher’s method. The comparison of the significant interactions identified by the models was evaluated using an objectively defined list of true positives. The genes identified by Xu, DeJesus (35) were used as the “ground truth” against which the other model’s results were compared. For LEVO, genes in the DNA Damaging pathway are used. Recall, Precision and F1-score columns are colored such that higher values are more green.

**Table S3. Matrices for comparison of significant interactions Identified by CRISPRi-DR and MAGeCK for each drug and pre-depletion day.**

The table presents the results of CRISPRi-DR and MAGeCK analyses for different drugs and pre-depletion days. Significant interactions are compared in matrix form. Cells with red font indicate low overlaps between the interactions found by the two models, while cells with green font represent high overlaps.

**Supplemental File S1**

We expand on the following four topics from the main text in this document: 1) An assessment of CRISPRi-DR, MAGeCK and MAGeCK-MLE on datasets with simulated noise, 2) Comparison of CRISPRi-DR to other analysis methods using CGI datasets, 3) Analysis of *E. coli* CRISPRi screens using CRISPRi-DR and, 4) The minimum number of sgRNAs recommended per gene in CRISPRi-DR.

