## Supplemental File S1 for "A dose-response model for statistical analysis of chemical genetic interactions in CRISPRi screens"

### Supplementary Material

We expand on the following four topics from the main text in this document:

1. An assessment of CRISPRi-DR, MAGeCK and MAGeCK-MLE on datasets with simulated noise (p. 1)
2. Comparison of CRISPRi-DR to other analysis methods using CGI datasets (p. 15)
3. Analysis of *E. coli* CRISPRi screens using CRISPRi-DR (p. 23)
4. The minimum number of sgRNAs recommended per gene in CRISPRi-DR. (p. 27)

### 1. Assessment of CRISPRi-DR, MAGeCK and MAGeCK-MLE on datasets with simulated noise

#### Simulation Design

To better understand the differences in performance, between CRISPRi-DR, MAGeCK-RRA, and MAGeCK-MLE, and asses their sensitivity to various sources of noise, we developed a hierarchical simulation model.

In this experiment, we simulated 1000 genes with 20 sgRNAs each. The first 50 genes are chosen as true negative interaction (with a virtual drug), the second 50 as positive interactions, and the last 50 as negative controls (for MAGeCK-RRA and MAGeCK-MLE). We simulated exposure to a virtual inhibitor over 4 concentrations (1μM, 2μM, 4μM, and 8μM), 3 replicates each.  Our objective was to quantify how much noise in the counts, both within concentrations and between concentrations, affects the precision and recall of each method.

We sample parameters for baseline abundances, sgRNA efficiencies, and drug sensitivities (concentration-dependence) from prior distributions. Then, we use the dose-response model (sigmoid transformation of linear model) to generate mean counts for each sgRNA at each concentration (see Eq. 1 below). Finally, we draw multiple replicates of actual counts surrounding these means from a Negative Binomial distribution while creating variance/noise in the observations (Eq. 2 below).

<u>Sampling Baseline Counts</u>

Baseline barcode counts were generated for each sgRNA *j* by sampling from a LogNormal distribution:

$$B_j \sim LogNormal(shape = exp(5), scale = 1)$$

This produces baseline counts in the range of single digits up to a few thousand, with a mean count in the hundreds (see histogram), which is typical of what is seen in real CRISPRi sequencing datasets.

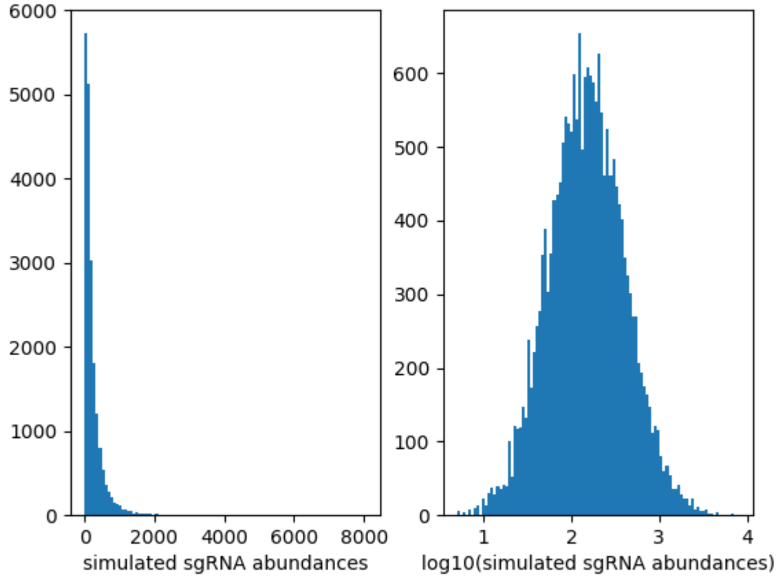


Sampling sgRNA efficiencies

To simulate the effect of CRISPRi depletion, an efficiency is chosen for each individual sgRNA $j$ in each gene $i$ from a uniform distribution:

$$S_{ij} \sim U(-25,0)$$

This range was chosen based on the data from the *Mtb* CRISPRi library in the Li, Poulton (1) paper, which was based on empirical estimates of growth rates (fitness defects), fit to a piecewise linear model and extrapolated to predicted LFCs at 25 generations [2]. Since sgRNA efficiencies empirically span a range of -25 to 0, and we want 0 to represent no depletion and -25 to represent high depletion (in induced vs. uninduced).

Sampling concentration-dependent slopes

Next, concentration-dependent coefficients (slopes) are chosen for each gene $i$. For interacting genes, slopes are chosen from a Normal distribution around $+K_2$ or $-K_2$, where $K_2$ is a parameter. For non-interacting genes, slopes are chosen from a Normal distribution around 0, with a standard deviation of $\sigma_{K2}$. The larger the variance, the more risk there is the slopes of non-interacting genes overlapping with interacting genes.

$$M_i \sim \begin{Bmatrix} \mathrm{N}(0, \sigma_{K2}^2) \text{ if } i \text{ is a non} - \text{interacting gene} \\ \mathrm{N}(+\mathrm{K}_2, \sigma_{K2}^2) \text{ if } i \text{ is a positive interaction} \\ \mathrm{N}(-\mathrm{K}_2, \sigma_{K2}^2) \text{ if } i \text{ is a negative interaction} \end{Bmatrix}$$

Using $K_2$=0.8 and $\sigma_{K2}$=0.2, the histogram below shows the overlapping distributions of slopes for the 3 types of genes.

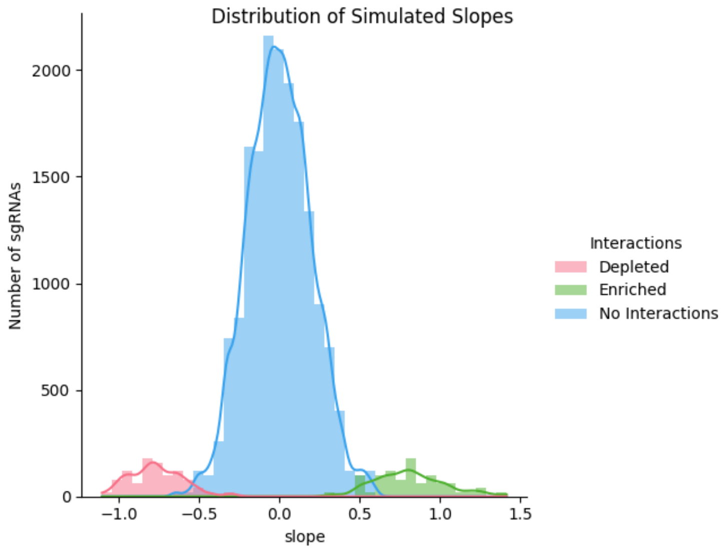


Simulation of sgRNA counts

Next, mean abundances for each sgRNA at each concentration were generated by sampling from the dose-response equation. First, a linear model was used to estimate a level which was then transformed to an abundance ($A_{ijc}$) by the sigmoid transformation, σ, reproducing the modified dose-response equation described in the main text.

$$A_{ijc} \ = \ B_j * \sigma( \, K_0 + K_1 * S_{ij} + M_i * log(\frac{c}{q}) \, ) \tag{1}$$

$$\sigma(x) = \frac{exp(x)}{1 + exp(x)}$$

Four concentrations were simulated: 1, 2, 4, 8 times the $IC_{50}$ (q). Thus, log-concentrations used in the linear model were 0, 1, 2, 3. The $IC_{50}$ is represented by q; a constant $IC_{50}$ of q=2.0 was used.

The coefficients $K_0$ and $K_1$ were set to 3 and 0.3, respectively, to simulate sensitivity to protein depletion.

Finally, given the level of abundance for each sgRNA at each concentration, 3 replicates (x) were drawn from a Negative Binomial distribution. The variability (noisiness) of the raw counts could be adjusted by manipulating the probability parameter p=$P_{nb}$, which was combined with the target mean abundance ($A_{ijc}$) to determine the size parameter $r = A_{ijc} * \frac{Pnb}{1-Pnb}$ for the sampling:

$$Y_{ijcx} \sim NegativeBinomial(p, r) \tag{2}$$

$$p = Pnb \ ; \ r \ = \ A_{ijc} * \frac{p}{1-p}$$

Simulating noise between concentrations

In addition to noise between replicate counts (controlled by $P_{nb}$) and noise affecting drug sensitivity (variability of slopes, controlled $\sigma_{K2}$), another source of noise was simulated by randomly shifting the abundance at each concentration, where the shifts were sampled from a Normal distribution, inflating or deflating the expected counts at a given concentration, deviating from a perfect linear trend. This was modeled as a gene-level effect using the parameter $\sigma_C$ controls the noise between concentrations.

$$D_{ic} \sim N(0, \sigma_C)$$
$$A_{ijc}' = exp(D_{ic}) * A_{ijc}$$

**Results**

The other parameters are set as the following:
- sgRNA efficiency sensitivity: $K_0$=3, $K_1$=0.3
- concentration dependence: q=2, $K_2$=0.8, $\sigma_{K2}$=0.2

We generated simulated datasets by varying the 2 noise parameters:
- noise between concentrations: low: $\sigma_C$ =0.01, med: $\sigma_C$ =0.15 ,high: $\sigma_C$ =0.3
- noise between replicates: low: $P_{nb}$=0.9, med: $P_{nb}$=0.5, high: $P_{nb}$=0.1

By forming all combinations of these parameters, we generated the following 9 scenarios:
- LL (Low noise between concentrations, Low noise within concentrations): $\sigma_C$ =0.01, $P_{nb}$=0.9
- LM (Low noise between concentrations, Medium noise within concentrations): $\sigma_C$ =0.01, $P_{nb}$=0.5
- LH (Low noise between concentrations, High noise within concentrations): $\sigma_C$ =0.01, $P_{nb}$=0.1
- ML (Medium noise between concentrations, Low noise within concentrations): $\sigma_C$ =0.15, $P_{nb}$=0.9
- MM (Medium noise between concentrations, Medium noise within concentrations): $\sigma_C$ =0.15, $P_{nb}$=0.5
- MH (Medium noise between concentrations, High noise within concentrations): $\sigma_C$ =0.15, $P_{nb}$=0.1
- HL (High noise between concentrations, Low noise within concentrations): $\sigma_C$ =0.3, $P_{nb}$=0.9
- HM (High noise between concentrations, Medium noise within concentrations): $\sigma_C$ =0.3, $P_{nb}$=0.5
- HH (High noise between concentrations, High noise within concentrations): $\sigma_C$ =0.3, $P_{nb}$=0.1

Fig 1 illustrates the different effects of simulated noise between concentrations vs. within concentrations (among replicates) in the low-high combination scenarios, for representative sgRNAs in negative interacting genes. The decreasing trend of counts is more variable in HL and HH than in the LL and LH scenarios. Whereas the dispersion of counts within a concentration is greater in LH and HH. The medium noise scenarios (LM, ML, MM, MH, HM) are somewhere in between these four levels of dispersions.

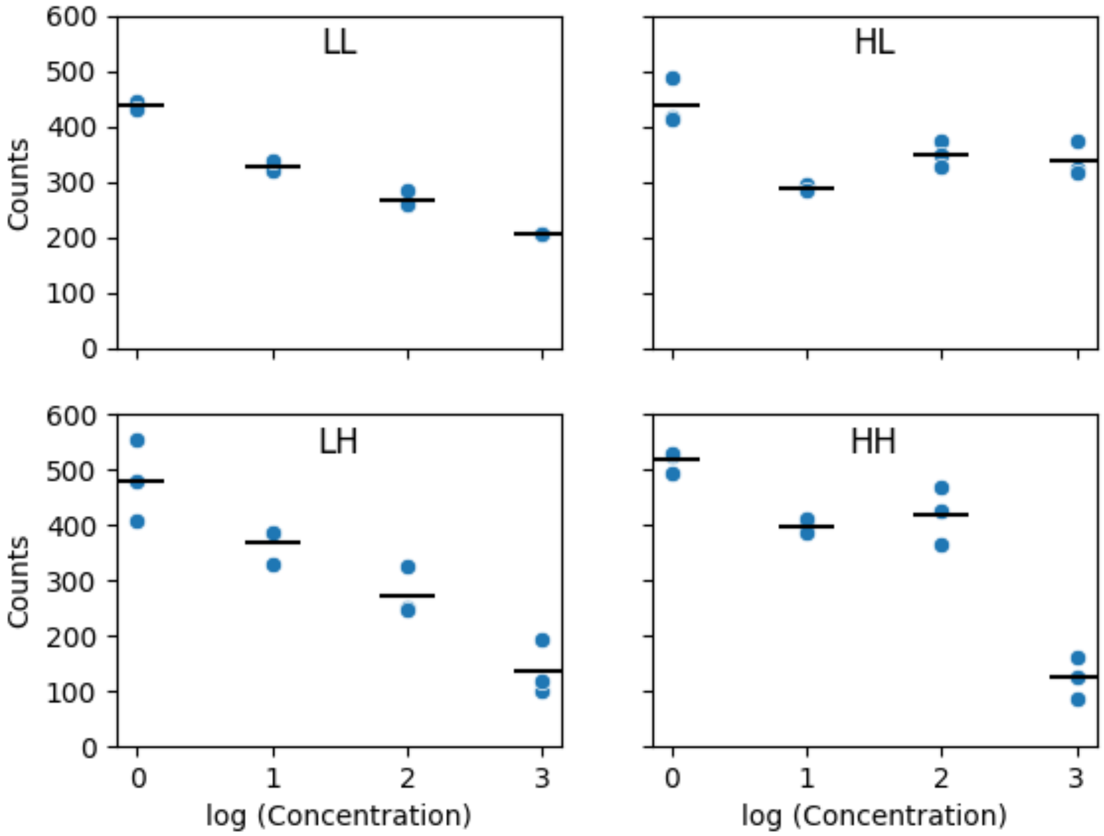


**Fig 1. Abundances of select sgRNA(s) in the low and high noise scenarios.** For each noise scenario, the counts of a representative sgRNA from simulated negative interacting genes are seen here with the means across replicates marked by the horizontal black line. The medium noise scenarios are somewhere in between levels of dispersion depicted.

We analyzed all 9 scenarios with MAGeCK-MLE, MAGeCK-RRA and CRISPRi-DR. For CRISPRi-DR, the criteria used to identify significant interactions was adjusted P-value<0.05 and $|Z_{slope}|>2$, as described in the main text. For MAGeCK-MLE, the criteria used to identify significant interactions was adjusted P-value based on Wald < 0.05. MAGeCK was run 3 times independently for each drug concentration: 2 μM, 4 μM, 8 μM. Each was compared to the simulated no-drug control (DMSO). A single P-value per gene was calculated from the three analyses using the Fisher's method, which was then adjusted using Benjamini-Hochberg. Additionally, a single LFC value per gene was set equal to the most significant LFC across the three concentrations. Significant interactions were classified as combined adjusted P-value < 0.05 and |gene LFC| >1.

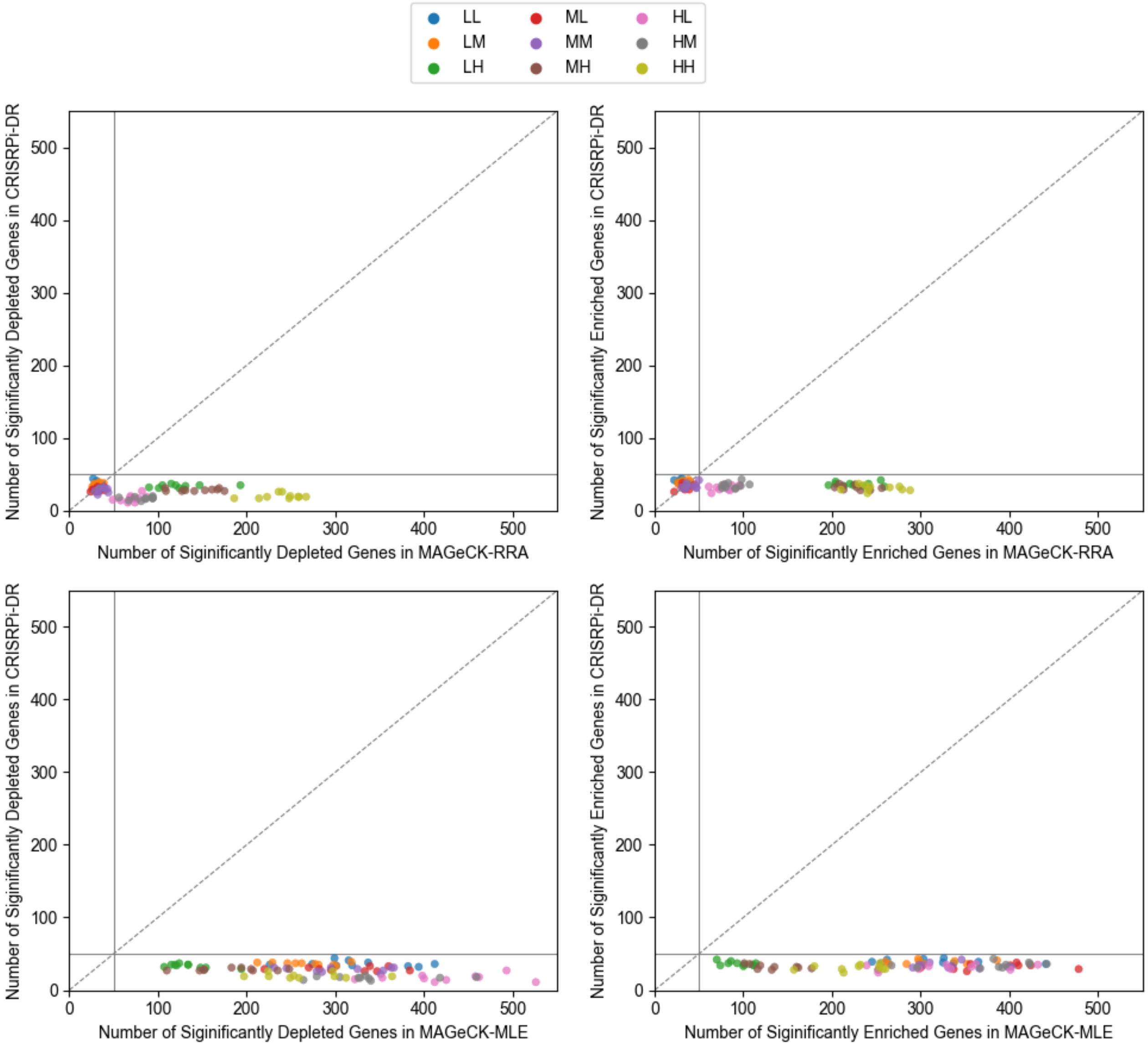


**Fig 2. Comparisons of significant number of depleted and enriched genes reported by the CRISPRi-DR, MAGeCK and MAGeCK-MLE.** The top two panels are comparison of CRIPSRi-DR to MAGeCK and bottom panels are comparison to MAGeCK-MLE. The left panels are comparisons of depleted genes, and the right panels are comparisons of enriched genes. The number of hits (both enriched and depleted) are greater in MAGeCK-MLE than in the CRISPRi-DR model and in some cases MAGeCK-RRA, with the greatest difference coming from HH noise datasets.

The top two panels of Fig 2 show the number of significant hits detected by CRISPRi DR and MAGeCK-RRA, mirroring Fig 4 in the main text. The solid lines represent the 50 true positives (simulated interacting depleted or enriched genes). The dashed diagonal line is a y=x line. Consistent with observations made with the experimental data, the number of genes reported by CRISPRi DR span a shorter range than the number of genes reported by MAGeCK-RRA. However, in this case we can see that the datasets with the highest noise that have the largest discrepancies between the number of hits for both depleted and enriched genes. Thus, we can infer that the experimental datasets seen in Fig 5 with higher discrepancy are those with high noise resulting in a possibly higher number of false positives. Comparatively as seen in the bottom two panels of Fig 2, the discrepancies between CRISPRi-DR and MAGeCK-MLE are high for all datasets, regardless of noise. CRISPRi-DR hits do not exceed the number of simulated interacting genes for both depleted and enriched cases, whereas MAGeCK-MLE nearly always does. Given there were 50 simulated depleted/enriched genes, the up to 300 depleted or enriched calls made by MAGeCK-MLE include many false positives.

Table 1 shows detailed confusion matrices of the calls made by the three methods, using the 50 simulated negative and 50 simulated positive interactions (genes 0-49 and genes 50-99) as the ground truth. Correct predictions are on the descending diagonals, and the off-diagonal entries represent errors (either false positives, FP, or false negatives, FN). The center of these matrices are correctly identified non-interacting genes. This is the largest square in all noise scenarios, for all three methods and is excluded from recall/precision calculations, to focus on predictive performances for depleted and enriched genes.

MAGeCK-MLE identifies a large number of interacting genes for all the simulated datasets and as a result has a consistently high recall rate. For example, in the lowest noise scenario (LL), CRISPRi-DR identified 74% of the simulated interacting genes, MAGeCK-RRA identifies 56.5% and MAGeCK-MLE identifies 99.9% of the genes. However, this also means MAGeCK-MLE makes many calls that are false positives. CRISPRi-DR falsely identifies an average of 1.4 genes, MAGeCK false identifies 3.9 genes and MAGeCK-MLE identified nearly 542.

As noise is increased through adjustments of the $\sigma_C$ or $P_{nb}$ parameters, the number calls made by MAGeCK increase, thus increasing its recall rate to be similar to that of MAGeCK MLE. In the HH scenario, the recall rate of MAGeCK-MLE remains quite high at 88.3% and MAGeCK-RRA's increases to 87.5%. Conversely, with the increased noise, CRISPRi-DR makes fewer calls resulting in a decreased recall rate of 30.1% in the HH scenario. However, CRISPRi-DR sustains a low false positive rate at 2.2%. Whereas the false positive rates of MAGeCK-MLE and MAGeCK increase substantially (MLE = 42.5%, RRA = 42.1%), diluting the sets of predicted enriched and depleted genes with non-interacting genes. In summary, with increasing noise CRISPRi-DR identifies less of the true interacting genes, yet maintains its ability to keep the set of reported interacting genes from being diluted with non-interacting genes.

**Table 1. Evaluation of CRISPRi-DR, MAGeCK-RRA, and MAGeCK-MLE performances in the nine noise scenarios.** For each of the noise scenarios, CRISPRi-DR, MAGeCK-RRA and MAGeCK-MLE are run. These confusion matrices reflect the average confusion matrix of the 10 runs per noise scenario.

| scenario LL | | | CRISPRi-DR predictions | | | MAGeCK-RRA predictions | | | MAGeCK-MLE predictions | | |
|---|---|---|---|---|---|---|---|---|---|---|---|
| | | | Depleted | Non-Interacting | Enriched | Depleted | Non-Interacting | Enriched | Depleted | Non-Interacting | Enriched |
| | Simulated Gene Types | Depleted | 35.8 | 14.2 | 0 | 29.1 | 20.9 | 0 | 49.9 | 0.1 | 0 |
| | | Non-Interacting | 0.5 | 848.6 | 0.9 | 1.3 | 846 | 2.6 | 276 | 308 | 266 |
| | | Enriched | 0 | 11.8 | 38.2 | 0 | 22.6 | 27.4 | 0 | 0 | 50 |

| scenario LM | | | CRISPRi-DR predictions | | | MAGeCK-RRA predictions | | | MAGeCK-MLE predictions | | |
|---|---|---|---|---|---|---|---|---|---|---|---|
| | | | Depleted | Non-Interacting | Enriched | Depleted | Non-Interacting | Enriched | Depleted | Non-Interacting | Enriched |
| | Simulated Gene Types | Depleted | 35.9 | 14.1 | 0 | 31.3 | 18.7 | 0 | 49.8 | 0.2 | 0 |
| | | Non-Interacting | 0.6 | 848.8 | 0.6 | 1.4 | 844 | 4.8 | 218.1 | 373.5 | 258.4 |
| | | Enriched | 0 | 12.5 | 37.5 | 0 | 21.1 | 28.9 | 0 | 0 | 50 |

| scenario LH | | | CRISPRi-DR predictions | | | MAGeCK-RRA predictions | | | MAGeCK-MLE predictions | | |
|---|---|---|---|---|---|---|---|---|---|---|---|
| | | | Depleted | Non-Interacting | Enriched | Depleted | Non-Interacting | Enriched | Depleted | Non-Interacting | Enriched |
| | Simulated Gene Types | Depleted | 32.1 | 17.8 | 0 | 41.3 | 7.6 | 1.1 | 49.3 | 0.7 | 0 |
| | | Non-Interacting | 1.3 | 847.1 | 1.2 | 84 | 585 | 181 | 85.7 | 718.7 | 45.6 |
| | | Enriched | 0 | 14.8 | 35.2 | 0.2 | 6.9 | 42.9 | 0 | 1.3 | 48.7 |

| scenario ML | | | CRISPRi-DR predictions | | | MAGeCK-RRA predictions | | | MAGeCK-MLE predictions | | |
|---|---|---|---|---|---|---|---|---|---|---|---|
| | | | Depleted | Non-Interacting | Enriched | Depleted | Non-Interacting | Enriched | Depleted | Non-Interacting | Enriched |
| | Simulated Gene Types | Depleted | 27.3 | 22.7 | 0 | 29.9 | 20.1 | 0 | 50 | 0 | 0 |
| | | Non-Interacting | 1.7 | 842.4 | 5.9 | 2 | 842 | 5.8 | 262.6 | 257.2 | 330.2 |
| | | Enriched | 0 | 23.6 | 26.4 | 0 | 22 | 28 | 0 | 0.1 | 49.9 |

| scenario MM | | | CRISPRi-DR predictions | | | MAGeCK-RRA predictions | | | MAGeCK-MLE predictions | | |
|---|---|---|---|---|---|---|---|---|---|---|---|
| | | | Depleted | Non-Interacting | Enriched | Depleted | Non-Interacting | Enriched | Depleted | Non-Interacting | Enriched |
| | Simulated Gene Types | Depleted | 26.9 | 23.1 | 0 | 33 | 16.9 | 0.1 | 49.8 | 0.2 | 0 |
| | | Non-Interacting | 1.4 | 842.3 | 6.3 | 3.5 | 839 | 8 | 259.5 | 302.4 | 288.1 |
| | | Enriched | 0 | 22.5 | 27.5 | 0 | 18.6 | 31.4 | 0.1 | 0.1 | 49.8 |

| scenario MH | | | CRISPRi-DR predictions | | | MAGeCK-RRA predictions | | | MAGeCK-MLE predictions | | |
|---|---|---|---|---|---|---|---|---|---|---|---|
| | | | Depleted | Non-Interacting | Enriched | Depleted | Non-Interacting | Enriched | Depleted | Non-Interacting | Enriched |
| | Simulated Gene Types | Depleted | 25.1 | 24.9 | 0 | 43.4 | 5.2 | 1.4 | 47.6 | 2.4 | 0 |
| | | Non-Interacting | 3.3 | 840.4 | 6 | 101 | 567 | 183 | 135.7 | 614.5 | 99.8 |
| | | Enriched | 0 | 24.3 | 25.7 | 0.1 | 7.6 | 42.3 | 0 | 3.7 | 46.3 |

| scenario HL | | | CRISPRi-DR predictions | | | MAGeCK-RRA predictions | | | MAGeCK-MLE predictions | | |
|---|---|---|---|---|---|---|---|---|---|---|---|
| | | | Depleted | Non-Interacting | Enriched | Depleted | Non-Interacting | Enriched | Depleted | Non-Interacting | Enriched |
| | Simulated Gene Types | Depleted | 13.7 | 36.3 | 0 | 33.6 | 16.2 | 0.2 | 48.6 | 0.9 | 0.5 |
| | | Non-Interacting | 3.1 | 831.4 | 15.5 | 37 | 766 | 46.9 | 375.3 | 190.1 | 284.6 |
| | | Enriched | 0 | 34 | 16 | 0 | 17.9 | 32.1 | 1 | 2.8 | 46.2 |

| scenario HM | | | CRISPRi-DR predictions | | | MAGeCK-RRA predictions | | | MAGeCK-MLE predictions | | |
|---|---|---|---|---|---|---|---|---|---|---|---|
| | | | Depleted | Non-Interacting | Enriched | Depleted | Non-Interacting | Enriched | Depleted | Non-Interacting | Enriched |
| | Simulated Gene Types | Depleted | 13.6 | 36.4 | 0 | 35.5 | 14.5 | 0 | 47.9 | 1.6 | 0.5 |
| | | Non-Interacting | 3.3 | 828.1 | 18.6 | 47.1 | 749 | 53.9 | 290.7 | 230 | 329.3 |
| | | Enriched | 0 | 34 | 16 | 0 | 16.1 | 33.9 | 0.5 | 1.8 | 47.7 |

| scenario HH | | | CRISPRi-DR predictions | | | MAGeCK-RRA predictions | | | MAGeCK-MLE predictions | | |
|---|---|---|---|---|---|---|---|---|---|---|---|
| | | | Depleted | Non-Interacting | Enriched | Depleted | Non-Interacting | Enriched | Depleted | Non-Interacting | Enriched |
| | Simulated Gene Types | Depleted | 15 | 35 | 0 | 43.9 | 4.4 | 1.7 | 45.8 | 4.1 | 0.1 |
| | | Non-Interacting | 4.9 | 828.7 | 15.8 | 194 | 451 | 206 | 221.4 | 446.5 | 182.1 |
| | | Enriched | 0 | 34.8 | 15.1 | 0.6 | 5.8 | 43.6 | 0.7 | 6.8 | 42.5 |

To clarify, the metrics are calculated as follows. For depleted genes, $Recall_D=TP_D/(TP_D+FN_D)$ and $Precision_D= TP_D/(TP_D+FP_D)$. For enriched genes, $Recall_E=TP_E/(TP_E+FN_E)$ and $Precision_E= TP_E/(TP_E+FP_E)$. For overall results, depleted and enriched genes are combined as follows: $Recall=(TP_D+TP_E)/(TP_D+FN_D+TP_E+FN_E)$ and $Precision=(TP_D+TP_E)/( TP_D+FP_D+TP_E+FP_E)$. Also, Recall=TPR, Precision=1-FPR.

| | | Predicted Gene Types | | |
|---|---|---|---|---|
| | | Depleted | Non-Interacting | Enriched |
| Simulated Gene Types | Depleted | $TP_D$ | $FN_D$ | |
| | Non-Interacting | $FP_D$ | | $FP_E$ |
| | Enriched | | $FN_E$ | $TP_E$ |

Table 2 is a summary of the average recall, precision, and F1 scores for the three methods across the 9 scenarios. These scores, averaged over positive and negative interactions (excluding non-interactions), reiterate the trend observed in Table 1. As noise increases, recall rates in MAGeCK-RRA and MAGeCK-MLE increase but precision substantially decreases. Conversely, CRISPRi-DR follows a more conservative approach resulting in slightly decreasing recall rate as noise increase but maintaining consistently high precision. Across most of the 9 noise scenarios, CRISPRi-DR has higher F1-scores than the other two methods where $F1\ score = 2 \times \frac{recall \times precision}{recall + precision}$ reflecting the tradeoff between recall and precision.

**Table 2. Average precision and recall values across 10 runs each for CRISPRi-DR, MAGeCK and MAGeCK-MLE for each of the 9 noise scenarios.**

| | CRISPRi-DR | | | MAGeCK-RRA | | | MAGeCK-MLE | | |
|---|---|---|---|---|---|---|---|---|---|
| | *recall* | *precision* | *F1-score* | *recall* | *precision* | *F1-score* | *recall* | *precision* | *F1-score* |
| **LL** | 0.740 | 0.982 | 0.844 | 0.565 | 0.936 | 0.705 | 0.999 | 0.156 | 0.270 |
| **LM** | 0.734 | 0.984 | 0.841 | 0.602 | 0.907 | 0.724 | 0.998 | 0.173 | 0.295 |
| **LH** | 0.674 | 0.964 | 0.793 | 0.853 | 0.242 | 0.377 | 0.980 | 0.429 | 0.597 |
| **ML** | 0.537 | 0.877 | 0.666 | 0.579 | 0.881 | 0.699 | 0.999 | 0.144 | 0.252 |
| **MM** | 0.544 | 0.876 | 0.671 | 0.645 | 0.850 | 0.733 | 0.997 | 0.154 | 0.267 |
| **MH** | 0.508 | 0.845 | 0.635 | 0.870 | 0.232 | 0.366 | 0.939 | 0.286 | 0.438 |
| **HL** | 0.297 | 0.615 | 0.401 | 0.658 | 0.441 | 0.528 | 0.962 | 0.126 | 0.223 |
| **HM** | 0.296 | 0.573 | 0.390 | 0.694 | 0.408 | 0.514 | 0.966 | 0.134 | 0.235 |
| **HH** | 0.301 | 0.594 | 0.400 | 0.895 | 0.180 | 0.300 | 0.890 | 0.180 | 0.299 |

The effect of noise on the results of the three methodologies can also visualized using a bar chart of true and false positives as in Fig 3 (calculated from enriched and depleted genes combined). We calculate the number of significant genes while increasing the amount of noise using either the $\sigma_C$ and $P_{nb}$ parameters (resulting in the 9 noise scenarios). In the left panel, the average number of significant genes were calculated for each method run for a specific $\sigma_C$ value across the possible $P_{nb}$ values. The errorbars seen are the 95% confidence interval of the number of significant genes. For example, the orange bar for $\sigma_C$ =0.01 represents the average number of genes found significant by MAGeCK in the LL, LM, LH scenarios (where $\sigma_C$ =0.01, but $P_{nb}$ varies). The same is done in the right panel, using $P_{nb}$. Noise between concentration increases as $\sigma_C$ is increased and noise between replicates is increases as $P_{nb}$ is decreased.

All three methods make a comparable number of true positive calls, regardless of noise parameters. When noise increases for either parameter, the number of false positive calls for all three methods also increases. However, the number of false positives calls are much higher for MAGeCK-RRA and MAGeCK-MLE than for CRISPRi-DR.

In this plot, it is clear how much more MAGeCK-RRA is affected by noise among replicates than between concentrations. The orange bar for $P_{nb}$=0.1 represents the results for MAGeCK-RRA in the LH, MH, HH scenarios and it shows the highest number of false positives compared to the other $P_{nb}$ values as well as the $\sigma_C$ values, consistent with the observations made in Fig 2. This is likely a result of stochastic fluctuations of counts at individual drug concentrations that are not necessarily supported at other concentrations. This could help explain the poor performance of MAGeCK-RRA on certain datasets that are especially noise, which often generates a large number of hits; our analysis suggests that many of these hits could be false positives. CRISPRi-DR and MAGeCK-MLE are more affected by noise within concentrations than noise within replicates, since these methods rely more on increasing or decreasing trends in abundance that must be (at least somewhat) consistent across concentrations and thus is less affected by within replicate noise as MAGeCK-RRA is.

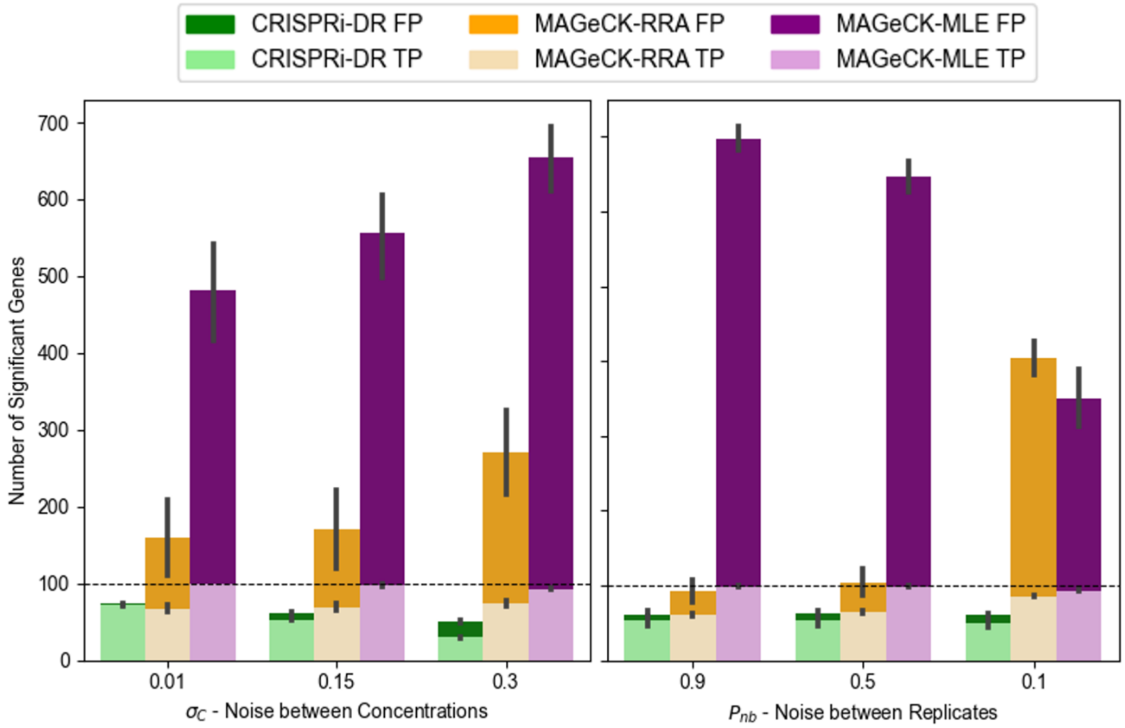


**Fig 3. Bar chart of the average true positives and false positives calls for CRISPRi-DR and MAGeCK-MLE as noise parameters are adjusted to increase noise.** The horizontal dashed line in both panels is the number of total simulated interacting genes (100 total). $\sigma_C$ is increased to increase the noise between concentrations and $P_{nb}$ is decreased to increase noise between replicates of concentrations. The leftmost bars of the plot are the lowest noise, and the rightmost bars are the highest noise.

Effect of $K_2$ (simulated concentration-dependent slopes)

The $K_2$ value in the simulation controls the mean of the interacting slopes in the experiment. The $\sigma_{K2}$, or the standard deviation of the simulation, is always 0.2. Fig 4 shows some of the distributions resulting from select $K_2$ values. As seen in Panel A with $K_2$=0.4, if $K_2$ is low, the slopes distributions of the interacting genes nearly overlap with the distribution of the slopes of the non-interacting genes. This complete overlap makes simulated genes hard to differentiate resulting in many false calls (false positives or false negatives). The other end of the $K_2$ range is seen in Panel C with $K_2$=1.2, where the slopes of non-interacting and interacting genes have almost no overlap. This makes it easy for all methods to identify nearly all the true positives. The value we use in this simulation $K_2$=0.8 is seen in Panel B, that has some overlap between distributions of slopes for interacting and non-interacting genes.

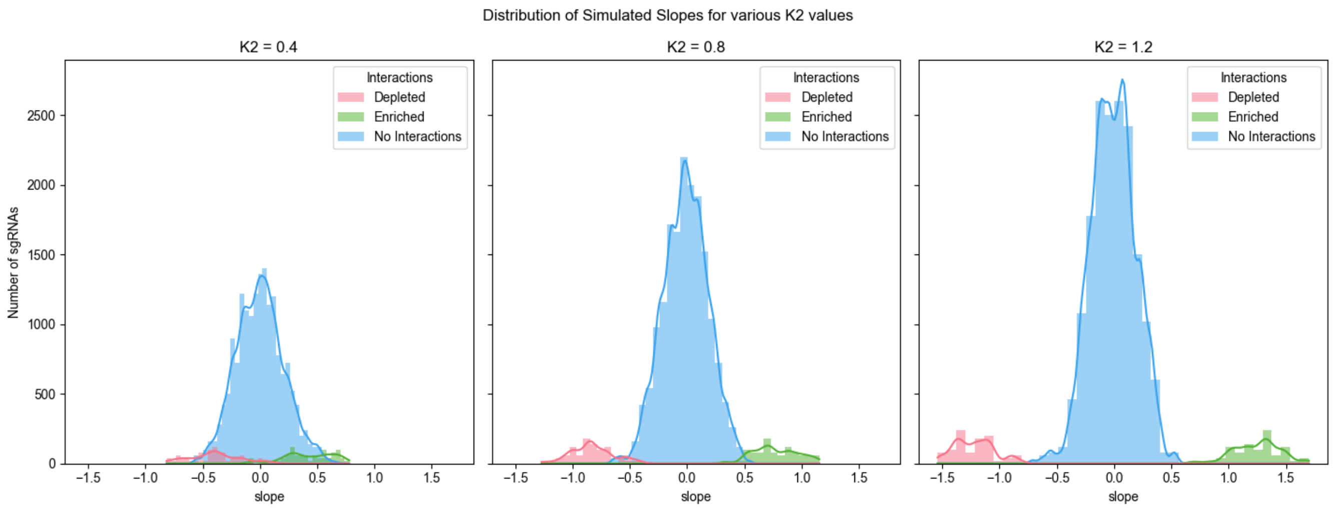


**Fig 4. Distribution of slopes sampled from for simulated non-interacting and interacting genes at multiple $K_2$ values.** The leftmost panel shows the highest amount of overlap of the distributions of simulated slopes with $K_2$ is low and the rightmost panel is the lowest amount of overlap of the distributions of simulated slopes with $K_2$ is high. The middle panel strikes a balance between these two.

Figure 5 shows the true and false positive calls made by CRISPRi-DR, MAGeCK-MLE and MAGeCK-RRA within the lowest noise scenario (LL) for a range of $K_2$ values, from 0.4 to 1.2. As expected, all three methods show increases in true positives and decreases in false positives as $K_2$ increases. Regardless of $K_2$ value, MAGeCK-MLE makes many more calls than CRISPRi-DR and MAGeCK-RRA and thus makes the highest number of true positive calls across the range of $K_2$ values but also makes a highest number of false positives calls. Comparatively, the number of false positive calls made by MAGeCK-RRA and CRISPRi-DR are low regardless of $K_2$ value.

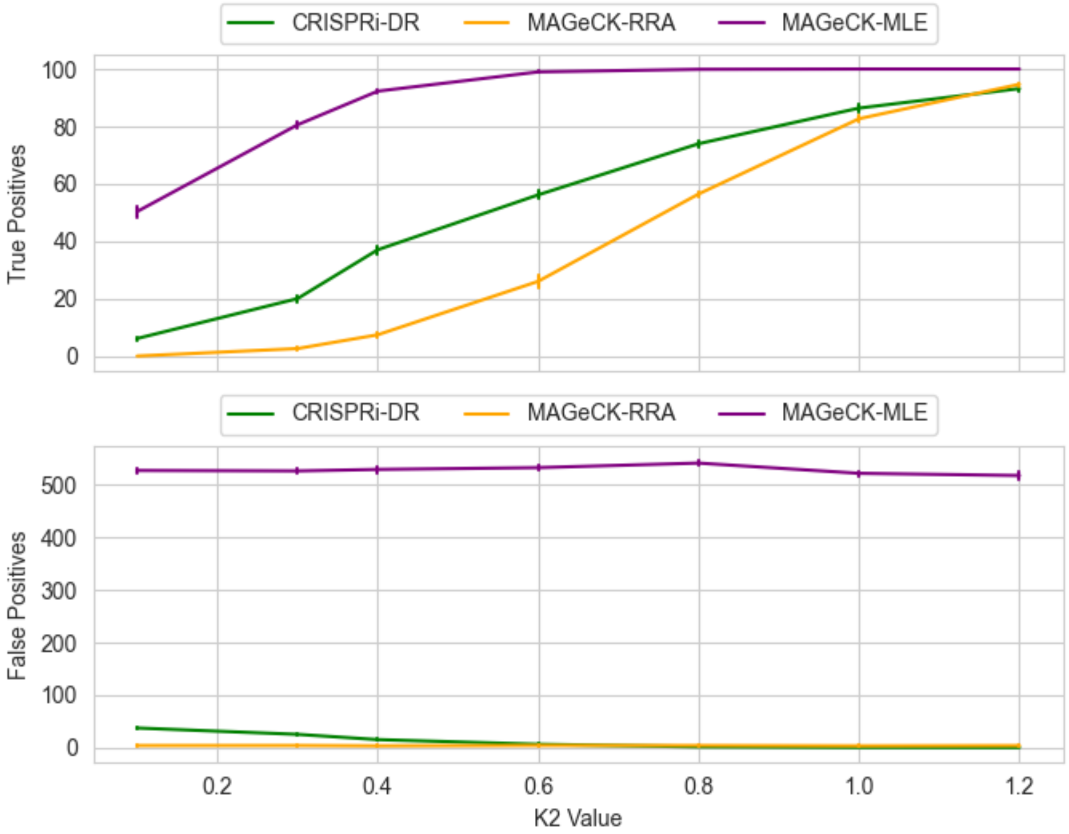


**Fig 5. Calls made by CRISPRi-DR and MAGeCK-MLE at different $K_2$ values.** CRISPRi-DR and MAGeCK-MLE were both run using the LL scenario for 3 total concentrations at a range of $K_2$ values. Each $K_2$ value was run 10 times each.

<u>Effect of Multiple Concentrations on Significant Genes detected</u>

To evaluate the benefit of profiling a CRISPRi library on multiple concentrations on the performance of CRISPRi-DR and the MAGeCK methods, we adapted the simulation above to compare their precision and recall when using one, two, or three drug concentrations. We re-ran the simulation (10 iterations per concentration amount), keeping the same concentration range (2-8 μM), and used the high-noise (HH) parameter settings ($\sigma_C$ =0.1, $P_{nb}$ =0.1). In all runs, we always kept the highest concentration (8 μM), along with the no-drug control. First, we evaluated performance using only one concentration (the highest, 8 μM), then the two highest (4 and 8 μM), then all three concentrations (the full range, 2-8 μM). For MAGeCK-RRA, each concentration was compared independently to the no-drug control, and then a single LFC value was calculated per gene as the most significant LFC across the concentrations and a single P-value was calculated per gene using the Fisher's method, which was then adjusted with Benjamini-Hochberg. For CRISPRi-DR and MAGeCK-MLE, the no-drug control was treated as the lowest concentration, and it was combined with either the highest concentration, top two, or all three drug concentrations, which were then used for doing regressions (over 2-4 effective concentration points).
As seen in Fig 6, the recall of CRISRi-DR and MAGECK-MLE (number of true positives) held constant as concentrations were added, whereas the number of true positives identified by MAGeCK-RRA increased

slightly with more concentrations. Adding concentration points caused a significant increase in the number of false positives found by of MAGeCK-RRA. This is apparently due to the commitment of false positive errors, i.e. non-interacting simulated genes that are classified as significant due to high variation in barcode counts in this high noise setting. MAGeCK is susceptible to false positives when evaluating only a single concentration point, but this gets amplified as more concentrations are added, because each concentration is evaluated independently, and the hits (including false positive genes) are combined post-analysis, explaining why precision drops as concentrations are added, because errors accumulate.

In contrast, CRISPRi-DR and MAGeCK-MLE are more robust with respect to false-positive errors, because regression incorporates data from all concentrations available and looks for significant trends. This allows CRISPRi-DR to maintain higher precision, which does not decrease as additional concentration points are added.

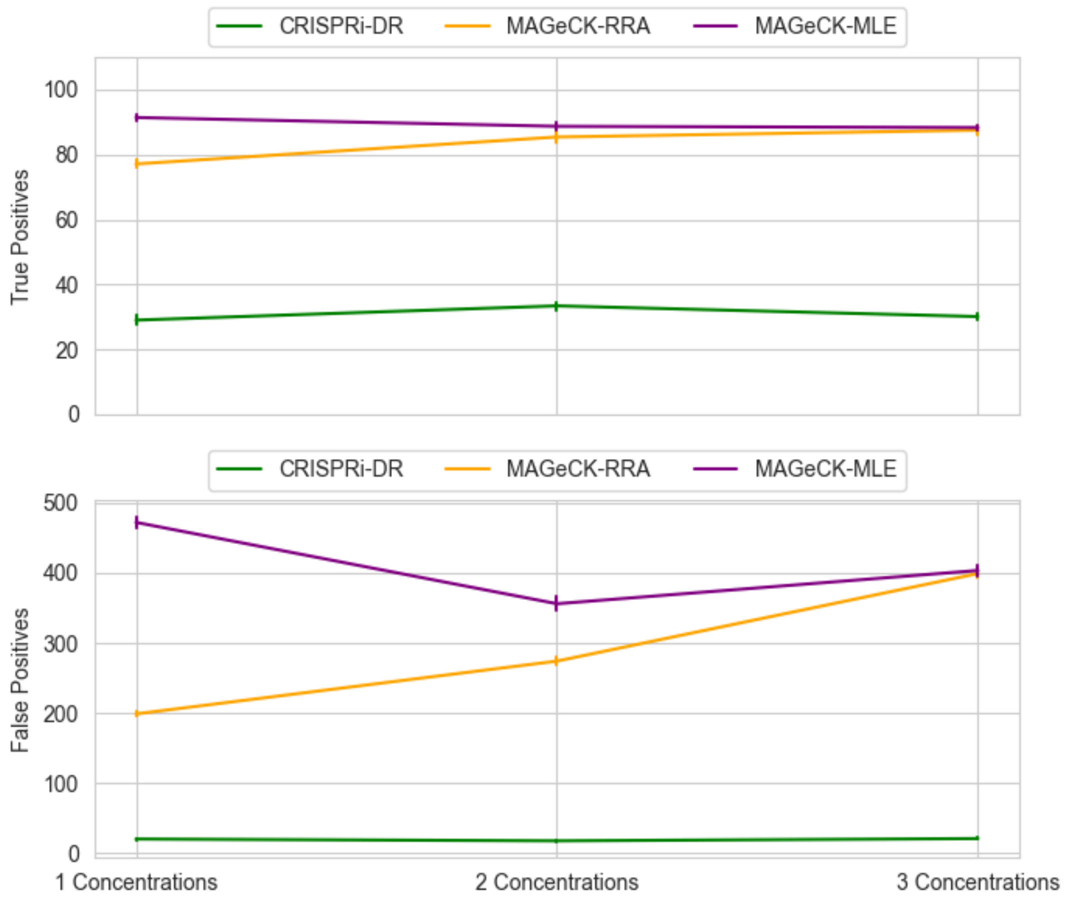


**Fig 6. Changes in true and false positives with number of concentrations.** CRISPRi-DR, MAGeCK-RRA, MAGeCK-MLE were all run using the HH scenario for 1, 2 and 3 total concentrations. Each concentration was run 10 times, i.e. on 10 independent simulated datasets.

<u>CRISPRi-DR can detect interacting genes where trends are not perfectly linear</u>

It is not always the case that the highest concentration should be the most informative one for detecting CGIs, as it might cause too much growth inhibition, making it difficult to assess dose-dependent behavior.

Sometimes the largest effect occurs at the edge of the range, like dropping off a cliff, due to uncertainty about the optimal concentration. CRISPRi-DR can detect these kinds of trends. It only expects abundances to show an overall trend depletion or enrichment across concentrations. Fig 6 shows a few sgRNAs depicting these kinds of trends for representative sgRNAs targeting simulated interacting genes detected as significant by CRISPRi-DR. sgRNA 1 shows an example of an ideal (nearly linear) pattern of depletion. Comparatively, sgRNA 2 shows a sharp decline in abundance at concentration 1 (the lowest end of the concentration range), sgRNA 3 shows a sharp decline at concentration 2 and sgRNA 4 shows a sharp decline at concentration 3 (the highest end of the concentration range). Regardless of depletion pattern, sgRNAs 1, 2, 3 and 4 all target genes detected as significant negative interactions by the CRISPRi-DR model.

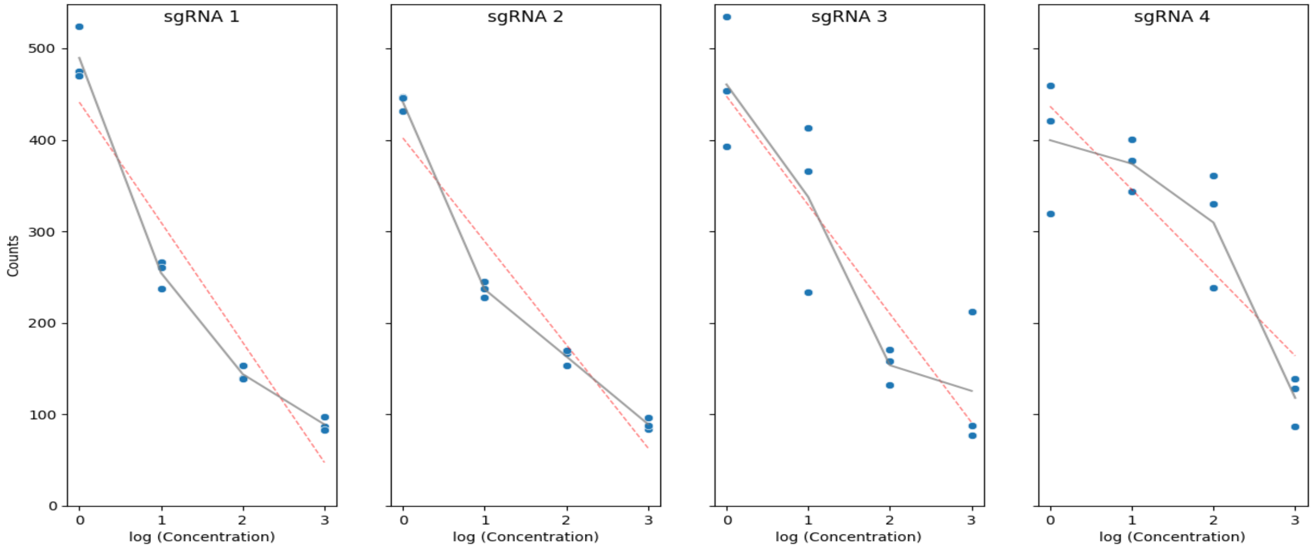


**Fig 6. Trends of select sgRNAs in the various noise scenario.** For each of the sgRNAs seen here, the blue dots are abundances for the three simulated replicates at each concentration. The gray line shows the change in abundance mean as concentration increases.

### 2. Comparison of CRISPRi-DR to other CRISPR analysis methods using CGI datasets

#### Algorithms Summary

The performance of CRISPRi-DR is compared to six other software packages intended for analyzing CRISPRi and CRISPRko libraries.

Below are general overviews of some existing methodologies comparable to CRISPRi-DR used to assess CRISPR screens. We describe the general math used in the analyses, as well as how we adjusted our datasets to be run using each methodology. Some methods work directly on counts, whereas others require log fold changes. The result of some of these methods is at the sgRNA level, which were combined post hoc to obtain gene level information. Additionally, some of these methods do not take multiple concentrations into account, thus we combined outputs of analyses at different concentrations using Fisher's method of combing P-values:

$$p_i = \sum_{c=0}^{n} \log p_{ic} = \log(p_{i_{LOW}}) + \log(p_{i_{MED}}) + \log(p_{i_{HIGH}})$$

where n=3 for the 3 concentration levels (low, medium, high) for each gene $i$.

DEBRA (DESeq2-based RNA-seq Analysis)

R Function Call:

```
> DEBRA(counts, c("DMSO1","DMSO2", "DMSO3"), condition_names=
  c("DRUG1","DRUG2", "DRUG3"), method="DESeq2(Wald)")
```

DEBRA [3] is a methodology that uses the DESeq2 package for RNA sequencing data analysis. It processes RNA-seq count data, normalizing to account for variations in sequencing depth and biases. Then, using Negative Binomial models, DEBRA estimates gene expression variability. Through differential analysis, it identifies genes with statistically significant barcode count changes across different conditions, employing DESeq2's robust testing methods.

To use CRISPRi datasets as inputs to DEBRA, counts of the most efficient sgRNA for each gene were provided as input to DEBRA. The output of DEBRA for these datasets is log fold changes along with P-values calculated using the Ward method. Since DEBRA does not account for increasing concentration, each concentration (low, medium and high) in a given dataset was run separately, with the corresponding DMSO condition set as the control. These gene-wise P-values of the three concentration outputs were combined using the Fisher's method and then adjusted using Benjamini-Hochberg. Genes were ranked by combined P-value and marked significant if adjusted combined P-value < 0.05.

CGA-LMM

Command Line Prompt:

```
> Rscript ./CGA_LMM.R single_sgRNA_per_gene_counts.txt CGA_LMM_out
```

Designed for hypomorph libraries, CGA-LMM [4] assesses the concentration-dependent variation in mutant abundance (in a library) using slope coefficients derived from linear mixed models. CGA-LMM assumes one set of counts per gene and was not designed to incorporate information from multiple sgRNAs. This method uses a conservative population-based approach by identifying genes as significant only if they have slopes that are outliers when compared to the general population.

To use CRISPRi data as the input for this method, we use the most efficient sgRNA for each gene. In line with the approach of Dutta, DeJesus (4), significant genes were marked as those with adjusted P-value< 0.5 and |Zrobust| > 3.5. Genes were ranked by P-value.

CRISPhieRmix

R Function Call:

```
> CRISPhieRmix(log2fc_from_DESeq2, geneIds, negCtrl, nMesh = 100,
  PLOT = TRUE, VERBOSE = TRUE, mu=-6, pq=0.1,BIMODAL=T)
```

CRISPhieRmix [5] is a methodology created specifically for the CRISPRi variant to address guide efficiency present in CRISPRi and CRISPRa screens. The method takes log fold changes of sgRNA counts between two conditions, typically derived from DESeq2 or edgeR outputs. It fits a hierarchical mixture model on these log fold changes, based on the assumption that genes are represented by a mixture distribution of effective and ineffective guides. CRISPhieRmix then computes False Discovery Rates (FDRs) as the posterior probability that a gene is non-essential. These probabilities are aggregated over all possible mixtures to finalize the FDRs for each gene.

In analyzing our CRISPRi datasets, we utilized the *bimodal* option available in the method's implementation. Typically, this analysis resulted in one positive and one negative mode, consistent with the option's assumption that genes are a mixture of ineffective, depleting, or enriching guides. Since CRISPhieRmix does not account for increasing concentration, each concentration (low, medium, and high) in each dataset was run separately, with the corresponding DMSO condition set as the control. In the actual counts files, negative control genes were excluded, as they were causing noise in the mixture model estimations. The local FDR s from the analysis of the three concentrations were combined using the Fisher's method. Genes were ranked by were combined local FDR and marked significant if combined local FDR < 0.05.

MAGeCK (Model-based Analysis of Genome-wide CRISPR-Cas9 Knockout)

Command Line Prompt:

```
> mageck test -k drug_counts.txt -c DMSO1,DMSO2,DMSO3 -t
  CONC1,CONC2,CON3 -n Mag_out --gene-lfc-method alphamedian --norm-
  method control --control-sgrna negatives.txt
```

Li, Xu (6) designed the Robust Ranking Algorithm (RRA), one of the first algorithms for CRISPRko screens, available to researchers as MAGeCK [7]. The input to the method is raw control and experimental sgRNA counts. These counts are fitted to a Negative Binomial model to assess if counts vary significantly (similar to DESeq2). The sgRNA level P-values for each gene are the combined using a modified version of robust rank aggregation to evaluate whether a subset of them is enriched (RRA), resulting in a list of genes with False Discovery Rates (FDRs) for both positive and negative interactions.

Since MAGeCK does not account for multiple concentrations, each concentration (low, medium, and high) in a given dataset was run separately, with the corresponding DMSO condition set as the control. MAGeCK utilizes a separate set of controls be provided, from which it calculates its P-values. We provided

"Negative" sgRNA controls in list form to the method for this purpose (using 1750 non-targeting sgRNAs in the Mtb CRISPRi library). For each concentration, we determined a gene's overall P-values based on the lowest FDR, whether positive or negative. We then merged the resulting gene P-values from the three concentrations using the Fisher's method and then adjusted using Benjamini-Hochberg. Genes were ranked by combined P-value and marked significant if adjusted combined P-value < 0.05.

Direct comparisons of the significant genes in the CGI libraries from Li, Poulton (1) found by CRISPRi-DR and MAGeCK-RRA can be seen in Supplemental Table S3, where significant calls made by MAGeCK-RRA have an additional constraint of |LFC|>1, to be more comparable to the publication's analysis of the data.

<u>DrugZ</u>

Command Line Prompt:

```
> drugz.py -i drug_counts.txt -o drugZ_out.txt -c DMSO1,DMSO2,DMSO3
  -x CONC1,CONC2,CONC3
```

DrugZ [8] is a method for analyzing chemical genetic interactions with drug treatments in CRISPRi, CRISPRko and CRIPSRa libraries treated with drugs. Raw sgRNA counts of the control and experimental condition are provided as input to the method. The log fold changes of the normalized counts are then calculated, along with guide level z-scores and variance estimated by empirical Bayes. The z-scores at the guide level are summed to get gene level z-scores, from which P-values can be obtained from a Normal distribution. The output of the method is similar to MAGeCK in that it provides the statistics for both the suppressive and synergistic interactions of the genes.

In our approach to analyzing CRISPRi datasets with drugZ, sgRNA counts were used directly without modification. Similar to many other methodologies, drugZ does not simultaneously accommodate multiple concentrations. Thus, each concentration (low, medium, and high) in a given dataset was run separately, with the corresponding DMSO condition set as the control. At each concentration, we determined a gene's overall P-value based on the lowest P-value, whether suppressive or synergistic. We then merged the resulting gene P-values across the three concentrations using the Fisher's method and then adjusted using Benjamini-Hochberg. Genes were ranked by were combined P-value and marked significant if adjusted combined P-value < 0.05.

<u>MAGeCK MLE</u>

Command Line Prompt:

```
> mageck mle --norm-method control -n MLE_out --genes-var 0 --
  update-efficiency --count-table subsampled_drug_counts.txt --
  threads 16 --control-sgrna negatives.txt --design-matrix
  design_matrix.txt --sgrna-efficiency squashed_sgRNA_info.txt --
  sgrna-eff-name-column 1 --sgrna-eff-score-column 2
```

MAGeCK MLE [9], an extension of MAGeCK, estimates gene effects across multiple conditions (i.e. cell lines or drug treatments), while accounting for sgRNA knockout efficiency. Like MAGeCK, the input to the method is a set of raw sgRNA counts but also requires a design matrix specifying which counts come from which condition along with sgRNA efficiencies between the range of 0 and 1. Raw sgRNA counts are fitted to a Negative Binomial GLM with log link to sgRNA level counts. Maximum likelihood estimation (MLE) of fitting the guide counts across all samples is used to calculate the beta scores. The significance of these beta scores is calculated through the Wald test.

When using MAGeCK MLE with default settings, we found that a maximum of 39 sgRNAs per gene could be processed without triggering "gene too large" errors. Therefore, for genes with more than 39 sgRNAs in our CRISPRi datasets, we randomly selected 39 sgRNAs for analysis. In our design matrices, we treated increasing concentration as a time series variable. Additionally, we included sgRNA efficiencies, expressed as estimated log fold change values normalized to a 0-1 scale, where 1 represents higher sgRNA efficiency. Like MAGeCK, MAGeCK-MLE requires a list of control sgRNAs which we fulfilled with a list of "Negative" sgRNAs in each CRISPRi dataset. Genes in the MAGeCK MLE analysis results were ranked by Wald P-value and significant genes were marked as those with Wald FDR < 0.05.

**Combining Results from Multiple Concentrations using Fisher's Method for P-values**

In our study, for methods that weren't designed to handle multiple concentrations simultaneously, we ran each method three times for a given CRISPRi drug dataset, once for each concentration level: low, medium, and high. We combined the significance of the genes across the three analysis results using Fisher's method. Fig 7 illustrates the impact of this combined approach. The ROC Curves are for RIF in 1 day pre-depletion, setting target genes as the 75 conditionally essential genes (adjusted P-value < 0.5 from resampling in Transit) from a previously published TnSeq study of *M. tuberculosis H37Rv* exposed to sub-MIC concentrations of various antibiotics, including rifampicin [10]. While changes in essentiality due to transposon insertions are not technically the same as fitness defects resulting from CRISPRi depletion, there is substantial overlap between essentiality and vulnerability [2].

Panel A displays ROC Curves for all runs of the tested methods, with gene rankings determined by respective P-values. For CRISPhieRmix, we used "locFDR" to rank the genes and for MAGeCK and drugZ, which provide statistics for both positive and negative interactions, we selected the minimum P-value for each gene to rank them. In this case, the high concentration runs performed the best for methods that did not account for multiple concentrations simultaneously. CRISPRi-DR performs among the best (black curve, panel A), partly because it utilizes info from all 3 concentrations. However, one cannot just consider the highest concentration in this analysis, as sometimes the MIC of a drug can be unknown, where the highest concentration might be excessively strong, potentially leading to an overestimation of depletion effects and false positives compared to the control. In Panel B, the ROC Curves after application of the Fisher's method closely resembled, but were not identical to, the high concentration curves in Panel A. This indicates that lower concentrations also contribute to a gene's overall significance. Using the Fisher's method to combine significance of the three concentrations provides a more comprehensive representation of the overall significance of gene interactions across different concentration levels. Nonetheless, CRISPRi-DR still has a ROC curve that is competitive with the best of these other analysis methods (Panel B).

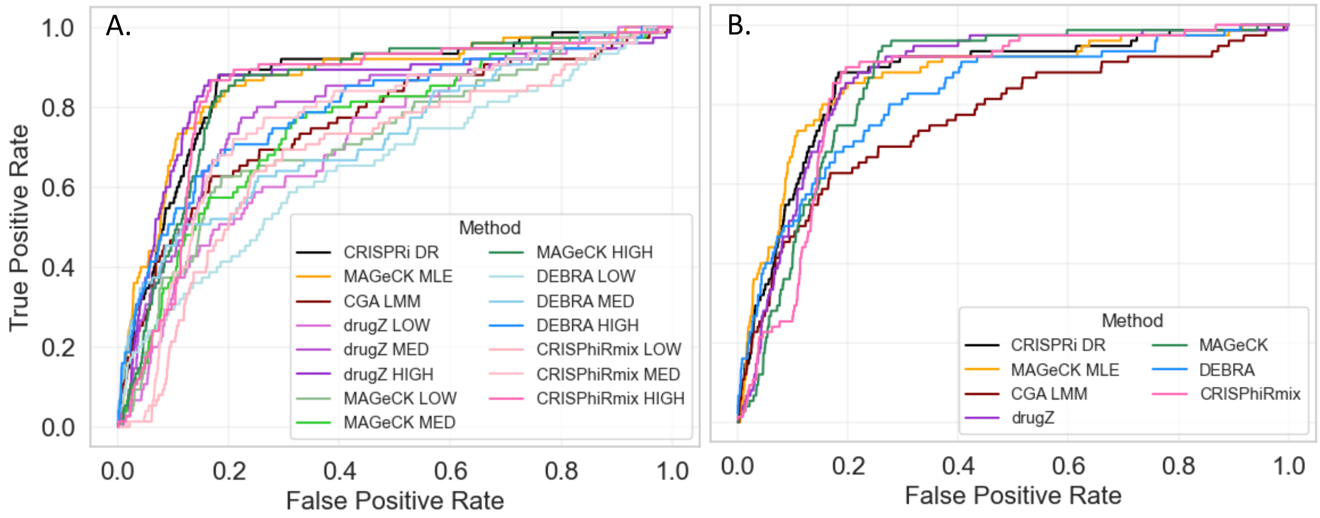


**Fig 7. ROC Curves of RIF in 1 day pre-depletion using known target genes from TnSeq screens as true positives**. A) ROC curves per run of a tested methodology. Rankings based on gene P-values calculated by the methodology, or with minimal post-processing. B) One ROC Curve per method based on ranking of final P-values post combining multiple concentrations for methods that do not account for them.

**Results**

Analysis of RIF D1 CRISPRi dataset by CRISPR Methods

CRISPRi-DR performs as well as the best of the other methods in identifying the target genes by significance-based ranking (ROC curves). While the highest AUC (0.866) is achieved by MAGeCK-MLE on the RIF D1 dataset, CRISRPi-DR has similar AUC of 0.850). However, the number of significant genes both by adjusted P-value < 0.05 per concentration and Fisher's combined adjusted P-value < 0.05 for the methods necessary show much higher number of false positives. The methods that find nearly all the target genes are CRISPhieRmix, MAGeCK, and MAGeCK MLE, but these methods report a very high number of false positives (thousands of genes that are putatively statistically significant). Although CRISPRi-DR and drugZ are the two methods that report a relatively lower number of total hits, they both detect only ~40 of the 75 target genes as significant, and drugZ reports many more false positives (433 overall). Overall, the incorporation of both sgRNA efficiency and increasing concentration in a model such as CRISPRi-DR along with the additional Z-score constraint allows for rankings similar to the other CRISPR analysis techniques, but it reduces the number of false positives reported. Thus, CRISPRi-DR has the highest F1-score of the methodologies tested.

**Table 3. Number of significant genes reported and AUC values of ROC Curves of RIF in 1 day pre-depletion using known target genes from TnSeq screens as true positives**

| Method | AUC Value | Total Number of Significant Genes | True Positives | False Positives | F1-score |
|---|---|---|---|---|---|

| | | | | | |
|---|---|---|---|---|---|
| **CGA-LMM** | 0.765 | 328 | 23 | 305 | 0.114 |
| **CRISPRi-DR** | 0.850 | 183 | 42 | 141 | 0.326 |
| **CRISPhieRmix [Overall]** | 0.844 | 3146 | 74 | 3072 | 0.046 |
| **CRISPhieRmix LOW** | 0.689 | 781 | 14 | 767 | 0.039 |
| **CRISPhieRmix MED** | 0.767 | 869 | 15 | 854 | 0.033 |
| **CRISPhieRmix HIGH** | 0.836 | 2025 | 41 | 1984 | 0.032 |
| **DEBRA [Overall]** | 0.822 | 3063 | 72 | 2991 | 0.046 |
| **DEBRA LOW** | 0.657 | 103 | 13 | 90 | 0.146 |
| **DEBRA MED** | 0.731 | 495 | 37 | 458 | 0.130 |
| **DEBRA HIGH** | 0.798 | 1960 | 65 | 1895 | 0.064 |
| **MAGeCK [Overall]** | 0.855 | 3899 | 75 | 3824 | 0.106 |
| **MAGeCK LOW** | 0.722 | 455 | 28 | 427 | 0.081 |
| **MAGeCK MED** | 0.756 | 1087 | 47 | 1040 | 0.080 |
| **MAGeCK HIGH** | 0.848 | 1634 | 68 | 1566 | 0.038 |
| **MAGeCK MLE** | 0.866 | 2833 | 72 | 2761 | 0.050 |
| **drugZ [Overall]** | 0.867 | 473 | 40 | 433 | 0.146 |
| **drugZ LOW** | 0.716 | 55 | 2 | 53 | 0.064 |
| **drugZ MED** | 0.798 | 60 | 4 | 56 | 0.059 |
| **drugZ HIGH** | 0.847 | 82 | 5 | 77 | 0.031 |

### Sensitivity of CRISPR methods to noise

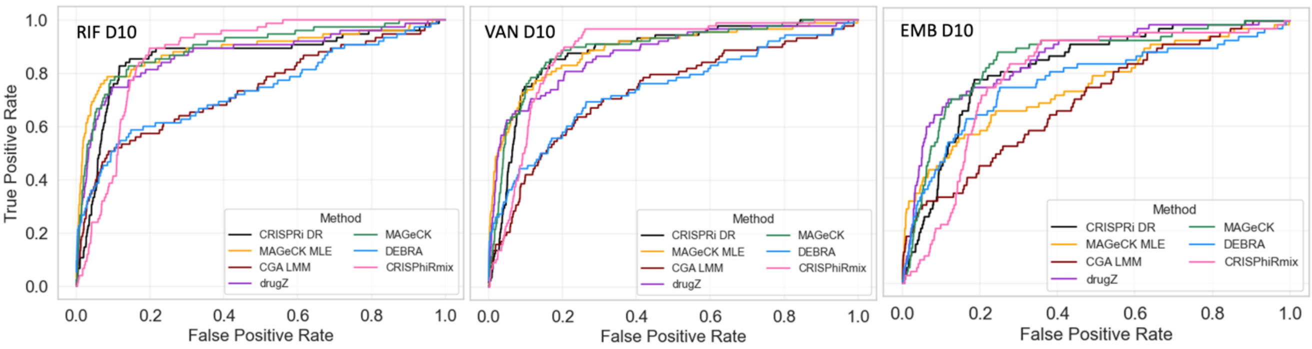


**Fig 8. ROC Curves of RIF, VAN and EMB in 10 day pre-depletion using known target genes from TnSeq screens as true positives**.

We assessed the performance of the methods on 10 day pre-depletion (D10) CRISPRi datasets. With greater pre-depletion days, there is expectedly more noise in the datasets. We compare the methodologies' sensitivity to higher noise through the RIF, VAN and EMB datasets with 10-day pre-depletion, using conditionally essential genes found by Xu, DeJesus (10) in their TnSeq study of antibiotic exposure as target genes. The ROC Curves seen in Fig 8 were generated using the ranking methods mentioned previously. Per AUCs, the performance of CRISPRi-DR remains consistently high for all three datasets. Even in high depletion datasets, CRISPRi-DR identified target genes as well as if not better than the other methodologies tested. Although CRISPRi-DR identified less of the target genes as significant, it marked less false positive genes than the other methods and thus overall had the highest F1 scores.

**Table 4. Number of significant genes reported and AUC values of ROC Curves of RIF, VAN and EMB in 10 day pre-depletion using known target genes from TnSeq screens as true positives (75 genes for RIF, 89 for VAN, and 67 for EMB).**

| ***RIF D10*** | | | | | |
|---|---|---|---|---|---|
| ***Method*** | **AUC Value** | **Total Number of Significant Genes** | **True Positives** | **False Positives** | **F1 Score** |
| *CGA-LMM* | 0.738 | 371 | 25 | 346 | 0.112 |
| *CRISPRi-DR* | 0.860 | 218 | 51 | 167 | 0.348 |
| *CRISPhieRmix* | 0.860 | 3665 | 74 | 3591 | 0.040 |
| *DEBRA* | 0.735 | 2125 | 57 | 2068 | 0.052 |
| *MAGeCK* | 0.892 | 3596 | 74 | 3522 | 0.040 |
| *MAGeCK-MLE* | 0.888 | 2394 | 69 | 2325 | 0.056 |
| *drugZ* | 0.866 | 436 | 54 | 382 | 0.211 |
| ***VAN D10*** | | | | | |
| ***Method*** | **AUC Value** | **Total Number of Significant Genes** | **True Positives** | **False Positives** | **F1 Score** |
| *CGA-LMM* | 0.733 | 514 | 27 | 487 | 0.090 |
| *CRISPRi-DR* | 0.883 | 185 | 50 | 135 | 0.365 |
| *CRISPhieRmix* | 0.865 | 3611 | 89 | 3522 | 0.048 |
| *DEBRA* | 0.741 | 2817 | 77 | 2740 | 0.053 |
| *MAGeCK* | 0.880 | 2849 | 87 | 2762 | 0.059 |
| *MAGeCK-MLE* | 0.889 | 2753 | 86 | 2667 | 0.061 |
| *drugZ* | 0.871 | 408 | 59 | 349 | 0.237 |
| ***EMB D10*** | | | | | |
| ***Method*** | **AUC Value** | **Total Number of Significant Genes** | **True Positives** | **False Positives** | **F1 Score** |
| *CGA-LMM* | 0.622 | 522 | 17 | 505 | 0.058 |
| *CRISPRi-DR* | 0.821 | 172 | 22 | 150 | 0.184 |
| *CRISPhieRmix* | 0.789 | 3342 | 67 | 3275 | 0.039 |
| *DEBRA* | 0.767 | 2423 | 51 | 2372 | 0.041 |
| *MAGeCK* | 0.853 | 3913 | 67 | 3846 | 0.034 |
| *MAGeCK-MLE* | 0.750 | 3142 | 62 | 3080 | 0.039 |
| *drugZ* | 0.857 | 549 | 47 | 502 | 0.153 |

CRISPRi-DR and CGA-LMM both show higher precision than the other methods. However, CRISPRi-DR has a much higher precision than CGA-LMM, perhaps attributed to its ability to utilize both drug concentration and sgRNA efficiency in the model, whereas CGA-LMM only uses concentration. As seen in Table 5, CRISPRi-DR is the best method based on precision and F1-score for nearly all the datasets. There are quite a few methods that show high recall in the datasets (these are usually 100% recall). Precision here is calculated as TP/(FP+TP) and Recall is calculated as TP/(TP+FN). Based on AUC, MAGeCK and MAGeCK-MLE seems to have the highest values. However, Supplemental Table 2 shows that for certain datasets, such as RIF D5, the AUC values are comparably high for most methods.

**Table 5. Assessment of Best CRISPR Method for the EMB, INH, LEVO, VAN and RIF CRISPRi screens in Day 1, 5 and 10 day pre-depletion based on recall, precision, AUC and F1-scrore.** For the methods that require individual dosage runs, they were combined using Fisher's method across concentrations and then assessed. CRISPRi-DR is the best method by F1-Score for most of the drug-treated datasets.

| *Drug* | Days Pre-depletion | Best Method(s) by Recall | Best Method by Precision | Best Method by AUC | Best Method by F1-Score |
|---|---|---|---|---|---|
| *EMB* | 1 | MAGeCK | CRISPRi-DR | drugZ | CRISPRi DR |
| *EMB* | 5 | MAGeCK | CRISPRi-DR | MAGeCK | drugZ |
| *EMB* | 10 | CRISPhieRmix, MAGeCK | CRISPRi-DR | drugZ | CRISPRi DR |
| *INH* | 1 | MAGeCK | CRISPRi-DR | MAGeCK | CRISPRi-DR |
| *INH* | 5 | MAGeCK | CRISPRi-DR | MAGeCK | CRISPRi-DR |
| *INH* | 10 | MAGeCK | CRISPRi-DR | MAGeCK | CRISPRi-DR |
| *RIF* | 1 | MAGeCK | CRISPRi-DR | drugZ | CRISPRi-DR |
| *RIF* | 5 | CRISPhieRmix,MAGeCK, MAGeCK MLE | CRISPRi-DR | MAGeCK MLE | CRISPRi-DR |
| *RIF* | 10 | CRISPhieRmix, MAGeCK | CRISPRi-DR | MAGeCK | CRISPRi-DR |
| *VAN* | 1 | MAGeCK | CRISPRi-DR | MAGeCK MLE | CRISPRi-DR |
| *VAN* | 5 | CRISPhieRmix, MAGeCK | CRISPRi-DR | MAGeCK MLE | CRISPRi-DR |
| *VAN* | 10 | CRISPhieRmix | CRISPRi-DR | MAGeCK MLE | CRISPRi-DR |
| *LEVO* | 1 | DEBRA, MAGeCK | CRISPRi-DR | DEBRA | CRISPRi-DR |
| *LEVO* | 5 | DEBRA | CRISPRi-DR | DEBRA | CRISPRi-DR |
| *LEVO* | 10 | MAGeCK | CRISPRi-DR | DEBRA | CRISPRi-DR |

### 3. Analysis of *E. coli* CRISPRi screen using CRISPRi-DR

#### *E. coli* CRISRPi Dataset for growth on different carbon sources

Mathis, Otto and Reynolds (11) quantified growth rate changes as a function of varied gene expression levels using CRISPRi through a library of modified single guide RNAs (sgRNAs). The dataset consisted of a library of 5927 sgRNAs targeting 88 genes in *Escherichia coli MG1655*, for which they observed their effects on growth rate on media with different carbon sources. To generate diversity, they incrementally added mutations to sgRNAs in the targeting sequence or position in gene and created showed a link between the number of mutations and their impact on growth rate.

The authors demonstrated that many gene-environment interactions not detected at maximum knockdown levels are seen at intermediate levels of expression interference. The authors quantified growth rate effects of modified sgRNAs in different carbon sources (glucose and glycerol) under turbidostat growth conditions. Compared to the conventional method of using a single (maximal-efficiency) knockdown per gene, Mathis, Otto and Reynolds (11) found 37% more interacting genes assessing the differences in the fitted parameters of a logistic fit. This fit included the quantified growth rates and the Hill coefficient.

While this is not technically a chemical-genetics (CGI) experiment, the data included multiple time points along, with sgRNAs designed to span efficiencies, satisfying requirements for our model. Thus, the abundances in this dataset can be represented through a modified version of Eq. (3) in the main text:

$$A_{ijk} = \frac{1}{1+\left(\frac{T_{1/2}}{Time_j}\right)^{H_d}\left(\frac{K_s}{Growth\ Rate_i}\right)^{H_s}}$$

where time replaces log concentration, and growth rate replaces sgRNA efficiency. After logsigmoid transformation, the equation becomes:

$$\log\left(\frac{A_{ijk}}{1-A_{ijk}}\right) = H_T \cdot Time_j + H_s \cdot S_i + C$$
$$C = H_s \cdot \log(K_s) - H_d \cdot T_{1/2}$$

Where the intercept folds in the inflection points $T_{1/2}$ (time results in 50% depletion) and $K_s$ (growth rate results in 50% depletion).

Therefore two linear regressions (one for glucose and one glycerol) we fit for the dataset are :

$$\log\left(\frac{A_{ijk_{Glucose}}}{1-A_{ijk_{Glucose}}}\right) = \beta_0 + \beta_{T_{Glucose}} \cdot Time_j + \beta_{s_{Glucose}} \cdot Growth\ Rate_i$$
$$\log\left(\frac{A_{ijk_{Glycerol}}}{1-A_{ijk_{Glycerol}}}\right) = \beta_0 + \beta_{T_{Glycerol}} \cdot Time_j + \beta_{s_{Glycerol}} \cdot Growth\ Rate_i$$

analogous to Eq (5) in the main text for fitting sgRNA abundances for chemical genetic interaction screens.

As seen in Fig 9, the growth curves of CRISPRi knock-down mutants (depletion over time: 7 timepoints for glucose, 5 for glycerol) follows sigmoidal behavior very analogous to dose-response curves for antibiotic exposure (depletion with increasing concentration), making it a good dataset to analyze with CRISPRi-DR. In Fig 9, the changes in abundance for *fbaA* in glucose and glycerol over time can be seen. *fbaA* is a gene involved in glycolysis but not gluconeogenesis, thus being essential only for growth on glucose. The sigmoidal curve in leftmost panel is similar to the curves seen in Fig 2 of the main text. After applying the logsigmoid transformation, both sets of curves become linear, and we can assess the time-based depletion of the gene. In glycerol, there is minimal interaction of the gene with the carbon source and thus the slope of time-based dependence post-logsigmoid transformation is slightly positive; hence, depleting *fbaA* does not produce a growth defect for growth on glycerol, in contrast to what is seen for glucose.

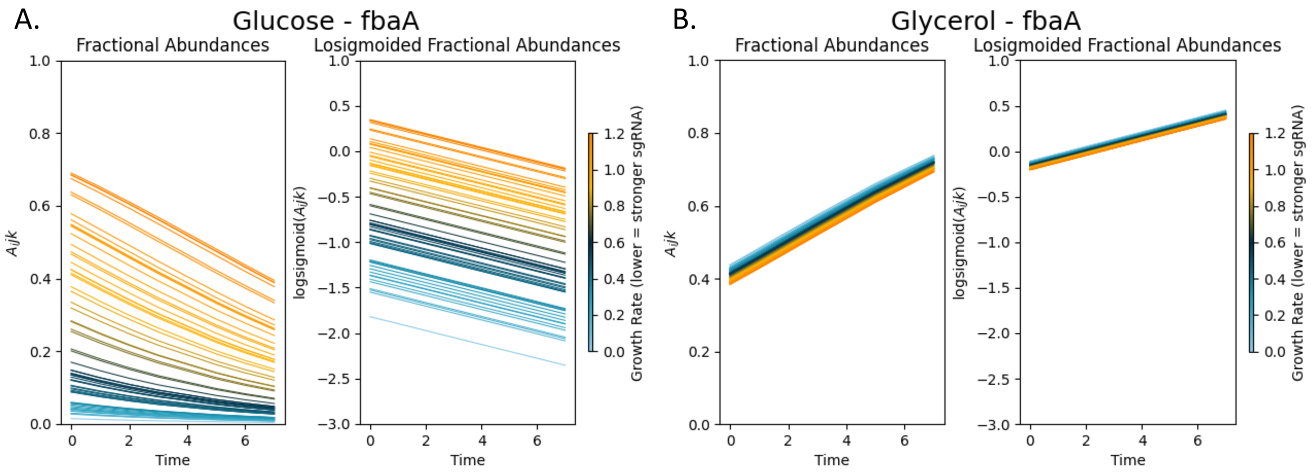


**Fig 9. Time vs. Abundance Curves for *fbaA* for growth in glucose and glycerol.** A) Relative abundances in *fbaA* versus time in glucose shows sigmoidal curves that can be linearized, revealing a strong depletion of *fbaA* for growth in glucose. B) Relative abundances in *fbaA* versus time in glycerol does not shows as obvious sigmoidal curves that can be linearized, revealing an almost enrichment of *fbaA* for growth in glycerol.

**Predicting uninduced Abundances from SCV**

The dataset generated by Mathis, Otto and Reynolds (11) quantified growth rates of the sgRNA mutants but they did not have any measurements equivalent to our uninduced abundances, i.e. abundances of the mutants in the growth mediums without ATC induction. However, the uninduced abundances could be estimated from the induced (no drug) using the SCV (standard coefficient of variation). We observed (on other datasets) that genes with greater depletion due to CRISPR interference had higher noise among their counts (abundance), which could be quantified by SCV. Fig 10 demonstrates the correlation seen of the SCV of the abundances at all concentrations is correlated with the uninduced abundances in the simulated highest noise scenario (HH). The more depleted sgRNAs have higher SCVs since lower number of total counts can increase the amount of noise. We observe similar relationships in the CRISPRi datasets as well.

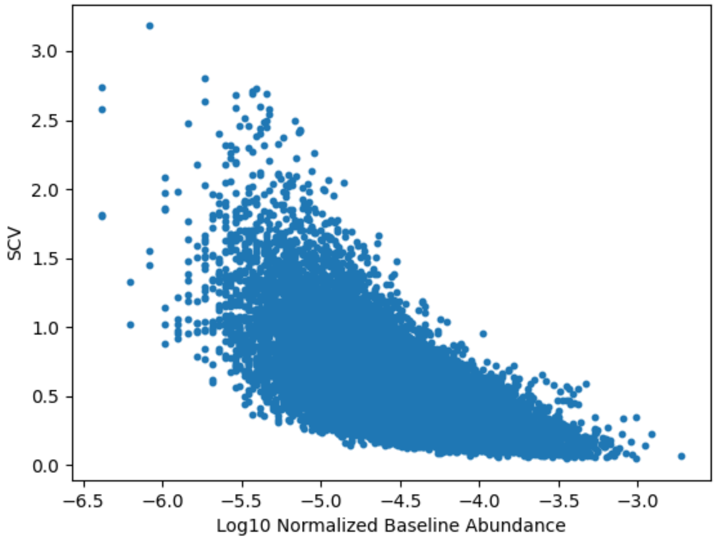


**Fig 10. Correlation of SCV across concentrations vs. uninduced abundances in the simulated HH scenario.** The points are linearly correlated with higher depletion resulting in higher SCV.

The calculation of the uninduced abundances using SCV of the induced counts available is:

$$SCV = \frac{stdev(Y_i)}{mean(Y_i)}$$

$$B_i = \overline{Y_{i0}} \cdot e^{\lambda \cdot SCV}$$

where $B_i$ = baseline abundances calculated for gene $i$, $Y_i$ = counts across all concentrations ($Y_{i0}$ are counts for gene $i$ specifically at 0 concentration) and $\lambda = 2$ for this dataset. Therefore, we used this method to estimate uninduced abundances for the *E. coli* CRISPRi data.

**Results**

We ran CRISPRi-DR independently for each carbon source, using the SCV method to generate uninduced baseline abundances. In both analyses, a significant number of genes exhibited notable depletion (indicating reduced fitness) or had a time-dependence coefficient q value of less than 0.05. This may be because many of the 88 genes selected for this experiment are specifically because they are essential for growth (on either carbon source).

As depicted in Fig 11A, most of the coefficients of time-dependence have a Z score between -1 and 1. While the two curves are similar, they are not identical. The deviation in the distribution curves around a Z-score of 1 can be traced back to two genes labeled in orange in Fig 11B (*fbaA and pfkA*). In Fig 11B, we observe a strong correlation in the coefficients of time dependence of most genes; they align closely with the y=x line. These time parameter coefficients ($\beta_{T_{Glucose}}, \beta_{T_{Glycerol}}$) are analogous to the coefficients of concentration dependence is reflective of the interaction of a gene with the chemical in typical CRISPRi-DR outputs. The glycolytic genes, highlighted fuchsia, are necessary for growth on both carbon sources. These genes are closest to the y=x line, showing similar fitness changes over time in both carbon sources, as expected. As mentioned previously, there are two notable genes (highlighted orange) *fbaA* (fructose bisphosphate aldolase) and *pfkA* (phosphofructokinase). These genes, identified in the Mathis, Otto and Reynolds (11) analysis are well-known examples of genes involved in glycolysis but not for incorporation of glycerol, as aptly have more negative coefficients in the glucose dataset analysis than the glycerol.

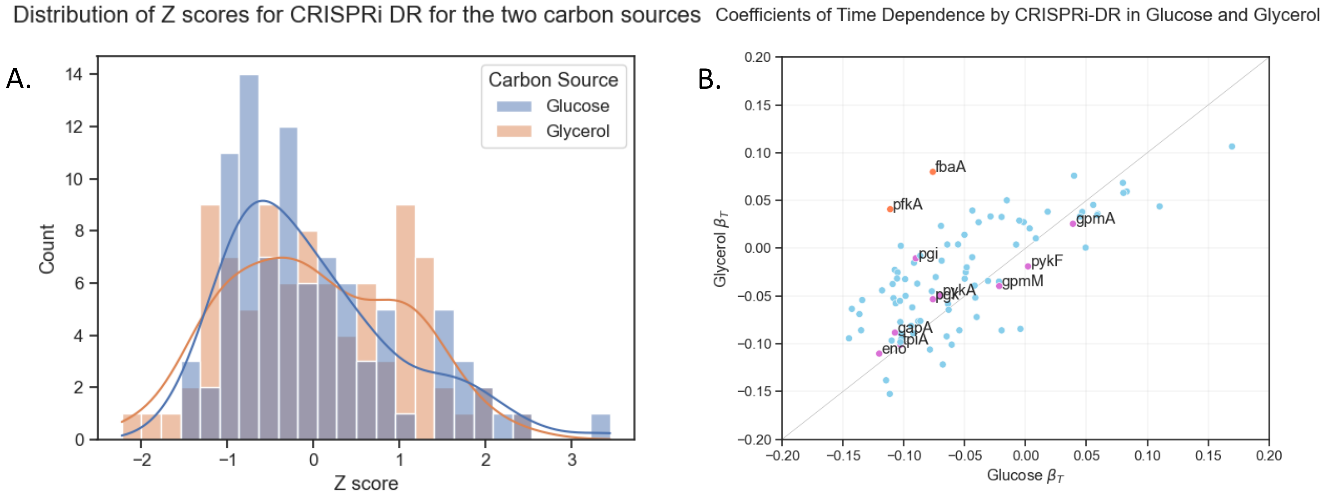


**Fig 11. Coefficients of Time-Dependence in CRISPRi-DR Analyses of Glucose and Glycerol Data.** A) Z-Score Distribution of the Coefficients. The distribution of coefficients B) Correlation plot of the coefficients of the genes. The solid diagonal line is y=x. The fuschia labeled points closest to the line are genes involved in both gluconeogenesis and glycolysis. The points farther away from this line, the orange labeled points (*pfkA* and *fbaA),* are genes involved in glycolysis but not gluconeogenesis and; they have more negative coefficients in glucose than in glycerol.

The analysis of this dataset demonstrates that the CRISPRi-DR method can be applied to other datasets, including those not explicitly designed for chemical-genetics. The modified Dose-Response model nicely incorporates the simultaneous effects of time and the variable efficiencies of sgRNAs of varying effiency on mutant abundance.

### 4. Minimum Number of sgRNAs per gene needed for CRISPRi-DR

Creating a library of sgRNAs can be expensive and time-consuming. A user may want to know how many sgRNAs per gene are necessary to span a range of predicted efficiencies and reflect genuine interactions with a given treatment. Based on our investigation below, we recommend at least 5 sgRNAs per gene.

We subsampled our existing library of about 96,000 total sgRNAs such that each gene has a maximum of 2, 4, 6, 8, 10, and so on sgRNAs per gene. We re-rank the genes after running these subsampled libraries through the CRISPRi-DR model. ROC Curves with target genes obtained from Xu, DeJesus (10) reveals that 2 sgRNAs per gene is not enough to capture expected interactions, but at least 5 sgRNAs spanning a range of predicted efficiencies is sufficient.

Rankings of Select Genes with Reduced Sets of sgRNAs

In a library treated with isoniazid at 1 day pre-depletion, we sampled all the genes to have a maximum of 2-20 sgRNAs, incrementing at intervals of 2, and ran the sampled libraries through the CRISPRi-DR model. We repeated this sampling 10x each at every increment.

In Panel A of Fig 12, we see *inhA* (enoyl-ACP reductase, in mycolic acid pathway) is an essential gene that is the target of INH [12], and *nadA*, an enriched gene in this dataset (with all sgRNAs). The figure depicts changes in ranking of these two genes as the number of sampled sgRNAs is increased. Depletion ranking in this context is defined as genes in the order of increasing concentration dependence slope and enrichment ranking is genes in the order of decreasing concentration dependence slope. With all the sgRNAs, the depletion ranking of *inhA* is #12 and the enrichment ranking of *nadA* is #21. The shaded region surrounding the line is the standard deviation of the rankings across the 10 iterations performed at a particular sampling level. At the left end of the plot, with a low number of sgRNAs sampled per gene, the standard deviations are high, at 30.2 for *inhA* and 134.6 for *nadA*. These standard deviations reduce sustainably and rankings for both genes start to converge to their true rankings (based on all sgRNAs) at around the 5 sgRNA sampling level.

We observed a similar phenomenon in Panel B of Fig 12, with *rpoC* and *eccD3,* a few genes that interact with rifampicin with 1 day pre-depletion. The variation of the rankings of these genes is possibly more variable than those in the isoniazid library depicted in Panel A, with a standard deviation of 283.8 for *eccD3* and 202.6 for *rpoC* at the 2 sgRNA sampling level. However, the rankings of these genes also converge at the 5 sgRNA subsampling level.

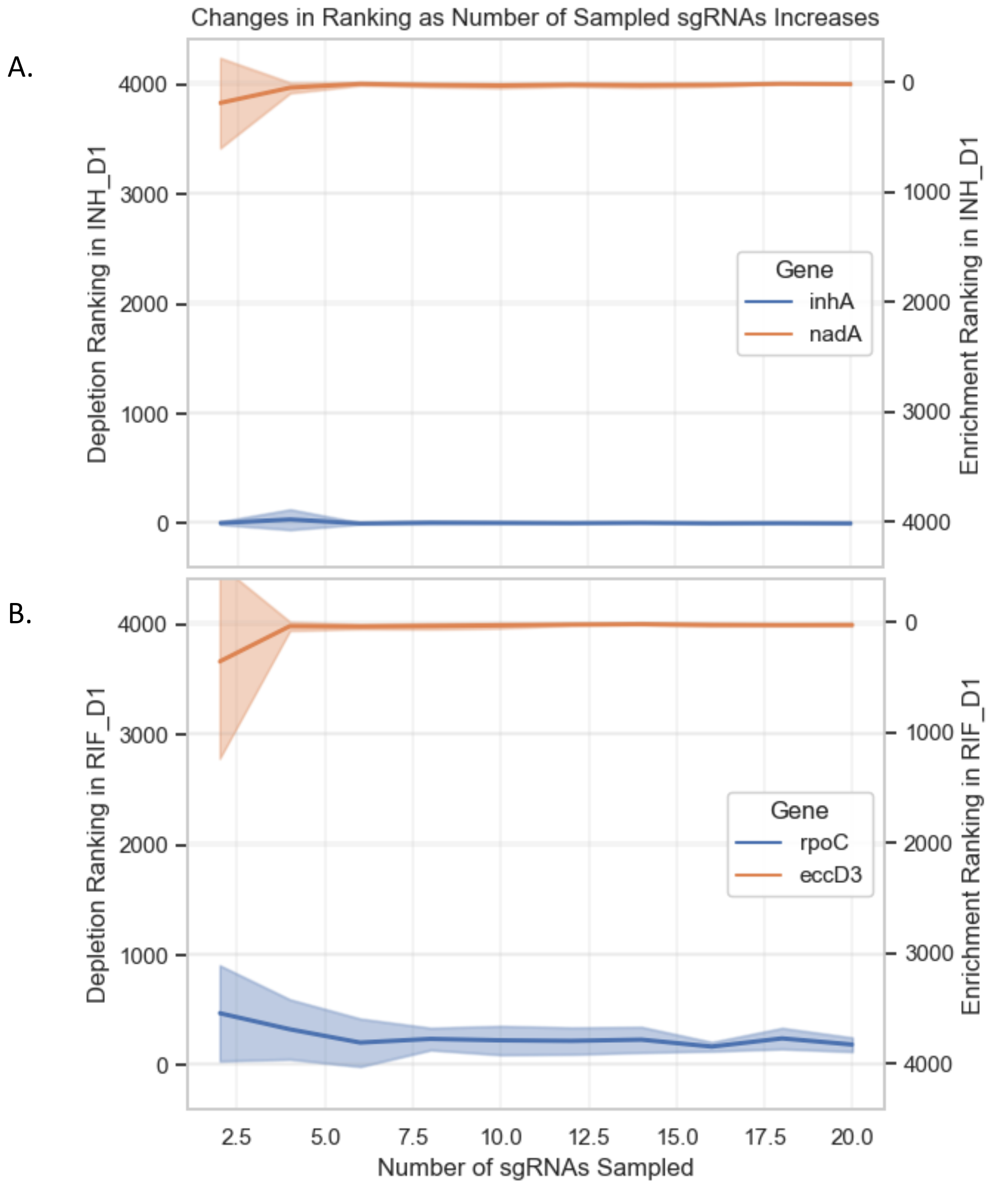


**Fig 12. The rankings for select genes based on maximum number of sgRNAs sampled per gene.** (A) select genes from INH 1 day pre-depletion. *inhA* is significantly depleted and *nadA* is significantly enriched in the presence of isoniazid (B) Select genes from RIF 1 day pre-depletion. *rpoC* is highly depleted and *eccD3* is highly enriched in the presence of rifampicin. Each sampled sgRNA library is run through the CRISPRi-DR model 10 times. The shaded regions represent the standard deviations of rankings over 10 iterations at every sgRNA sampling value. In both panels, gene rankings converge at about 5 sgRNAs.

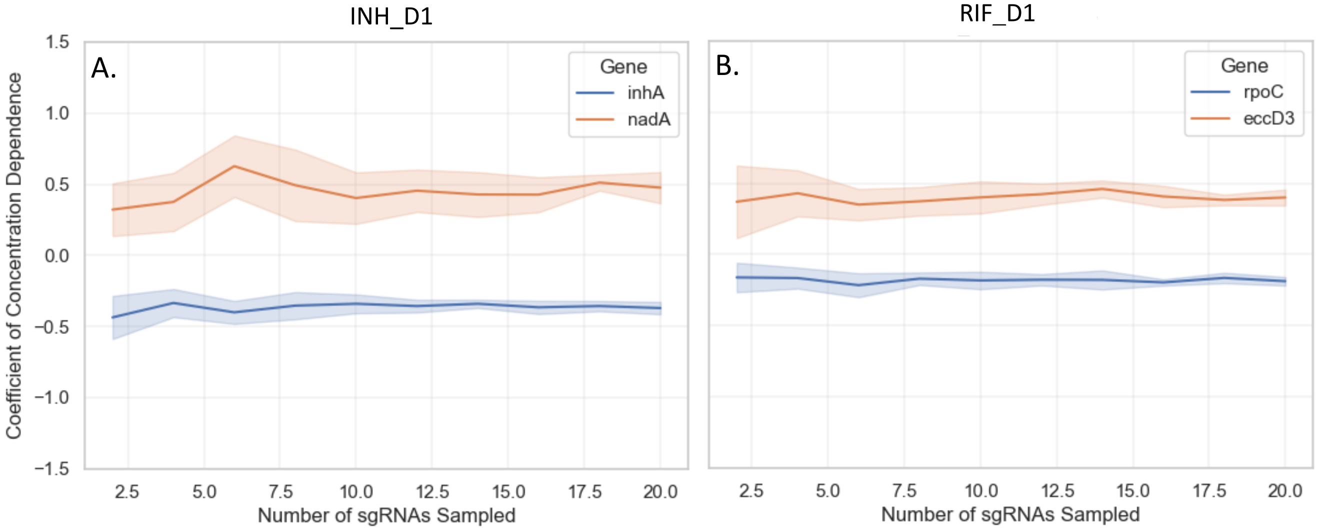


**Fig 13. Coefficient of concentration dependence select genes based on maximum number of sgRNAs sampled per gene.** (A) select genes from INH 1 day pre-depletion. (B) Select genes from RIF 1 day pre-depletion Each sampled sgRNA library is run through the CRISPRi-DR model 10 times and the slopes of concentration dependence extracted for the select genes.

Too few sgRNAs could reduce the stability of the regressions and lead to less accurate (or more variable) estimates of slope coefficients. As seen in Fig 14, the $r^2$ values of the select genes show a high standard deviation when less than 5 sgRNAs are sampled (which means the slope coefficients indicating concentration-dependence vary depending on which sgRNAs are selected see Fig 13), which in turn impedes the model's ability to detect significant interactions.

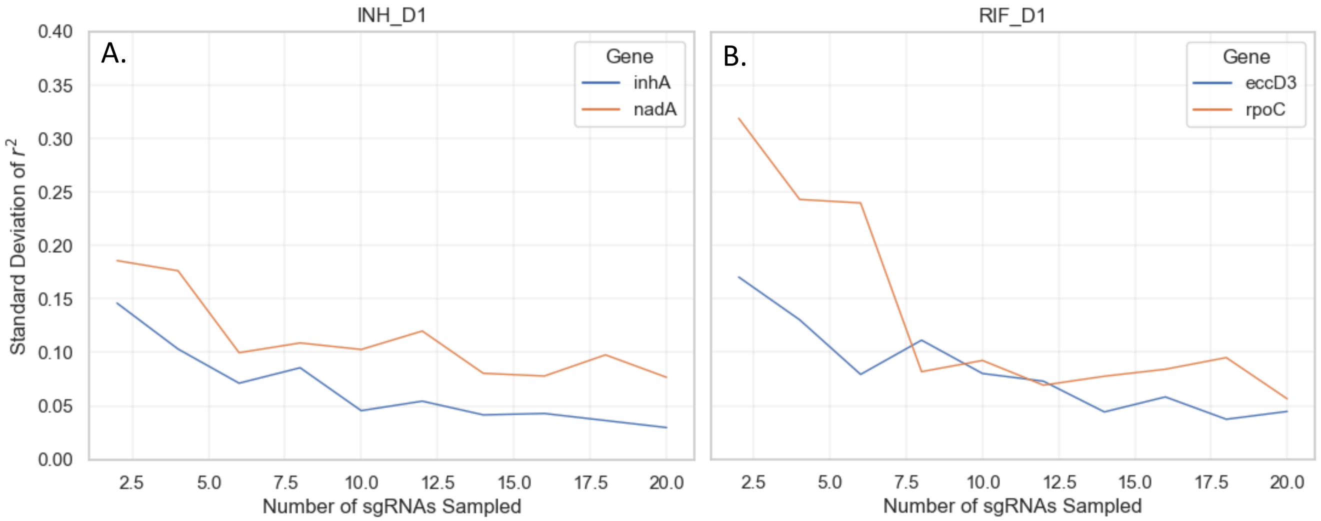


**Fig 14. Standard Deviations of $r^2$ for fitting of select genes based on maximum number of sgRNAs sampled per gene.** (A) select genes from INH 1 day pre-depletion. (B) Select genes from RIF 1 day pre-depletion Each sampled sgRNA library is run through the CRISPRi-DR model 10 times and the $r^2$ values extracted for the select genes. The standard deviations of these $r^2$ values are obtained and for each sampling level and plotted here. In both panels, standard deviation starts to converge at about 5 sgRNAs.

ROC Curves of Subsampled Libraries

Using target genes from Xu, DeJesus (10) for INH and RIF, ROC Curves were generated for each of the sampled sgRNA libraries. As Fig 15 shows, in both RIF and INH, the lowest performing library is that where 2 sgRNAs are sampled for all genes in INH and RIF screens. After that (6 sgRNAs and upward), performance remains consistently high. Combined with the ranking changes observed in Fig 12 and the changes in standard deviation of $r^2$ values observed in Fig 14, this suggests that each target gene should be represented by a minimum of 5 sgRNAs in designing a CRISPRi library.

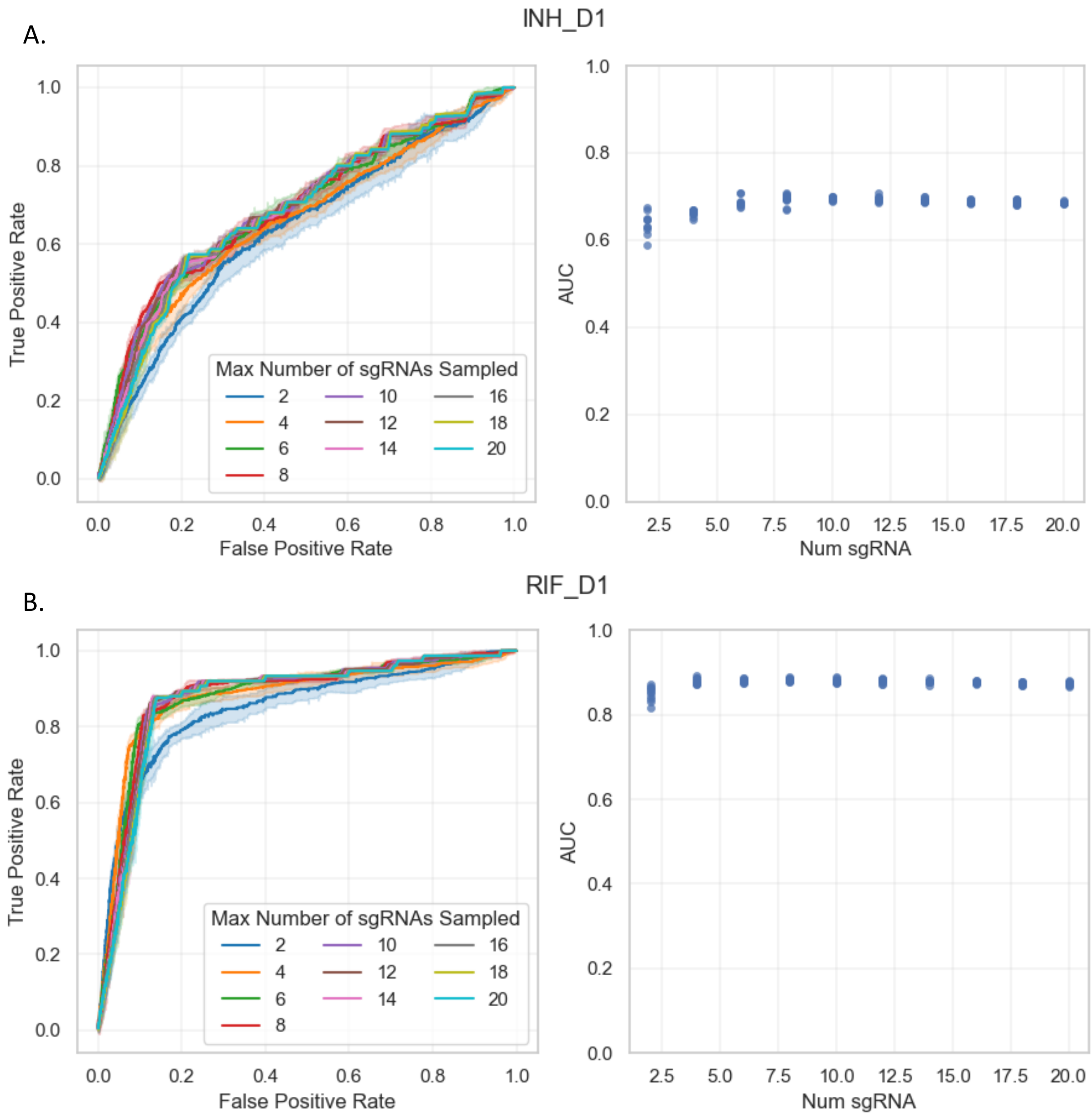


**Fig 15. ROC Curves for each of the maximum number of sgRNAs sampled.** (A) ROC Curves for the different number of maximum sgRNAs sampled from the INH 1 day pre-depletion, along with the AUC values. (B) ROC Curves for the different number of maximum sgRNAs sampled from the RIF 1 day pre-

depletion, along with the AUC values. The shaded regions surrounding the lines is the standard deviation of the true positive rate at each false positive rate. In both Panels, the worst performing sgRNA sampling amount is 2, with the lowest AUC across the 10 runs. Starting at 4 sgRNAs, the performance of the library remains constant.

Thus, we conclude that at least 5 sgRNAs (of diverse efficiency) should be included for each gene when designing a CRISPRi library, to ensure adequate regression fits with CRISPRi-DR.
